# Mitochondrial DNA mutations in human oocytes undergo frequency-dependent selection but do not increase with age

**DOI:** 10.1101/2024.12.09.627454

**Authors:** Barbara Arbeithuber, Kate Anthony, Bonnie Higgins, Peter Oppelt, Omar Shebl, Irene Tiemann-Boege, Francesca Chiaromonte, Thomas Ebner, Kateryna D. Makova

## Abstract

Mitochondria, cellular powerhouses, harbor DNA (mtDNA) inherited from the mothers. MtDNA mutations can cause diseases, yet whether they increase with age in human germline cells—oocytes—remains understudied. Here, using highly accurate duplex sequencing of full-length mtDNA, we detected *de novo* mutations in single oocytes, blood, and saliva in women between 20 and 42 years of age. We found that, with age, mutations increased in blood and saliva but not in oocytes. In oocytes, mutations with high allele frequencies (≥1%) were less prevalent in coding than non-coding regions, whereas mutations with low allele frequencies (<1%) were more uniformly distributed along mtDNA, suggesting frequency-dependent purifying selection. In somatic tissues, mutations caused elevated amino acid changes in protein-coding regions, suggesting positive or destructive selection. Thus, mtDNA in human oocytes is protected against accumulation of mutations having functional consequences and with aging. These findings are particularly timely as humans tend to reproduce later in life.

**Graphical abstract:** 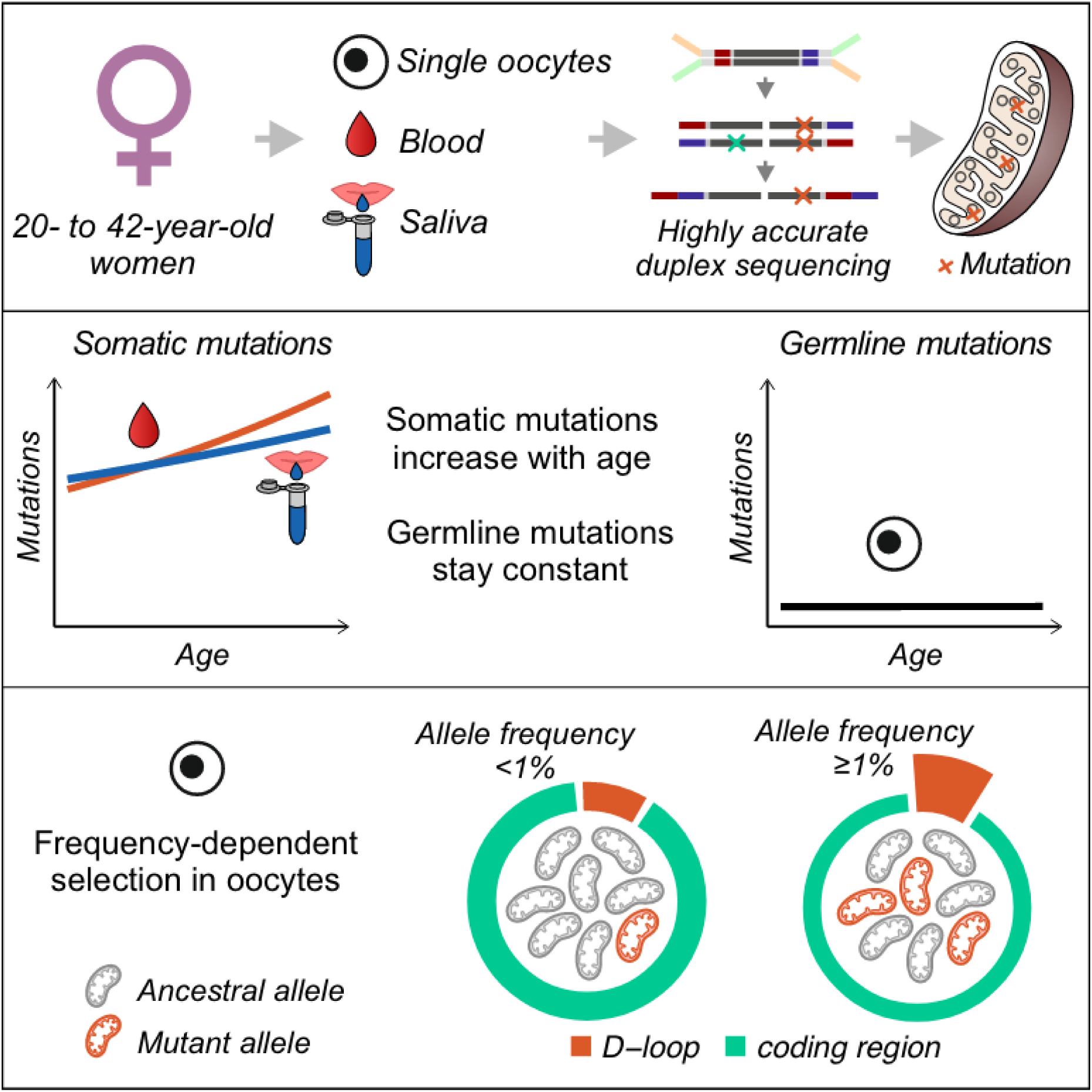

## Introduction

Mitochondria are organelles that have a multitude of functions central to the eukaryotic cell, including the production of ATP via oxidative phosphorylation, response to intrinsic and extrinsic stress by inducing metabolic signaling, etc. ^reviewed^ ^in^ ^1^. Mitochondrial dysfunction manifests itself in diverse and frequently multi-systemic diseases ^1^. How mitochondrial diseases originate is of great interest to the biomedical community.

Mitochondrial dysfunction and subsequent diseases are caused in part by mutations in the mitochondrial genome, i.e., mtDNA ^reviewed^ ^in^ ^2–4^, a 16.6-kb long circular double-stranded DNA with 37 genes encoding 13 proteins, 22 transfer RNAs (tRNAs), and two ribosomal RNAs (rRNAs) ^reviewed^ ^in^ ^5^. Many other mitochondrial genes are located in the nuclear genome ^6^. In contrast to nuclear DNA, with each chromosome usually present in two copies in somatic cells and one copy in germline cells, mtDNA is present in hundreds to thousands of copies in somatic cells (e.g., a median mtDNA copy number of 127 was measured in whole blood ^7^) and >100,000 copies in mature oocytes ^reviewed^ ^in^ ^8^. Human mtDNA is thought to be inherited exclusively through the maternal lineage ^9^.

Examining whether germline mutation frequency increases with age is highly relevant to modern societies in which reproduction later in life has become common. Whereas parental age has been linked to an increase in the frequency of germline mutations *in the nucleus* ^10–15^, the effect of maternal age on germline mutations *in mtDNA* has been debated. Pedigree studies have suggested an increased mtDNA mutation frequency in children born to older mothers ^e.g. 16–18^; however, post-fertilization selection might have affected them. Indeed, without the direct analysis of oocytes, we cannot ascertain whether an increased frequency of *de novo* mutations in the offspring is due to enhanced mutagenesis and selection in oocytes in older mothers vs. due to post-fertilization selection in children born to older mothers.

Whereas mtDNA mutations have been analyzed in human somatic tissues in detail ^e.g. 19^, the direct examination of mtDNA mutations in human oocytes has been challenging due to methodological limitations. Most previous studies either focused on particular mtDNA sites ^e.g. 20,21^ or used sequencing methods with high error rates ^e.g. 22–24^. Using a low-error duplex sequencing approach ^25^, we have recently shown that mutations across the whole mtDNA increase with age in *mouse* oocytes ^26^. Using the same approach, we have demonstrated that mtDNA mutations increase in *macaque* oocytes until ∼9 years of age and do not increase afterward ^27^. Importantly, we still do not know definitively whether the frequency of *de novo* mtDNA mutations increases with age in *human* oocytes.

To address this knowledge gap, we analyzed mtDNA substitution mutations in single oocytes, blood, and saliva from women of different ages. We used the highly accurate duplex sequencing method ^25^, which we had previously modified to generate high-quality mtDNA sequences directly from single oocytes ^26,27^. We obtained a comprehensive set of mutations to study the impact of age on frequencies of germline and somatic mutations, as well as on their distribution across mtDNA. We identified genetic variant hotspots and analyzed the effect of selection on the mutational landscape. In addition, we separately analyzed inheritable heteroplasmies—variants present in at least one oocyte and all somatic tissues assayed, and therefore likely already segregating in the germline during oogenesis. This allowed us to estimate the size of the effective germline bottleneck occurring in the oocyte lineage prior to fertilization and embryogenesis. Thus, we present a direct analysis of mtDNA variants in the female germ line, unraveling their dynamics depending on age and allele frequency.

## Results

### Duplex sequencing and mutation detection

To study the role of age in germline and somatic mtDNA mutation accumulation in humans, we generated high-quality, full-length mtDNA sequences for 22 women (mainly of European ancestry) ranging in age from 20 to 42 years (**Fig. 1A, Table S1**). For every woman, we assayed mtDNA from 1 to 5 single oocytes (mean=3.64; standard deviation, or SD=1.05; a total of 80 single oocytes were assayed), blood (n=18), and/or saliva (n=8; **Table 1, Table S1**).

**Fig. 1.**
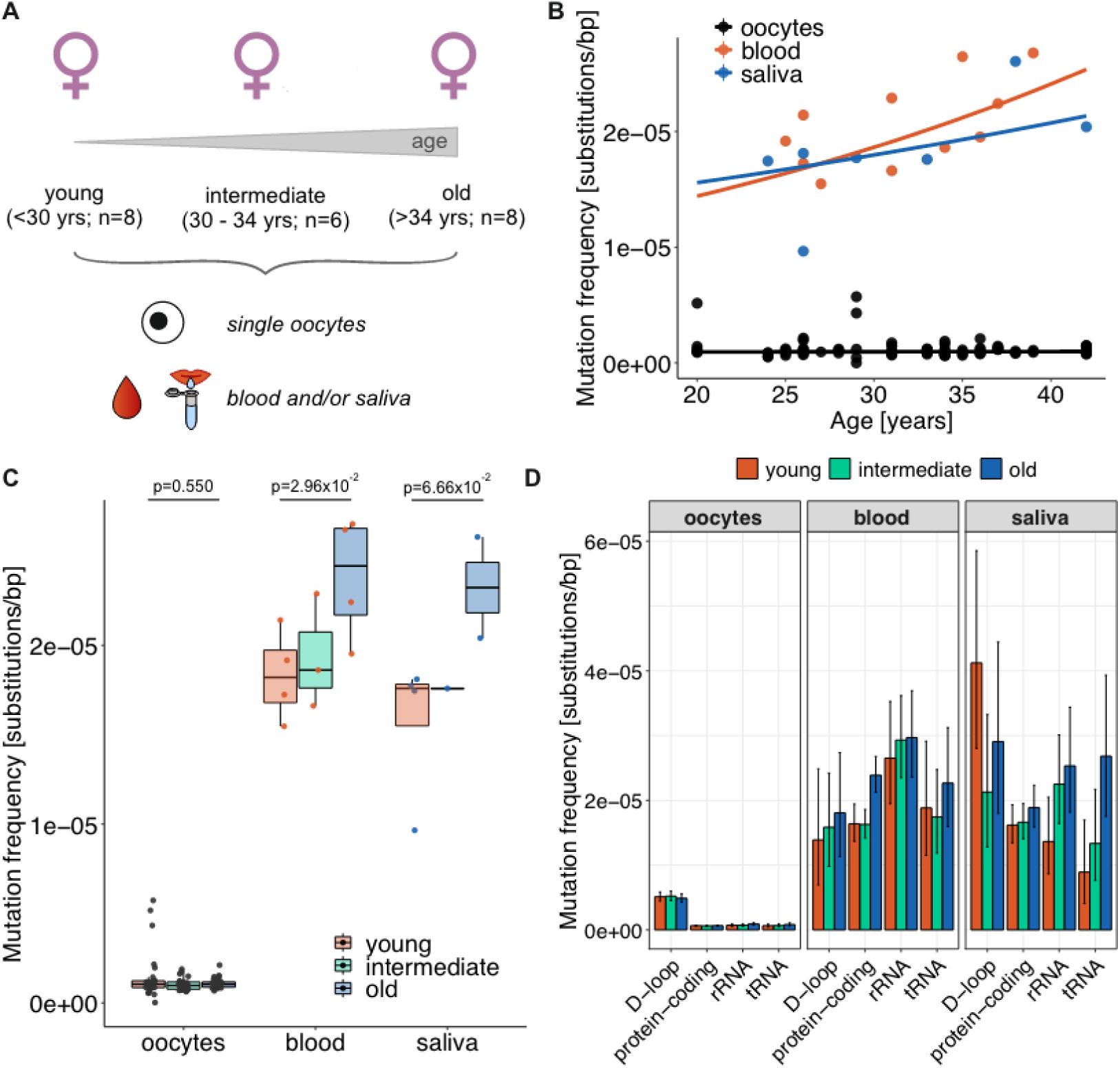
MtDNA mutation frequencies in oocytes, blood, and saliva. **(A)** Oocytes, blood, and saliva samples were sequenced from 20-42 year-old women, assigned to three age groups: young, intermediate, and old. **(B)** Mixed-effects linear model analyzing the effect of age on mutation frequencies. Curves show the predicted mutation frequency based on the fixed-effect part of the model. Dots are the observed mutation frequencies per individual oocyte for oocytes, and per tissue per donor for somatic tissues. **(C)** Mutation frequencies measured in single oocytes, blood, and saliva are shown for each donor and each age group. *P*-values are from a permutation test comparing young and old age groups (one-sided test based on medians; 10,000 permutations; corrected for multiple testing; see Methods). **(D)** Tissue-specific mutation frequencies (i.e., the total number of tissue-specific mutations divided by the product of mtDNA region length and average DCS depth) for each age group in the D-loop (1,122 bp), protein-coding regions (11,341 bp), rRNA (2,513 bp), and tRNA (1,505 bp). Mutation frequencies for non-coding sequences outside of the D-loop (88 bp) are shown in **Table S6**. Mutation frequency bars are shown with 95% Poisson confidence intervals.

**Table 1.** Donors and samples assessed with mtDNA duplex sequencing.

|  | Young<br>(20-29 years) | Intermediate<br>(30-34 years) | Old<br>(35-42 years) |
| --- | --- | --- | --- |
| **Number of donors** | 8 | 6 | 8 |
| **Average age in years (SD)** | 25.6 (2.8) | 32.7 (1.4) | 38.1 (2.7) |
| **Oocytes (DCS depth <50x)\*** | 28 (0) | 25 (2) | 27 (0) |
| **Blood samples (DCS depth <50x)\*** | 6 (2) | 5 (2) | 7 (3) |
| **Saliva samples (DCS depth <50x)\*** | 5 (1) | 1 (0) | 2 (0) |

SD—standard deviation
DCS—duplex sequence consensus
\*The number of samples with DCS<50x is shown in the parentheses.

To identify DNA variants, we used *duplex sequencing* ^25^. By barcoding each double-stranded sequencing template prior to DNA amplification during library preparation, duplex sequencing separates true DNA variants from PCR errors arising during library preparation and from sequencing errors. This is achieved by first forming consensus sequences for reads originating from each of the two template strands separately, i.e., *single-strand consensus sequences (SSCSs)*, which still contain some PCR errors and artifacts resulting from DNA lesions, and next by forming a consensus of the two SSCSs, i.e., *duplex consensus sequence (DCS)*, where the errors and artifacts are eliminated. We followed the protocol we previously developed for enriched mtDNA in mice ^26^. Briefly, the samples were enriched for circular mtDNA with the exonuclease V digestion of linear nuclear DNA (**Table S2, Fig. S1A**). The remaining DNA was sheared, sequenced with short (250-nt) paired-end reads, and analyzed with the Du Novo software ^28,29^ to form DCSs (see **Methods** for details). We achieved a median mtDNA enrichment of 85.4%, 0.7%, and 4.0%, and a median DCS depth of 1,440×, 78×, and 158×, per site per sample for single oocytes, blood, and saliva, respectively (**Table S2, Fig. S1AB**), with a rather uniform DCS depth across the mtDNA length (**Fig. S1C**). The DCSs were mapped to the human reference genome including nuclear DNA and mtDNA, and only alignments with the best match to the mtDNA were retained (see **Methods** for details). At 219 different mtDNA positions, we identified fixed differences from the reference (supported by all DCSs in all tissues studied for an individual), with 8-40 such sites per individual (**Fig. S2**). Additionally, we called *variants*—sites with nucleotides different from those present in most DCSs for a sample (see **Methods**).

Across all samples studied, we identified 4,643 variants. Among them, we detected and analyzed separately (see below) 148 *inheritable heteroplasmies* (i.e., variants present in all somatic tissues assessed and ≥1 oocyte of a donor). From the remaining variants, we filtered out (a) 146 variants with ambiguous timing of occurrence during development (i.e., representing potential inheritable heteroplasmies, early somatic mutations, or early germline mutations; see **Methods**); (b) 539 variants present at the beginning or the end of long homopolymer runs (representing potential read mapping errors); (c) 222 linked variants (with >3 variants present in single paired-end reads mostly occurring in multiple unrelated samples, representing potential regions of nuclear DNA that are homologous to mtDNA, or Numts); and (d) 63 low-confidence variants present in oocytes with a mean DCS depth <50× (this way we excluded variants from 2 oocytes, but were still able to retain variants from 1 to 5 oocytes per woman). As a result, we obtained 3,525 high-confidence tissue-specific *de novo mutations* (**Fig. S3**).

To study the frequencies and patterns of *de novo* mutations and inheritable heteroplasmies depending on age, we assigned the studied women to three age groups of approximately equal size: (1) young (20-29 years; 8 women; 28 single oocytes), (2) intermediate (30-34 years; 6 women; 25 single oocytes), and (3) old (35-42 years; 8 women; 27 single oocytes; **Fig. 1A**, **Table S1**). The distribution of 3,525 *de novo* mutations among the age groups and tissues is shown in **Table S3**. Most *de novo* mutations (3,051) were each measured in a single DCS, and thus had low Minor Allele Frequencies (MAFs). A total of 247 *de novo* mutations (69, 108, and 70 in oocytes, blood, and saliva, respectively) had a MAF ≥1%, a common cutoff for mtDNA mutation detection in other studies ^e.g. 17,18^. A total of 184 of them were supported by single DCSs, and for these mutations high MAFs likely resulted from low sample sequencing depth. To alleviate this potential limitation, we also identified *de novo mutations (*and later also inheritable heteroplasmies*) with high-confidence MAFs* as those supported by ≥3 DCSs (allowing a more reliable MAF estimation). A total of 64 of 247 (59, 3, and 2 in oocytes, blood, and saliva, respectively) *de novo* mutations satisfied this criterion (**Fig. S4**). Because these mutations are tissue-specific, they likely rose to high frequencies due to clonal expansions. However, we cannot exclude the possibility of multiple independent mutations originating at the same position in the same sample.

### *De novo* mutations

#### Increase in mutation frequency with age in somatic tissues but not in oocytes

Among all age groups and regions analyzed, oocytes displayed a ∼17-24-fold lower mutation frequency than blood and saliva (**Fig. 1B-D**). To study the effect of age on mutation frequency, we built a generalized mixed-effects linear model considering each individual and each tissue separately (each oocyte was also considered separately; see **Methods**), following our approach in ^27^. The model predicted the probability of having a mutation at a site as a function of age (**Fig. 1B**) and included the mutation frequency as a response, the number of sequenced nucleotides per sample (i.e., sequencing depth per site) as weight, and age, tissue, and their interactions as fixed-effect predictors. The donor ID was used as a random effect. The model indicated a significant increase in mutation frequency with age in blood, a marginally significant increase in saliva, and no difference in oocytes (Z-test *p*=2.7×10^-3^, *p*=5.7×10^-2^, and *p*=0.688 for blood, saliva, and oocytes, respectively; **Fig. 1B, Table S4**). The donor ID had a marginally significant effect (Likelihood Ratio test *p*=0.063), indicating potential differences in mtDNA mutability among different women.

We then compared the mutation frequencies among women from different age groups (i.e., young, intermediate, and old). In agreement with results from the generalized mixed-effects linear model, in blood, the mutation frequency was significantly higher in old compared to young women (i.e., between the two extreme age groups; one-sided permutation test for equality in medians *p*=2.96×10^-2^), with a 1.34-fold increase. In saliva, the mutation frequency was marginally significantly higher in old compared to young women (one-sided permutation test for equality in medians *p*=6.66×10^-2^), with a 1.32-fold increase. In oocytes, no significant difference in mutation frequency was observed (one-sided permutation test for equality in medians *p*=0.550), and the mutation frequencies for old vs. young women were very similar (0.99-fold decrease; **Fig. 1C, Table S3**). No significant differences were observed in any two adjacent age groups for any tissue. Similar results were obtained when aggregating mutations across women in each tissue and age group (**Table S5**). Such aggregated mutations were used in subsequent analyses of *de novo* mutations to increase statistical power.

#### Mutation types and strand bias

Among all tissues and age groups, most *de novo* mutations were transitions, with transition/transversion ratios of 18.3, 10.3, and 1.7 in blood, saliva, and oocytes, respectively (**Table S6**). The only significant accumulation of a specific mutation type with aging was observed for A>G/T>C transitions in blood (a previously identified mtDNA-specific mutational signature of aging ^30^), with a 1.6-fold increase in old vs. young women (Fisher’s exact test *p*=4.26×10^-4^; **Table S7**). The C>T/G>A mutation frequency was 1.6-to 2.8-fold higher at CpG sites than at non-CpG sites in blood and saliva (significant for both tissues and all three age groups except for saliva of old women), but was not significantly different between CpG and non-CpG sites in oocytes (see **Table S8** for *p*-values).

Consistent with previous reports ^19,27,31^, several mtDNA mutation types in our data exhibited strand bias, defined as a bias in the occurrence of mutations between the light (L) and heavy (H) mtDNA strands. With the L-strand as a reference, we observed a bias of G>A over C>T (4.6- to 39.6-fold), T>C over A>G (2.0- to 3.2-fold), and C>G over G>C (2.3- to 14.3-fold) mutations (**Fig. S5AB**, **Table S9**). When analyzing the D-loop separately, strand bias was evident for transitions in oocytes but not in somatic tissues (**Fig. S5CD**).

#### Mutation frequency and germline mutation rate along the mtDNA

In oocytes, the mutation frequency was significantly higher (6.9- to 8.2-fold) in the D-loop than in the ‘coding region’ (i.e., protein-coding, rRNA, and tRNA regions combined) for each age group studied (**Fig. 1D, Fig. S6**; see **Table S5** for Fisher’s Exact test *p*-values). Using these data and considering only *de novo* mutations with MAF≥1% (n=52; only oocytes sequenced at a mean DCS depth ≥500× were included ensuring that mutations can be measured in ≥3 DCS) and 70 mother-to-oocyte transmissions, we computed the germline mutation rate to be 3.95×10^-4^ (95% CI 2.68-5.60×10^-4^) and 1.95×10^-5^ (95% CI 1.21-2.99×10^-5^) mutations per site per generation for the D-loop and coding region, respectively. Thus, the germline mutation rate was 20.2-fold higher in the D-loop than in the coding region. These results are consistent with elevated mutation rates in the D-loop ^16^. In somatic tissues, a significant difference (a 2.7-fold increase) in mutation frequency between the D-loop and coding region was observed only in saliva of young women (**Fig. S7**, **Table S5**).

The same trend was confirmed when analyzing observed vs. expected numbers of *de novo* mutations based on the length of each mtDNA functional region (**Table S10**). In oocytes, we found the D-loop to harbor a significantly higher number of mutations than expected by chance in each age group analyzed (two-sided binomial test *p*=3.16×10^-96^, *p*=1.83×10^-98^, and *p*=1.02×10^-97^ for young, intermediate, and old, respectively), and protein-coding region to harbor a significantly lower number of mutations (two-sided binomial test *p*=1.20×10^-29^, *p*=6.63×10^-32^, and *p*=1.66×10^-38^ for young, intermediate, and old, respectively; **Table S10**). Similar trends were observed for somatic tissues but were significant only for a few combinations of age groups and functional regions (**Table S10**).

For *de novo* mutations located at protein-coding regions, we measured the nonsynonymous-to-synonymous rate ratios for *de novo* mutations, i.e., hN/hS ^32^. In oocytes, such ratios were within the range of neutral expectations (0.90, 1.19, and 0.99 for young, intermediate, and old women, respectively, **Table S11**). In somatic tissues, the hN/hS ratios were higher than expected under neutrality in young women (1.64 in blood and 2.01 in saliva) and in saliva of women from the intermediate age group (2.54), suggesting positive selection; these ratios were not significantly different from neutral expectations in old women (**Table S11**).

#### Disease-associated variants

We next assessed whether any *de novo* mutations were located at sites previously associated with diseases. A total of 109 confirmed disease-associated positions were considered as listed in MITOMAP ^33^ (see **Methods** for details). We identified a total of 32 disease-associated variants (0.91% of all mutations) located at 29 different positions (with disease-associated positions representing only 0.66% of the mtDNA; **Fig. S8**). Overall, the proportion of positions affected by mutations was higher for disease-associated vs. non-disease-associated sites (26.6%, or 29 out of 109, vs. 16.3%, or 2,685 out of 16,460; Fisher’s Exact test *p*=2.61×10^-2^), suggesting high mutation rate and/or positive/destructive selection acting on the former sites. However, lower proportions of disease-associated mutations among all mutations were measured in oocytes than in blood or saliva (0.75%, 1.00%, and 1.24% of all mutations, or 14, 10, and 8 disease-associated mutations in oocytes, blood, and saliva, respectively), suggesting selection against such mutations in oocytes. Two disease-associated mutations found in oocytes were measured at MAFs ≥1% and in ≥3 DCS: m.3946G>A associated with MELAS syndrome (myopathy, encephalopathy, lactic acidosis, and stroke-like episodes) ^34^ and m.1555A>G associated with DEAF (autism spectrum intellectual disability) ^35^. Three additional mutations in somatic tissues had MAFs ≥1%. However, they were only measured in a single DCS each, rendering their MAF estimation unreliable.

#### Variant hotspots

To identify variant hotspots, we built a probabilistic model that considers tissue-specific estimates of the average mutation frequencies, the high mutability in the D-loop, and the mean DCS depth in each sample ^27^. Based on the model results (**Fig. 2A**), we defined variant hotspots as sites mutated in somatic tissues of at least three women or in oocytes of at least four women. We detected a total of 259 *de novo* mutations at 73 hotspots: 51 mutations at 22 hotspots in blood, 19 mutations at 15 hotspots in saliva, and 189 mutations at 36 hotspots in oocytes (the number of tissue-specific hotspots depends on the total number of mutations measured in each tissue and thus is higher in oocytes). Fifteen hotspot sites overlapped between two different tissues (but never among all three), resulting in a total of 58 affected mtDNA sites (**Table S12**). Overall, as many as 3%, 5%, and 10% of all *de novo* variants in saliva, blood, and oocytes, respectively, were found at hotspot sites occupying only ∼0.4% of the mtDNA length (58/16,569).

**Fig. 2.**
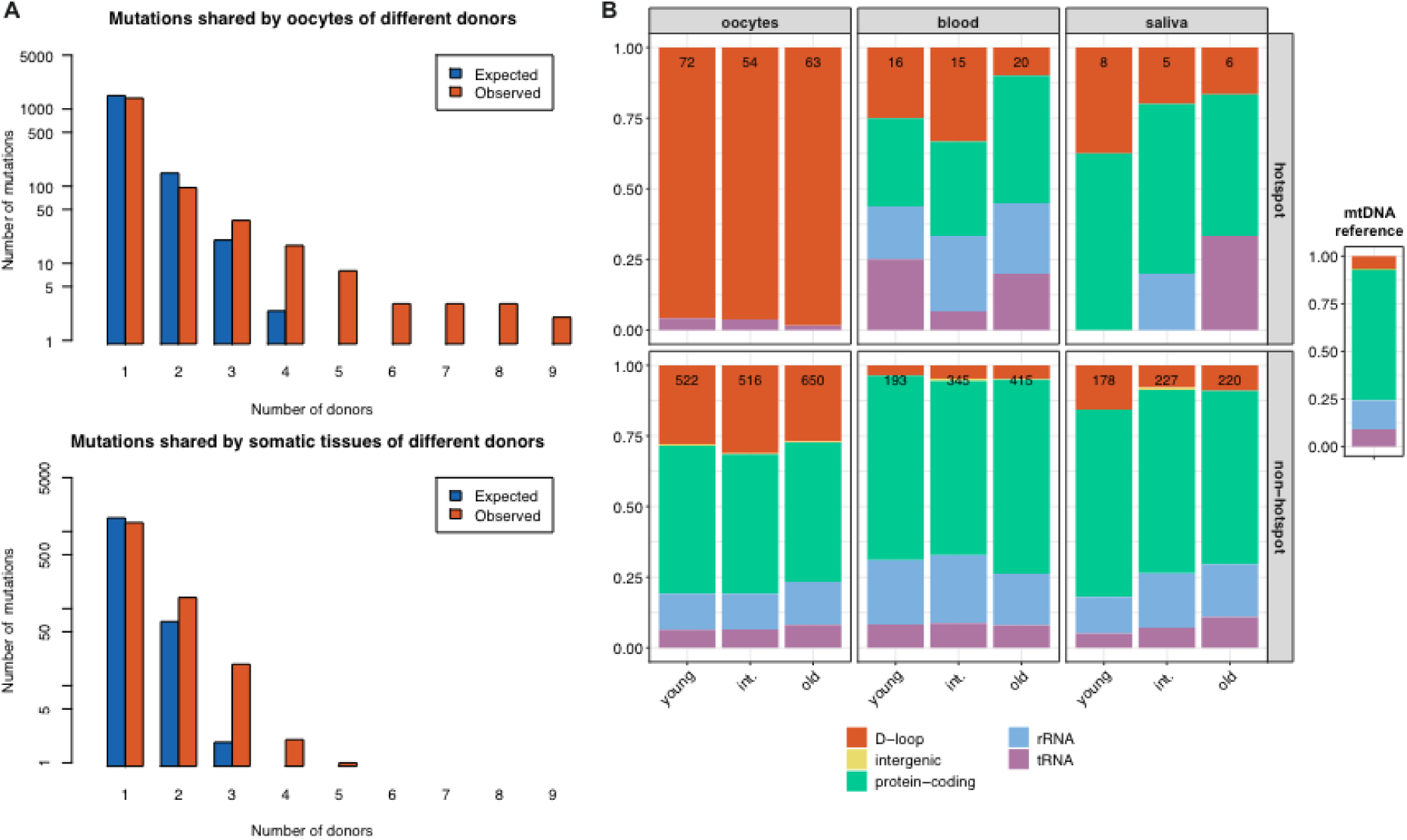
Analysis of variant hotspots. **(A)** The observed and expected distributions of the number of women sharing individual variants in oocytes (with all oocytes from a donor considered together) and somatic tissues. Due to similar mutation frequencies and lower numbers of measured mutations, blood and saliva were considered together as somatic tissues. We also considered the number of sequenced oocytes per donor to evaluate oocyte variant hotspots. For oocytes and somatic tissues separately, we computed the expected number of mutations present in exactly one woman and shared by several women. Our results suggest that, due to random chance, mutations at the same site can occur in two women, but are rarely expected to occur in somatic tissues of three or more women, and in oocytes of four or more women. **(B)** Distribution of mutations (normalized by the length of the respective region) among D-loop, intergenic regions (i.e. non-coding DNA outside of the D-loop), protein-coding regions, rRNA, and tRNA, shown separately for each tissue and hotspots vs. non-hotspots. Numbers indicate the total number of mutations analyzed. The proportion of mtDNA occupied by each region is shown on the right.

Among the 58 hotspot sites, 40 were located in the D-loop, 4 in rRNA, 4 in tRNA, and 10 in protein-coding regions—a distribution that is different from what is expected based on the length of mtDNA regions as well as on the distribution of mutations occurring at non-hotspot sites (**Fig. 2B**). In particular, hotspots were significantly over-represented in the D-loop, consistent with its high mutation rate ^16^, and significantly under-represented in the protein-coding regions (**Table S10D**). In oocytes, 35 out of the 36 hotspots were located in the D-loop. In contrast, in somatic tissues, only 5 out of 22 hotspots were located in the D-loop, but 10 were in protein-coding regions (all nonsynonymous), 4 in rRNA, and 3 in tRNA. These results corroborate our findings of high *de novo* mutation frequency in the D-loop, independent of age, specifically for oocytes (**Fig.1D**, **Table S5**). Only one variant hotspot (somatic) overlapped with a disease-associated mutation: m.3243A>G, a mutation in tRNA associated with MELAS syndrome, one of the most mutable positions in the mtDNA genome ^16^ suggested to be subject to positive selection ^36^.

### Inheritable heteroplasmies

In addition to *de novo* mutations, we investigated heteroplasmies present in somatic tissues and at least one oocyte of the same woman (**Table S13**), and thus were likely already inherited from her mother. Because of their usually higher MAFs, such heteroplasmies have a higher potential to be inherited by the offspring; we called them ‘inheritable heteroplasmies’. To boost our statistical power and confidence in correctly classifying them, in addition to the samples used to study *de novo* mutations, here we also considered samples with average DCS depth below 50×. We also used SSCSs and additional targeted duplex sequencing (see **Methods**) for blood and saliva to aid us in distinguishing between early germline mutations and inheritable heteroplasmies. MAFs measured using exonuclease V digestion and targeted mtDNA capture enrichment were highly correlated (Pearson *r*=0.995, *p*=1.61×10^-12^; **Fig. S9**).

In our data set, 19 women harbored 63 sites (1-9 sites per woman, **Fig. S10**) with inheritable heteroplasmies (13, 22, and 28 sites in young, intermediate, and old women, respectively; **Tables S1** and **S13**), which were all transitions. A total of 45 sites were located in the D-loop, 13 in protein-coding regions, and 5 in rRNA. A total of 14 sites overlapped variant hotspots in oocytes, and three overlapped variant hotspots in somatic tissues; all of them were located in the D-loop. Only one inheritable heteroplasmy was associated with a disease (m.10197G>A associated with Leigh Disease, Dystonia, Stroke, and Leber Optic Atrophy and Dystonia ^33^).

MAFs of inheritable heteroplasmies were correlated between oocytes and somatic tissues (Pearson *r*=0.802, *p*=2.66×10^-15^; **Fig. S11A**), as well as between blood and saliva (Pearson *r*=0.802, *p*=6.30×10^-5^; **Fig. S11B**). Similar results were obtained when including only heteroplasmies with high-confidence MAFs (i.e., those measured in at least three mtDNA copies; 30 heteroplasmic sites; **Fig. S12AB**).

### Genetic drift does not depend on age but depends on MAF

We next tested whether age had an effect on genetic drift, i.e. whether the normalized variance in MAF (variance divided by *P*(1-*P*), where *P* is the average MAF of the allele among single oocytes or across somatic tissues ^37,38^) increases with age. We did not observe a significant correlation between the normalized variance in MAF within each sample type and age (*p*=0.256 and *p*=0.907 for somatic tissues and oocytes, respectively; **Fig. S11CD**). Results were similar when including only heteroplasmies with high-confidence MAFs (**Fig. S12CD**).

To estimate the size of the effective germline bottleneck—the size required to explain observed genetic drift—for human mtDNA, we applied the population genetics approach described in ^38,39^ to the data on MAF shifts at 63 inheritable heteroplasmies measured in 19 women. Namely, we compared allele frequencies between somatic tissues (averaged among the tissues sequenced per donor for donors with more than one tissue sequenced) and single oocytes (with 2-5 oocytes per donor; donor hs001 with 1 oocyte did not have an inheritable heteroplasmy) for each site separately (245 transmissions to oocytes in total; **Table S12**). The effective bottleneck size was estimated to be 909.39 segregating mtDNA units (95% bootstrap CI: 0.47-5,329.73). The effective germline bottleneck computed based on heteroplasmies with high-confidence MAFs (30 heteroplasmies, 111 transmissions) was similar in magnitude: 1,040.03 segregating mtDNA units (95% bootstrap CI: 0.34-11,089.05). Large confidence intervals reflect the large variation in bottleneck size calculated for heteroplasmies measured at low MAFs.

To evaluate whether heteroplasmy MAF affects our estimates of the bottleneck size, we also separately analyzed high-frequency heteroplasmies, i.e. the ones in which at least one variant had MAF≥1%. In total, 27 out of 63 heteroplasmic sites (belonging to 17 women) satisfied this criterion. The effective bottleneck size estimated for such sites based on 100 transmissions to oocytes was 30.41 segregating mtDNA units (95% bootstrap CI: 0.02-178.53)—which was in line with the recent estimates ^e.g. summarized^ ^in^ ^16^. Thus, the effective bottleneck size estimates depend on MAF of heteroplasmic sites. Considering each age group separately, we estimated the effective bottleneck size to be 25.03 units (95% bootstrap CI: 0.47-98.42) for young women, 38.60 units (95% bootstrap CI: 0.02-74.42) for women in the intermediate age group, and 28.38 units (95% bootstrap CI: 0.38-178.53) for old women (**Table S13**). Thus, the bottleneck size was not significantly different among women in different age groups.

### Selection affects allele frequencies of inheritable heteroplasmies

Next, we analyzed whether selection influences changes in MAFs for inheritable heteroplasmies. We referred to the minor allele recorded in the somatic tissues as the minor allele of a heteroplasmic site in a donor, and calculated the MAF difference between oocytes and somatic tissues. A positive (negative) change represents a higher (lower) MAF in oocytes compared to somatic tissues. We observed a significantly lower number of heteroplasmic variants with a positive (n=56) vs. negative (n=189) change (*p*=4.75×10^-18^, binomial test; **Fig. 3AB**). The same remained true when considering only heteroplasmies in protein-coding regions (9 with a positive, and 41 with a negative change, *p*=2.16×10^-5^), in the D-loop (44 with a positive, and 130 with a negative change, *p*=6.53×10^-10^), and in rRNA (3 with a positive, and 18 with a negative change, *p*=3.48×10^-3^). Among heteroplasmies in protein-coding regions, a significantly higher number of negative shifts was observed for nonsynonymous (*p*=2.16×10^-5^), but not for synonymous variants (*p*=0.418; **Fig. 3B**). The same was true when including only heteroplasmies with high-confidence MAFs (**Fig. S13**). These results suggest purifying selection against heteroplasmies transmitted to oocytes and/or positive selection in somatic tissues.

**Fig. 3.**
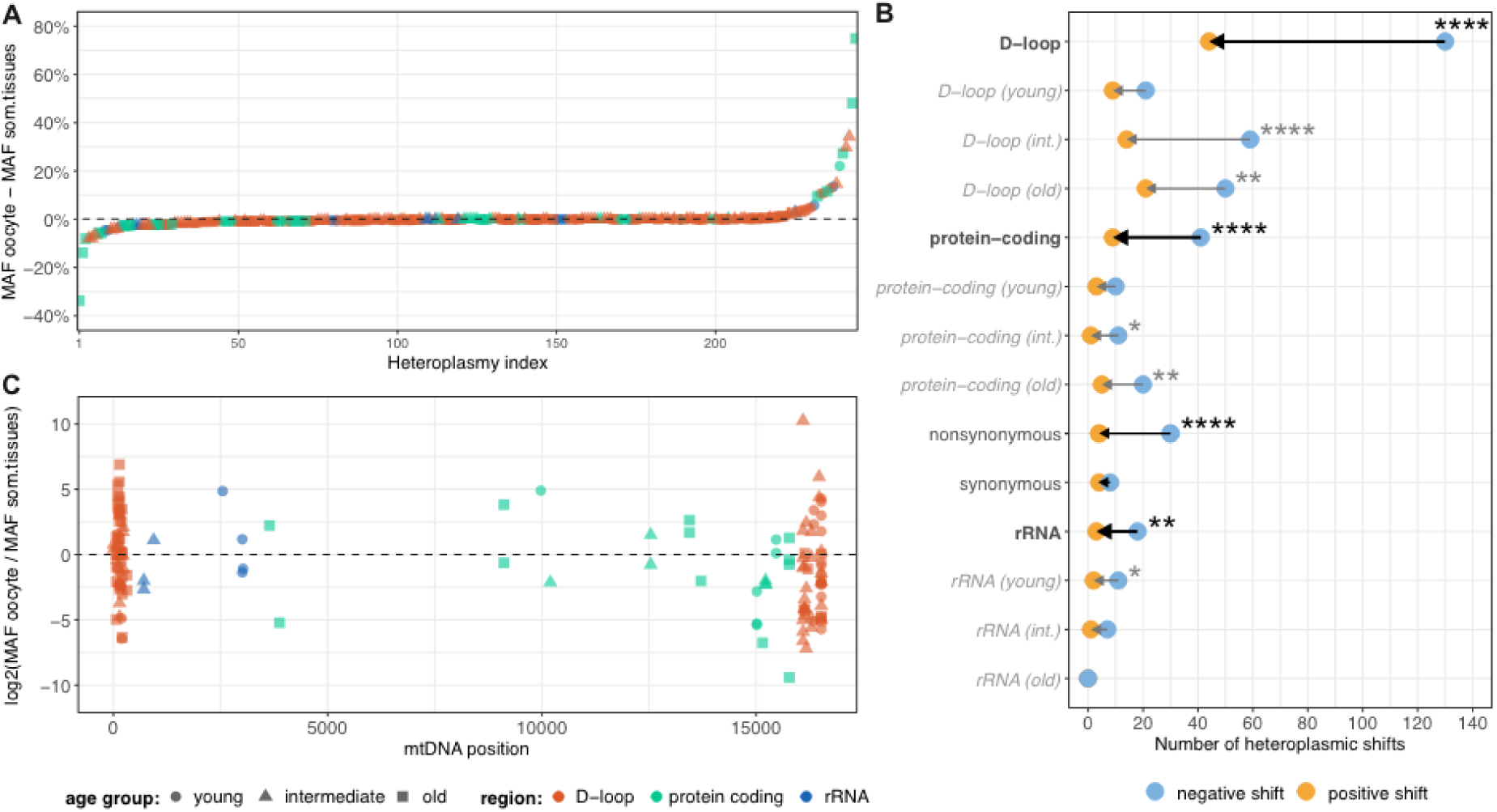
Evaluation of selection in heteroplasmic transmissions. (**A**) MAF difference between oocytes and somatic tissues measured in percentage points; a positive (negative) shift represents a higher (lower) MAF of a heteroplasmy in oocytes compared to somatic tissues for a given a woman. (**B**) Number of heteroplasmies showing an increased or decreased MAF in oocytes (positive or negative shift, respectively), shown separately for mtDNA regions, and either combined or separated by age group. Left-pointing arrows indicate higher numbers of negative versus positive shifts. \**p*<0.05, \*\**p*<0.01, \*\*\**p*< 0.001, and \*\*\*\**p*<0.0001; binomial test, corrected for multiple testing using the method of Benjamini-Hochberg to control the false discovery rate. (**C**) Log2 ratio of MAF difference between oocytes and somatic tissues, calculated to take into account the magnitude of the fold-change in MAFs ^40^. 109 transmissions with MAF=0 in an oocyte are not displayed due to the resulting negative infinite values. Colors indicate their localization in different mtDNA regions, and shapes show the age group of a woman.

Heteroplasmic shifts calculated as log2 fold-change in MAFs ^40^ reached levels around ±10. This represents a ∼1,000-fold change in MAF from a woman to her oocyte (**Fig. 3C**), consistent with a ∼1,000-fold reduction in mtDNA content during human germ cell development ^41^. However, most heteroplasmies had a much smaller change in MAF from a woman to her oocyte, with a <64-fold change in 94.1% of the heteroplasmic variants. Because we used the minor allele recorded in the somatic tissues as the minor allele of a heteroplasmic site in a donor, our results suggest again that purifying selection acts on inheritable heteroplasmies in oocytes, or alternatively positive selection acts in somatic tissues (or a combination of both).

#### Frequency-dependent selection on mtDNA variants in oocytes

We next analyzed the distribution of *oocyte* genetic variants across the mtDNA molecule depending on their frequency and inferred heritability by considering three non-overlapping categories (**Fig. 4, Table S15**): (1) *de novo* mutations with MAF <1%; (2) *de novo* mutations with MAF ≥1% (mean MAF=3.07%, SD=3.96%); and (3) inheritable heteroplasmies (mean MAF=2.07%, SD=8.90%). We found that the majority of variants in the 1st category were located in the coding region (protein-coding, rRNA, and tRNA regions); variants in the 2nd category were distributed approximately equally between the D-loop and the coding region (with slightly more mutations in the D-loop); and variants in the 3rd category were predominantly located in the D-loop. The distribution of mutations inside vs. outside the D-loop was significantly different between *de novo* mutations with MAF<1% and those with MAF ≥1% (*p*=2.04×10^-3^, Chi-squared test), as well as between *de novo* mutations with MAF<1% and inheritable heteroplasmies (*p*=1.75×10^-8^, Chi-squared test), but not between *de novo* mutations with MAF ≥1% and inheritable heteroplasmies (*p*=0.111, Chi-squared test). This pattern was shared by the three age groups considered and suggests selection operating against coding mutations for variants with high frequency in oocytes, and for inheritable heteroplasmies.

**Figure 4.**
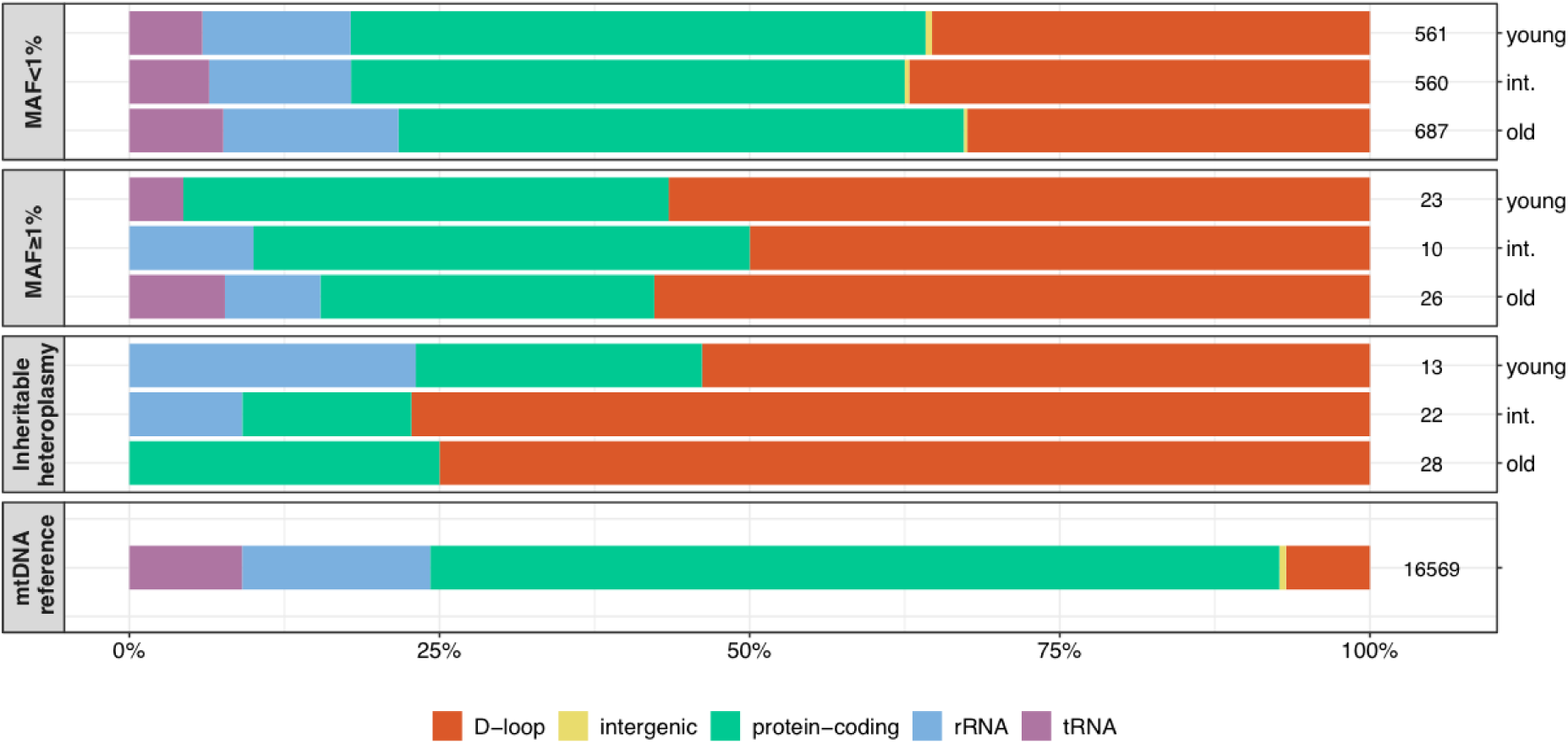
Proportions of mtDNA variants in oocytes depending on their frequency, heritability, and age group. Distribution of mutations (normalized by the length of the respective region) among D-loop, intergenic DNA, protein-coding regions, rRNA, and tRNA, shown separately for *de novo* mutations with MAF <1%, *de novo* mutations with high-confidence MAF≥1% (measured in ≥3 DCS to ensure proper representation on MAFs; 10 mutations measured in <3 DCS were excluded), and inheritable heteroplasmies. Numbers on the left represent the total number of mutations analyzed, or the number of base pairs annotated in each region for the mtDNA reference.

## Discussion

In our study of mtDNA mutations we utilized both germline (single oocytes) and somatic tissues from the same women. This provided us with several unique opportunities. Directly measuring mutations in single oocytes allowed us to disentangle mutations and selection in oocytes from post-fertilization selection. And sequencing at least one somatic tissue in addition to single oocytes allowed us to discriminate between *de novo* somatic and germline mutations, as well as between *de novo* germline mutations and mutations already segregating in the germline.

Additionally, using duplex sequencing, which has an inherently low error rate (<10^-7^) ^19^, allowed us to comprehensively study mtDNA variants with various allele frequencies, including those with MAF<1%, the latter constitute 96.9% of all *de novo* mutations in oocytes in our dataset. This provided a detailed picture of the mtDNA mutational frequency and landscape in oocytes. Due to methodological limitations, previous studies ^e.g. summarized^ ^in^ ^16^ considered only mtDNA variants with frequencies ≥1% (or even higher). This rather high MAF cutoff is particularly problematic for oocytes. Indeed, due to the very high copy number in human oocytes (∼100,000 on average) ^42^, even for a variant observed at a MAF of 1%, ∼1,000 mtDNA molecules would be affected in a single cell. Therefore, very low frequency variants can have considerable impact on mitochondrial function and be substantially affected by selection—yet they were excluded from most previous studies.

### Age does not increase mtDNA mutations in human oocytes

Analyzing women ranging in age from 20 to 42 years, both based on individual age and on age groups, we did not observe an increase in mutation frequency in the oocytes in older women. This differs from what we observed using the same methods (i.e. duplex sequencing) in mouse oocytes, where we found ∼2-fold higher mutation frequencies in 10-month-old compared to ∼1-month-old mice ^26^. However, for rhesus macaques, again using the same methods, we found that mutation frequencies in oocytes increased only until the age of ∼9 years and not further ^27^ (as rhesus macaques age approximately three times faster than humans, ∼9 years equivalent to ∼27 years for humans ^43^). This suggests that oocytes of older macaques maintain the quality of their mtDNA, which might be also true for humans. Since our samples were obtained from women undergoing IVF treatments, the age of donors was limited to 20-42—a subset of the human female reproductive life span (∼13 to ∼50 years ^44,45^, with fertility peaking around age 24 and diminishing after 30 ^46^). We therefore cannot exclude the possibility of an increase in mtDNA mutations at an earlier age in humans, similar to what we observed in macaques, sampled by us starting from the age of 1 year (equivalent to ∼7 years in humans). Further studies including younger women will be required to study such a potential early age effect. Furthermore, increased mtDNA mutations may occur at more advanced age, however, mitochondria with a high number of deleterious mutations might be removed by mitophagy ^reviewed^ ^in^ ^47^, or oocytes with high mtDNA mutational load might be eliminated by follicular atresia ^reviewed^ ^in^ ^48^.

Results from previous studies directly assessing mtDNA mutation increases in human oocytes have been contradictory and might have been affected by PCR and sequencing errors, as no previous studies used highly accurate duplex sequencing. Nevertheless, our findings are in agreement with a study analyzing 53 human oocytes, which identified 17 variants with MAF>2% and showed no increase in their frequency with ovarian aging ^23^. In contrast, Yang and colleagues reported higher numbers of mtDNA mutations with MAF>0.2% in oocytes from older vs. younger women ^49^. Their study analyzed oocyte pools (and not single oocytes) with an unreported number of oocytes in each pool; a higher number of oocytes in pools of older women could have led to a higher number of mutations. In general, using a less accurate method, they observed >8,000, or ∼4 times more mutations in 76 oocytes as compared to our observations of 1,877 mutations in 80 oocytes, suggesting the presence of sequencing errors in their study.

Another study ^24^ showed a correlation between age and the number of mtDNA *de novo* mutations with MAF≥2%. A possible explanation for this discrepancy in results might be a different selective pressure acting on variants measured at higher MAFs and/or a change in this pressure with age. To evaluate this for our data, we also considered only mutations with MAF≥2%; these mutations do appear to increase with age, but non-significantly. Indeed, when including only 22 oocyte mutations with MAF≥2% (to ensure an accurate representation of MAFs, only the 74 oocytes with a mean DCS depth >200x were included in this analysis), we found 7, 5, and 10 mutations analyzing 24, 23, and 27 oocytes, in the young, intermediate, and old age groups, respectively. This represents a 1.27-fold higher, not significant (Fisher’s Exact test *p*=0.809) mutation frequency increase in old versus young women. Larger numbers of mutations would be required to gain statistical power.

### Somatic mutations increase with age

Though we had limited power to study somatic mutations in our data because of low mtDNA enrichment and sequencing depth (partly explained by the low mtDNA copy number in these samples), we found a significant and marginally significant increase in mutation frequency in blood and saliva, respectively. Our observed mutation frequencies are in the same range as previous reports in human blood ^50^ and other somatic tissues ^19,31^. Interestingly, neither in blood nor in saliva, we observed a strong enrichment of mutations in the D-loop, which is in contrast to germline mutations as well as mutations measured in the brain ^19^, but is in agreement with previous reports for human lymphocytes and monocytes ^51^, suggesting tissue-specific differences in mtDNA mutational patterns ^52^.

### Mechanisms of mitochondrial mutations

The predominance of transition mutations, combined with strand bias, in all tissues assessed is consistent with a previously suggested major contribution of replication-associated errors in generating mtDNA mutations ^16,19,53,54^. Prior studies have shown that the polymerase responsible for mtDNA replication (DNA polymerase gamma) has a propensity for transition mutations ^55–57^. Another potentially contributing mechanism is the spontaneous deamination of cytosine (C>T) and adenosine (A>G) ^58,59^—also leading to transitions. The strong strand bias we observed for G>A over C>T and for T>C over A>G on the reference L-strand—especially outside of the D-loop—is consistent with a high incidence of C>T and A>G mutations occurring on the H-strand. A similar strand bias was reported in other studies ^16,19,53,54^. This high incidence of mutations on the H-strand might be explained by its single-stranded status during the initial stages of replication (reviewed in ^60^), which facilitates DNA damage, potentially accounting for the nucleotide content differences between the H- and L-strands (the H-strand is guanine- and thymine-rich). Note that in oocytes, the presence of strand bias in the D-loop is inconsistent with a replication-driven mutation pattern expected based on the bidirectional mtDNA replication operating there ^53,61^. As somatic cells from older women have undergone more cycles of replication and hence spent more time in a single-stranded state, our observation of A>G/T>C transitions being the only mutation type with a significant age-associated increase in blood, coupled with increased strand bias in older women, indicates a contribution of replication to their age-associated mutagenesis. Furthermore, our observation of higher rates of C>T/G>A transitions at CpG than non-CpG sites in somatic tissues suggests a role of spontaneous deamination of methylated cytosines ^58^ in mtDNA mutagenesis, despite controversial reports regarding CpG methylation in mtDNA ^e.g., 62–66^. This pattern was not observed in oocytes.

### Does selection impact the germline and somatic mtDNA mutational landscape?

We did not find strong evidence of selection operating on low-frequency mtDNA *de novo* mutations in oocytes. Such mutations might not be subject to selection due to their low frequency, at which they are not expected to have phenotypic effects on a cellular or mitochondrial level. Consistent with this, the distribution of low-frequency (MAF<1%) mutations in oocytes between coding and non-coding regions of mtDNA was more in line with expectations based on the lengths of these regions than that for mutations with higher frequency (Fig.4, **Table S15**). Nevertheless, we observed a lower incidence of disease-associated mutations in oocytes compared to somatic tissues. Thus, purifying selection might be acting against such mutations in oocytes.

When we analyzed germline mutations with higher frequencies, we observed a shift in their distribution from coding towards non-coding regions. This was the case for *de novo* mutations with MAF≥1, and for inheritable heteroplasmies, suggesting frequency-dependent purifying selection in the germline. Specifically, selection would act against higher frequency functional variants, i.e. variants located in coding regions. Consistent with this, previous reports of selection have been based on mtDNA variants with higher frequencies (≥1-5%, observed in pedigrees), mainly reporting purifying selection ^16–18,40^ in the female germ line ^40^.

Age-dependent selection after fertilization acting on variants occurring at higher MAFs could explain why some other studies have reported increased numbers of mutations in children of older mothers ^17,18^. Selective forces acting after fertilization and changes in selection pressure with aging, leading to increased numbers of variants measured at higher MAFs, might be responsible for this observation instead of increased oocyte mutagenesis *per se*. Consistent with this hypothesis, López-Otín and colleagues have previously suggested that the mtDNA quality control system in cells is less efficient at purging pathogenic mutations (or organelles with such mutations) with advancing age ^67^. Duplex sequencing of pedigree mutations would allow one to include variants at lower frequencies, which may further elucidate the causes of differences observed between our low-MAF oocyte variants and published pedigree data.

In contrast to the observed purifying selection acting on high-frequency variants in the germline, our observation of increased hN/hS ratios in blood and saliva is indicative of positive selection. This agrees with previous studies, reporting positive selection in human liver ^32^ and macaque liver and skeletal muscle ^27^, and therefore appears to result from a tissue-type independent mechanism. Alternatively, depending on the impact of these mutations on mitochondrial functions, destructive selection (i.e. selection promoting an increase in abundance of deleterious mutations) ^68^ might explain this observation. In agreement with this, we observed high prevalence of disease-associated mutations, suggesting positive or destructive selection acting on these sites, especially in somatic tissues.

Furthermore, high-frequency inheritable heteroplasmies were observed in a higher proportion of women as compared to the proportion of oocytes, which is indicative of positive selection in somatic tissues, and leads to increases in the MAFs of variants. Indeed, 54.5% of women in our study carried an inheritable heteroplasmy measured at a MAF ≥1% (in blood and/or saliva), which is in agreement with previously reported figures for mothers (47.8%) and offspring (42.5%) ^45^. In contrast, only 35.0% of all oocytes we assessed carried a heteroplasmy above this MAF threshold. This is also in agreement with our observation that, lowering the MAF, one finds higher numbers of inheritable heteroplasmies in oocytes compared to somatic tissues. MAFs measured in somatic tissues might increase with age, leading to more high-frequency variants in older people. In line with this, Liu and colleagues reported higher numbers of high-frequency variants (MAF≥1%) in somatic cells of older vs. younger mice ^69^.

### The effective germline bottleneck size depends on mutation frequency

By studying inheritable heteroplasmies in oocytes, we were able to estimate the size of the effective germline mtDNA bottleneck directly in human germ cells, resulting in an estimate that is not affected by age-related genetic drift in somatic tissues. Furthermore, the use of highly accurate duplex sequencing allowed us to include in the estimation heteroplasmies with MAFs <1%. Considering all inheritable heteroplasmies, we estimated the effective germline bottleneck size to be ∼909 segregating units (CI: 0.47-5,329.73). Considering only heteroplasmic sites for which at least one sample had MAF≥1%, we estimated a bottleneck size of ∼30 segregating units (CI: 0.02-178.53). This is similar to previous estimates in humans ranging between ∼3-32 ^16–18,70^ using MAF cutoffs of 1% or higher. In fact the study using the highest MAF cutoff (only including heteroplasmies with MAFs between 10% and 90%) yielded the smallest bottleneck size estimate of around 3 ^16^. The similarity between our bottleneck size estimates based directly on oocytes and prior estimates based on pedigrees suggests that the calculation is only negligibly affected by genetic drift in somatic tissues. Differences in bottleneck size estimates depending on MAF cutoffs suggest differences in selective pressure acting on variants with different frequencies. Considering the organization of multiple mtDNA copies in individual mitochondria and cells, selection likely has a different impact when a larger proportion of mtDNA molecules are affected, which is the case for variants at higher MAFs. Note that wide, overlapping confidence intervals have to be considered in these comparisons.

### Germline mtDNA mutation rate estimates

To compare germline mutation rate estimates between our study and prior ones, we computed a germline mutation rate of 3.95×10^-4^ (95% CI 2.68-5.60×10^-4^) mutations per position per generation for the D-loop, including only *de novo* mutations measured at MAF≥1%. This is about 5× higher than a previous estimate by Wei and colleagues based on a pedigree analysis using a similar MAF cutoff^40^. When including only mutations measured at MAFs ≥5%, our mutation rate estimate for the coding region is 2.50×10^-6^ (95% CI 5.16×10^-7^-7.32×10^-6^) mutations per position per generation—which is in the same range as the estimate produced by Árnadóttir and colleagues based on pedigrees and using the same MAF cutoff ^16^ (2.87×10^-6^,mutations per position per generation; 95% CI 2.79-2.95×10^-6^). Again, to obtain a more complete picture of potential differences in mutation rate estimated based on oocytes and pedigree studies, duplex sequencing of pedigrees could also aid in addressing mutations at lower MAFs, which are currently excluded from such studies.

### Potential caveats

When calling *de novo* mtDNA mutations, we utilized several experimental and computational steps to minimize the contribution of Numts (e.g., a mtDNA enrichment strategy not prone to co-enrichment of Numts, as previously described in detail for macaques ^27^). The enzymatic digestion of linear nuclear DNA, the comparatively long fragments after shearing, and the relatively high read length during sequencing, as well as retaining only reads mapping to mtDNA, all minimize the contribution from Numts to our mutation dataset. Moreover, while we cannot completely exclude their contribution, Numts should affect each woman similarly and not bias our comparison of mutations between the three age groups. Considering the efficient enrichment of mtDNA molecules in oocytes (on average, 86.1% of all DCS reads represented mtDNA sequences), we do not expect Numts to impact any interpretations considerably. Furthermore, when inferring MAFs of inheritable heteroplasmies, their frequencies were often based on small numbers of measured variants at a position in a sample. While due to the low error rate of duplex sequencing we can be confident that the variant is present in a sample, MAFs might be over- or underestimated.

## Materials and Methods

### Study participant details, samples, and their handling

All parts of this study were designed and conducted in compliance with the relevant regulations regarding the use of human study participants, and in accordance with the Declaration of Helsinki. Duplex sequencing was performed for somatic tissues and oocytes of 22 women who were scheduled for intracytoplasmic sperm injection (ICSI) at the Kinderwunsch Zentrum (fertility center) of the Kepler University Clinic, Linz, Austria. The women spanned an age range from 20 to 42 years. The study was approved by the Ethics Commission of Upper Austria (F-11-17), and followed the rules of informed consent, anonymization of samples, and avoidance of any ethical sensitivity that may harm an individual. In total, 80 single oocytes were sequenced. Moreover, upon availability and exonuclease V enrichment success, blood from 18 donors and/or saliva from 8 donors (for 4 donors both samples were available) were sequenced (**Table S1**). For an additional three women (hs001, hs002, and hs004), saliva was sequenced using the targeted mtDNA capture approach; those sequences were only included in the analyses of inheritable heteroplasmies.

As described previously ^26,27^, to avoid cross-sample contamination, all pre-PCR steps were performed in designated laminar-flow hoods located in a separate room. To minimize the time of sample processing with an open lid, individual samples were handled in separate low-binding tubes (Axygen). Only after DNA amplification, samples were transferred to a 96-well plate format during bead purifications with the epMotion automated liquid handling system (Eppendorf). Afterwards, samples were transferred back to separate tubes, minimizing potential contamination.

### Sample preparation

#### Single oocyte collection

The 22 ICSI female patients included in this study either had male partners who suffered from subfertility (59.1%), or themselves suffered from infertility that was not expected to cause germline mutations (e.g., bilateral tubal blockage or anovulation; 40.9%). Controlled ovarian hyperstimulation was performed. For the vast majority of patients (n=19, 86.4%) this was done using a short antagonist protocol, while three patients were treated with a long agonist regimen. On average, women were stimulated with 1,913 (±777) IU of purified (Menopur, Ferring; Fostimon, IBSA) or recombinant FSH (Puregon, MSD) and stimulation took 11 (± 2) days. Ovulation was induced with 10,000 IU hCG (Pregnyl, MSD). Oocyte retrieval was carried out transvaginally under ultrasound guidance 36 hours after ovulation induction.

Cumulus-oocyte-complexes (COC) were cultured in GM501 Wash medium (Gynemed) until further processing. Since ICSI requires removal of surrounding cumulus cells from the zona pellucida (ZP) of the oocytes, enzymatic digestion of the same was performed for 30 seconds (GM501 hyaluronidase, Gynemed, 80 IU). Remaining cumulus cells still obstructing the view were then removed mechanically until the maturity of the gamete could be evaluated properly.

In principle, all metaphase II (MII) oocytes were considered for ICSI and could not be used for study purposes. The only exception was a dysmorphism called ‘cluster of the smooth endoplasmic reticulum’ ^71^. Such affected MII-oocytes (n=13) were never injected but instead were transferred into a vial containing 2 µl phosphate buffered saline and immediately stored at -20°C.

All immature oocytes were potential research objects. In these cases, culture was attempted overnight to have the immature gametes reach MII stage. All cumulus cells that did not automatically detach through the manipulation process conducted the day before were removed using a diode laser (Fertilase, Vitrolife), so that finally the ZP of the gametes was completely free of somatic cells (this was done to exclude potential mutational bias coming from those cells). In cases where telophase I (TI) or MII was reached (as assessed by the presence of the first polar body within the perivitelline space) the ZP was opened by a series of laser pulses so that the first polar body could be biopsied by means of a glass pipette (15µm diameter, Gynemed). Polar bodies and oocytes were separately frozen as described above. As a result, oocytes included in this study comprise: MII oocytes (n=62), germinal vesicles (n=4), metaphase I oocytes (n=8), and TI oocytes (n=6). All oocytes were initially stored at -20°C, and later transferred to -80°C for longer-term storage until further analysis. No oocytes included in this study were ever exposed to sperm.

#### DNA extraction and mtDNA enrichment from somatic tissues

Total DNA was extracted from 6-9 ml EDTA-blood using the FlexiGene DNA Kit (Qiagen) according to manufacturer’s instructions. Saliva samples were collected using OG-500 tubes (DNA Genotek) and total DNA was extracted using the prepIT L2P Kit (DNA Genotek) according to manufacturer’s instructions.

The extracted DNA was enriched for mtDNA, using exonuclease V which selectively digests linear DNA while preserving the circular mtDNA ^72^, as described previously ^26,27^. This step was necessary to minimize the contribution from Numts ^73^. In short, ∼2.5 µg of total DNA was incubated at 37°C for 48 h in 60-µl reactions containing 1× NEB 4 buffer, 30 U Exonuclease V (RecBCD, NEB), and 2 mM ATP. After this first incubation, the reaction volume was increased to 100 µl by supplementing with further NEB 4 buffer (final concentration = 1×), 20 U Exonuclease V, ATP (final concentration = 2 mM), and 5 µg RNase A (Amresco) and incubated at 37°C for another 16 h. The reactions were purified using 1 volume of AMPure XP beads (Beckman Coulter).

#### Oocyte lysis and mtDNA enrichment

Oocytes were lysed and mtDNA enriched as previously described ^26,27^. In short, 4 µl cold oocyte lysis buffer (50 mM Tris-HCl, pH 8.0 (Invitrogen), 1 mM EDTA, pH 8.0 (Promega), 0.5% Polyoxywthylene (20) Sorbitan Monolaureate (OmniPur), and 0.2 mg/ml Proteinase K (IBI Scientific) were added to a single oocyte in a low-binding tube, incubated at 50°C for 15 min, followed by an overnight incubation at 37°C. Proteinase K was then partially inactivated by incubating at 65°C for 15 min. 1 mM ATP, 10 mM MgCl_2_, 10 mM Tris-HCl, pH 8.0, and 10 U Exonuclease V (RecBCD, NEB) were added to obtain a final volume of 13 µl, followed by incubation at 37°C for 1 h. 10 ng RNase A (Amresco) were added, and reactions were incubated at 37°C for 15 min. TE buffer was added up to the final volume of 50 µl. Enzymes were inactivated by incubation of the samples at 70°C for 30 min.

#### mtDNA enrichment estimation

mtDNA enrichment was estimated by real-time PCR, amplifying a region of the mitochondrial *ND6* gene, as well as nuclear *Alu* elements. Samples were amplified using primers specific for mtDNA (F-hsND6-mtDNA: CCACAGCACCAATCCTACCT and R-hsND6-mtDNA: GTCAGGGGTTGAGGTCTTGG (both IDT); NC_012920.1: positions 14,259-14,525) as well as for nuclear DNA (F-primateAlu: CCTGTAATCCCAGCACTTTG and R-primateAlu: TAGTAGAGACGGGGTYTCAC (both IDT), amplifies multiple regions in the genome). Reactions contained 2 µl 1:10 diluted ligation reaction, 1× PowerUp SYBR Green Master Mix (Thermo-Fisher Scientific), and 0.2 µM each primer and were amplified at 95°C for 2 min, 45 cycles of 95°C for 15 sec, 57°C for 20 sec, and 72°C for 30 sec, followed by 72°C for 2 min and a melting curve analysis. A standard consisting of tissue samples that where previously enriched for mtDNA at 9 different efficiencies (1.8%, 11.5%, 17.5%, 30.0%, 33.1%, 59.5%, 61.6%, 69.2%, and 93.0% of mtDNA in the total sample as determined using NGS), was used to estimate the mtDNA enrichment.

### Duplex library preparation and sequencing

Duplex libraries were prepared as described previously ^26,27^. In short, the enriched DNA was sheared with a Covaris M220 Focused-ultrasonicator using 50 µl reactions in microTUBE-50 AFA Fiber Screw-Cap (Covaris), using the protocol for ∼550 bp fragments. Samples were transferred into 200 µl low-binding tubes (Axygen) for library preparation. Duplex sequencing libraries were prepared following a previously described protocol ^74^ with several modifications, which are provided in more detail in a prior study ^27^. In short, T-tailed duplex adapters were prepared with sequences MWS51 and MWS55. End-repair and A-tailing were performed with the End Prep module of the NEBNext Ultra II DNA Library Prep Kit for Illumina (NEB) according to manufacturer’s instructions, followed by adapter ligation with the NEBNext Ultra II Ligation Mix and NEBNext Ligation Enhancer, using ∼25 fmole duplex adapters for somatic tissues, and ∼18 fmole for oocytes. Adapter-ligated DNA was purified twice with 0.8 volumes AMPure XP beads (Beckman Coulter) and eluted in 15 µl 10 mM Tris-HCl (pH 8.0). The amount of amplifiable fragments and hence the input for tag-family PCR was estimated with real-time PCR: 2 µl of the 1:10 diluted adapter-ligated library aliquot were amplified in 10-µl reactions containing 1 µM NEBNext Universal PCR Primer for Illumina (NEB), 1 µM NEB_mws20 primer (GTGACTGGAGTTCAGACGTGTGCTCTTCCGATC*T (IDT); the asterisk indicates a phosphorothioate bond), 1× Kapa HiFi HotStart Reaction Mix (KAPA Biosystems), and 1× EvaGreen Dye (Biotium). Reactions were amplified in a CFX96 Real-Time PCR machine (Bio-Rad) at 98°C for 45 sec, 35 cycles of 98°C for 15 sec, 65°C for 30 sec, and 72°C for 45 sec, followed by 72°C for 2 min and a melting curve analysis. PCR products were additionally inspected on a 1.5% agarose gel. Tag-family PCR reactions contained up to 14 µl adapter-ligated sample. If the Cq-value in the real-time PCR was ≥23, all 14 µl of the sample were used as PCR template, samples with lower Cq-values were diluted accordingly. For samples with a real-time PCR Cq-value of ∼23, 7M paired-end reads were anticipated during sequencing, ensuring efficient DCS formation. For tag-family PCR, first, 12 cycles of linear amplification were performed in 38 µl reactions: 14 µl (accordingly diluted) DNA, 1× Kapa HiFi HotStart Reaction Mix (KAPA Biosystems), 1 µM NEB_mws20 primer at 98°C for 45 sec, 12 cycles of 98°C for 15 sec, 60°C for 30 sec, and 72°C for 45 sec, followed by 72°C for 2 min. 1 µM NEBNext Universal PCR Primer for Illumina (NEB) was added to each reaction followed by amplification at 98°C for 45 sec, 9 cycles of 98°C for 15 sec, 65°C for 30 sec, and 72°C for 45 sec, followed by 72°C for 2 min. PCR products were purified with 0.8 volumes AMPure XP beads (Beckman Coulter) using the epMotion automated liquid handling system (Eppendorf), and different indexes were added via PCR: in 50 µl reactions, 15 µl purified library was amplified with 1 µM NEBNext Universal PCR Primer for Illumina (NEB), 1 µM NEBNext Multiplex Oligos for Illumina (Index Primers Sets 1-4), and 1× Kapa HiFi HotStart Reaction Mix (KAPA Biosystems) at 98°C for 45 sec, 9-16 cycles of 98°C for 15 sec, 65°C for 30 sec, and 72°C for 45 sec, followed by 72°C for 2 min. The number of required cycles was estimated based on the amount of sample used as input in the linear PCR. PCR products were purified with 0.8 volumes AMPure XP beads (Beckman Coulter) using the epMotion automated liquid handling system (Eppendorf), their quality checked with a Bioanalyzer High Sensitivity DNA Kit (Agilent), quantified with the KAPA Library Quantification Kit (Kapa Biosystems) according to manufacturer’s instructions, and pooled depending on the number of reads needed per sample to obtain a family size around six. Sequencing was performed on an Illumina HiSeq 2500 platform with 250 nt paired-end reads.

#### Targeted mtDNA capture sequencing

Since mtDNA enrichment with exonuclease V turned out to be inefficient for low mtDNA copy number samples such as blood and saliva, mtDNA in somatic tissues was also enriched using the targeted mtDNA capture approach, as previously described for duplex sequencing ^75^. This was done for the 18 blood and for the 11 saliva samples (see **Table S1** and the section on Study participant details, samples, and their handling for more details). Duplex libraries were prepared as described above (with minor modifications), starting with 500 ng of total DNA in a volume of 50 µl, sheared to ∼550 bp with a Covaris M220 Focused-ultrasonicator. Modifications included the use of ∼25 pmole duplex adapters and 120 ng adapter-ligated DNA as input for tag-family PCR (run for 3 cycles; no linear amplification).

Targeted mtDNA capture was performed using the xGen Hybridization and Wash Kit and the xGen Human mtDNA Research Panel v1.0 (both IDT) according to manufacturer’s instructions, with minor modifications for compatibility with the duplex sequencing protocol. Modification included the use of the previously published mws60 and mws61 blocking oligos ^75^ and the post-capture (indexing) PCR that was performed in 60 µl reactions as follows: 20 µl purified library was amplified with 1 µM NEBNext Universal PCR Primer for Illumina (NEB), 1 µM NEBNext Multiplex Oligos for Illumina (Index Primers Sets 1-4), and 1× Kapa HiFi HotStart Reaction Mix (KAPA Biosystems) at 98°C for 45 sec, 15 cycles of 98°C for 15 sec, 65°C for 30 sec, and 72°C for 30 sec, followed by 72°C for 2 min. For each sample, ∼9M paired-end reads were anticipated during sequencing. Sequencing was performed on an Illumina HiSeq 2500 platform with 250 nt paired-end reads.

### Consensus read formation, mapping, and variant calling

Single-strand (SSCS) and duplex (DCS) consensus formation was performed in Galaxy ^76^ using the Du Novo pipeline (version 2.14) including barcode error correction ^28,29^, as described previously ^26,27^, and quality of the sequencing reads was verified using FastQC. In short, barcode error correction allowed ≤3 mismatches and required a mapping quality ≥20; a family size ≥3 was required for consensus formation, calling a mutation if present in ≥70% of the sequencing reads. To avoid errors associated with end-preparation, the first 12 nts of the consensus sequences were trimmed using trimmomatic (HEADCROP) ^77^. Trimmed DCSs and SSCSs were mapped to the human reference genome (hg38, which contains the rCRS mtDNA reference sequence NC_012920.1) using BWA-MEM ^78^.

Fixed differences between individual donors were used to check for potential cross-sample contamination during library preparation. When comparing libraries prepared in the same batches, no evidence of cross-sample contamination was found. mtDNA enrichment efficiency was estimated by calculating the percentage of sequences mapping to the mtDNA reference. To minimize mapping bias at the edges from mapping a circular genome to a linearized reference, in addition to mapping to the whole mtDNA reference (as part of hg38), reads were mapped to hg38 in which chrM was replaced by a re-linearized reference sequence (consisting of positions 14,570-16,569 and 1-2,000). Positions 16,070-500 were analyzed from the sequences mapped to the re-linearized mtDNA reference version. BAM files were filtered using BAMTools ^79^, requiring a mapping quality >20, mapping to chrM, being the primary alignment, paired reads, proper read pairing, and mapping of the mate sequence. Reads were left-aligned using the Bam Left Align tool.

To avoid a bias in variant frequencies from overlapping paired-end reads, overlaps were clipped using the clipOverlap function of bamUtil ^80^. Variants were called using the Naive Variant Caller (NVC) ^81^ with the following settings: a minimum number of one read needed to be considered as REF/ALT, minimum base quality of 20, minimum mapping quality of 50, and ploidy of 1. DCSs were used for the initial variant calling and for *de novo* mutation analysis, whereas SSCSs were used in addition to DCSs for the analysis of inheritable heteroplasmies (see corresponding section for more details).

### Variant filtering and classification of tissue-specific *de novo* mutations

We identified 4,643 variants; for 11 of these, two different mutant alleles were recorded, however, by default only one allele was displayed (the variant of the second mutant allele was added manually). We excluded from our analyses 468 variants, located at the following 15 mtDNA positions: 302, 310, 312, 315-317, 513, 514, 515, 523, 525, 3107, 16183, 16189, and 16194. These are mainly in repetitive sequences, and the excluded variants were identified as false positive mutations occurring at the beginning or at the end of mononucleotide repeats due to mapping errors. In addition, we excluded 71 donor-specific variants resulting from repetitive sequences at the following positions: hs022 545 G>C, hs022 564 G>A, hs022 567 A>C, hs022 574 A>C, hs022 8281 C>A, hs004 8279 T>C, hs015 8279 T>C, hs022 8285 C>T, hs004 8281 C>G, hs015 8281 C>G, hs011 16186 T>C, hs015 5895 C>A, hs015 5899 C>A, hs003 16180 A>C, hs005 16180 A>C, hs015 16180 A>C, hs005 16181 A>C, hs005 16181 C>A, and hs005 16182 C>A. We also excluded 222 linked variants (with >3 variants present in single paired-end reads, mostly occurring in multiple unrelated samples) which represent potential Numts. Among the remaining variants, we further excluded 146 variants with ambiguous timing of occurrence during development (for these, we could not unambiguously determine if they represented inheritable heteroplasmies, early somatic mutations, or early germline mutations). A total of 148 inheritable heteroplasmies (i.e. variants present in all somatic tissues assessed and ≥1 oocyte(s) of a donor) were analyzed separately, as described in more detail in the section ‘Analysis of inheritable heteroplasmies’. Finally, 63 low-confidence variants identified in oocytes with a mean DCS depth <50× (originating from 2 oocytes) were excluded since the computation of mutation frequencies could be biased by the low number of total nucleotides sequenced for each oocyte. As a result, we obtained 3,525 high-confidence tissue-specific *de novo* mutations. Mutations measured in >1 DCS per sample were counted as a single mutation, assuming that they resulted from a single mutation event. Mutation frequency was computed as the number of mutations divided by the product of mtDNA length (16,569 bp) and mean DCS depth across samples.

### Analysis of *de novo* mutations

#### Generalized mixed-effects linear model to assess the effect of age

The effect of age on mutations was assessed using a generalized mixed-effects linear model, as described previously ^27^. In short, for each sample, we computed the mutation frequency as the number of observed mutations divided by the total number of sequenced nucleotides, and employed the R function `glmer` (from the package `lme4`) to fit a generalized mixed-effects linear model with binomial family and logit link, the mutation frequency as response and the numbers of sequenced nucleotides as weights (prior weights are used to provide the number of trials in a binomial model when the response is the proportion of successes). We included age, the categorical variable tissue, and their interaction as fixed-effect predictors, while donor ID was employed as a random effect. The variable age was standardized prior to fitting the model in order to avoid convergence issues. The Wald test—as provided by the summary of the fitted glmer model—was employed to ascertain whether each of the coefficients was null. Age effect, tissue effect, and random effects were tested using the likelihood ratio test based on the Chi-square distribution provided by the function anova.merMod. The conditional modes of the random effects were extracted using the function `ranef`. We also tested the effect of age for each of the tissues considered using the Z-test provided by the function `glht` of the R package `multcomp`. Marginal and conditional pseudo-*R*² (representing the variability explained by the fixed effects and by the entire model, respectively) were computed using the function `r.squaredGLMM` (from the R package `MuMIn`). Predictions of mutation frequencies based on the fixed-effect part of the model were computed using the function predict.merMod from the R package lme4, while bootstrap pointwise confidence intervals based on the quantile approach were computed using the function `bootMer` (R package `lme4`).

#### Estimation of the rate of nonsynonymous vs. synonymous mutations

We tested for selection in protein-coding sequences by calculating *hN*/*hS* ratios, as described in ^32^. *hN* is the number of nonsynonymous variants per nonsynonymous sites, and *hS* is the number of synonymous variants per synonymous sites. The numbers of nonsynonymous and synonymous sites were calculated using the Nei-Gojobori method ^82^. Under neutrality, the *hN*/*hS* ratio is expected to be equal to 1; a ratio <1 is suggestive of purifying selection, and a ratio >1 is suggestive of positive selection. The neutral distribution of *hN*/*hS* was generated by bootstrapping ^32^ (100 replicates) from the observed mutation spectrum, and used to assess if the observed *hN*/*hS* ratio deviates from neutral expectations.

#### Identification of variant hotspots

Variant hotspots were identified as described previously ^27^. We define variant hotspots as mutations that are shared by multiple donors, but are not expected to be shared by chance (168 mutations were shared in oocytes and 161 mutations were shared in somatic tissues). Blood and saliva were considered together as ‘somatic tissues’ due to similar mutation frequencies and lower numbers of measured mutations, hence increasing our power to detect somatic variant hotspots. For the evaluation of oocyte variant hotspots, we also considered the number of sequenced oocytes per donor (ranging from 1 to 5).

In short, we built a probabilistic model that takes into account tissue-specific estimates of the average mutation frequencies inside and outside of the D-loop, as well as the mean DCS depth in each sample. We computed the expected number of mutations present in exactly one woman and shared by several women, separately for oocytes and somatic tissues. As a result, we defined oocyte variant hotspots as sites mutated in oocytes of at least four different women, and somatic hotspots as sites mutated in somatic tissues of at least three different women.

#### Annotation of disease-associated variants

Mutations known to be associated with diseases were annotated based on the 109 disease-associated confirmed substitutions reported by MITOMAP ^33^ (downloaded on June 18, 2024). They were located in protein-coding regions (n=56), tRNA (n=51), or rRNA (n=2).

### Analysis of inheritable heteroplasmies

#### Identification of inheritable heteroplasmies

All sequenced oocytes and somatic tissues were included in the analysis of inheritable heteroplasmies. MAFs obtained from variant calling on DCSs were primarily used in the analysis of inherited heteroplasmies. However, if the sequencing depth for a minor allele was <3 for DCSs, the MAFs obtained from the SSCS analysis were also used, as described previously ^26,27^. For somatic tissues, in addition to exonuclease V mtDNA enriched sequencing libraries, we included duplex libraries enriched for mtDNA using a targeted capture approach. Though such samples harbor the chance of Numt co-enrichment, the sole assessment of variants previously identified in donors using Exonuclease V mtDNA enrichment diminishes such bias. This increased the average SSCS sequencing depth from 445× and 514× to 3,750× (n=18, SD=824.6) and 5,019× (n=11, SD=1,864.1) per site per sample in blood and saliva, respectively. MAFs measured using the two different enrichment strategies correlated well (**Fig. S9**).

#### mtDNA bottleneck calculation

The size of the effective germline mtDNA bottleneck was estimated as described previously ^27^. In short, the effective mtDNA bottleneck *N* was calculated by first computing *N_ij_* for individual *i* and position *j* as

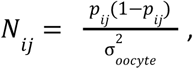

where *p_ij_* is the mean allele frequency of blood and saliva (if both sample types were available), and the denominator is the mean squared deviation (across *n_ij_* oocytes for the same donor and position) calculated as

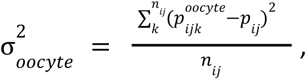

where *p^oocyte^_ijk_* is the allele frequency for the *k^th^* oocyte. To obtain *N*, we averaged the *N_ij_*′s. The 95% confidence intervals were generated by bootstrapping (sampling with replacement 1,000 times).

### Statistical analysis: Significance tests

One-sided permutation tests were performed to assess whether the median mutation frequencies were higher in older age groups for each of the tissues analyzed, using 10,000 permutations. Larger expected quantities in older donors justify the use of a one-sided test. Significance of age-related increases after combining all mutations across samples belonging to the same group (e.g. mutations in blood of young women) was assessed using the Fisher’s exact test (function `fisher.test` from the `stats` package in R). *p*-values were corrected for multiple comparisons (function `p.adjust` from the `stats` package in R) using the method of Benjamini, Hochberg, and Yekutieli to control the false discovery rate ^83^.

## Resource availability

The sequencing reads are available at SRA under accession PRJNA1181658. The data were analyzed in R using the packages listed above. Tables with all mutations were uploaded to GitHub and are available at https://github.com/makovalab-psu/human_mtDNA_duplexSeq.

## Acknowledgments

We are grateful to Nicholas Stoler for developing the duplex sequencing instance on Galaxy and providing advice on the analysis, and to Edmundo Torres Gonzalez for his comments on an earlier version of the manuscript. We thank all sample donors for their contributions.

## Author contributions

Conceptualization: B.A., I.T.B., T.E., and K.D.M. Methodology: B.A. Software: B.A. Validation: B.A. and K.D.M. Formal analysis: B.A. Investigation: B.A., K.A., B.H., and T.E. Resources: P.O., O.S., and T.E. Writing - Original Draft: B.A. and K.D.M. Writing - Review & Editing: B.A., F.C., I.T.B., P.O., O.S., T.E., and K.D.M. Visualization: B.A. Supervision: P.O. and K.D.M. Funding acquisition: B.A. and K.D.M.

## References

1. Suomalainen, A., and Nunnari, J. (2024). Mitochondria at the crossroads of health and disease. Cell 187, 2601–2627. 10.1016/j.cell.2024.04.037.

2. Taylor, R.W., and Turnbull, D.M. (2005). Mitochondrial DNA mutations in human disease. Nat. Rev. Genet. 6, 389–402. 10.1038/nrg1606.

3. Dabravolski, S.A., Khotina, V.A., Sukhorukov, V.N., Kalmykov, V.A., Mikhaleva, L.M., and Orekhov, A.N. (2022). The Role of Mitochondrial DNA Mutations in Cardiovascular Diseases. Int. J. Mol. Sci. 23. 10.3390/ijms23020952.

4. Gorman, G.S., Schaefer, A.M., Ng, Y., Gomez, N., Blakely, E.L., Alston, C.L., Feeney, C., Horvath, R., Yu-Wai-Man, P., Chinnery, P.F., et al. (2015). Prevalence of nuclear and mitochondrial DNA mutations related to adult mitochondrial disease. Ann. Neurol. 77, 753–759. 10.1002/ana.24362.

5. Pakendorf, B., and Stoneking, M. (2005). Mitochondrial DNA and human evolution. Annu. Rev. Genomics Hum. Genet. 6, 165–183. 10.1146/annurev.genom.6.080604.162249.

6. Timmis, J.N., Ayliffe, M.A., Huang, C.Y., and Martin, W. (2004). Endosymbiotic gene transfer: organelle genomes forge eukaryotic chromosomes. Nat. Rev. Genet. 5, 123–135. 10.1038/nrg1271.

7. Rath, S.P., Gupta, R., Todres, E., Wang, H., Jourdain, A.A., Ardlie, K.G., Calvo, S.E., and Mootha, V.K. (2024). Mitochondrial genome copy number variation across tissues in mice and humans. Proceedings of the National Academy of Sciences 121, e2402291121. 10.1073/pnas.2402291121.

8. Shoubridge, E.A., and Wai, T. (2007). Mitochondrial DNA and the mammalian oocyte. Curr. Top. Dev. Biol. 77, 87–111. 10.1016/S0070-2153(06)77004-1.

9. Pagnamenta, A.T., Wei, W., Rahman, S., and Chinnery, P.F. (2021). Biparental inheritance of mitochondrial DNA revisited. Nat. Rev. Genet. 22, 477–478. 10.1038/s41576-021-00380-6.

10. Halldorsson, B.V., Palsson, G., Stefansson, O.A., Jonsson, H., Hardarson, M.T., Eggertsson, H.P., Gunnarsson, B., Oddsson, A., Halldorsson, G.H., Zink, F., et al. (2019). Characterizing mutagenic effects of recombination through a sequence-level genetic map. Science 363. 10.1126/science.aau1043.

11. Jónsson, H., Sulem, P., Kehr, B., Kristmundsdottir, S., Zink, F., Hjartarson, E., Hardarson, M.T., Hjorleifsson, K.E., Eggertsson, H.P., Gudjonsson, S.A., et al. (2017). Parental influence on human germline de novo mutations in 1,548 trios from Iceland. Nature 549, 519–522. 10.1038/nature24018.

12. Kong, A., Frigge, M.L., Masson, G., Besenbacher, S., Sulem, P., Magnusson, G., Gudjonsson, S.A., Sigurdsson, A., Jonasdottir, A., Jonasdottir, A., et al. (2012). Rate of de novo mutations and the importance of father’s age to disease risk. Nature 488, 471–475. 10.1038/nature11396.

13. Kessler, M.D., Loesch, D.P., Perry, J.A., Heard-Costa, N.L., Taliun, D., Cade, B.E., Wang, H., Daya, M., Ziniti, J., Datta, S., et al. (2020). De novo mutations across 1,465 diverse genomes reveal mutational insights and reductions in the Amish founder population. Proc. Natl. Acad. Sci. U. S. A. 117, 2560–2569. 10.1073/pnas.1902766117.

14. Wong, W.S.W., Solomon, B.D., Bodian, D.L., Kothiyal, P., Eley, G., Huddleston, K.C., Baker, R., Thach, D.C., Iyer, R.K., Vockley, J.G., et al. (2016). New observations on maternal age effect on germline de novo mutations. Nat. Commun. 7, 10486. 10.1038/ncomms10486.

15. Rahbari, R., Wuster, A., Lindsay, S.J., Hardwick, R.J., Alexandrov, L.B., Turki, S.A., Dominiczak, A., Morris, A., Porteous, D., Smith, B., et al. (2016). Timing, rates and spectra of human germline mutation. Nat. Genet. 48, 126–133. 10.1038/ng.3469.

16. Árnadóttir, E.R., Moore, K.H.S., Guðmundsdóttir, V.B., Ebenesersdóttir, S.S., Guity, K., Jónsson, H., Stefánsson, K., and Helgason, A. (2024). The rate and nature of mitochondrial DNA mutations in human pedigrees. Cell. 10.1016/j.cell.2024.05.022.

17. Rebolledo-Jaramillo, B., Su, M.S.-W., Stoler, N., McElhoe, J.A., Dickins, B., Blankenberg, D., Korneliussen, T.S., Chiaromonte, F., Nielsen, R., Holland, M.M., et al. (2014). Maternal age effect and severe germ-line bottleneck in the inheritance of human mitochondrial DNA. Proc. Natl. Acad. Sci. U. S. A. 111, 15474–15479. 10.1073/pnas.1409328111.

18. Zaidi, A.A., Wilton, P.R., Su, M.S.-W., Paul, I.M., Arbeithuber, B., Anthony, K., Nekrutenko, A., Nielsen, R., and Makova, K.D. (2019). Bottleneck and selection in the germline and maternal age influence transmission of mitochondrial DNA in human pedigrees. Proc. Natl. Acad. Sci. U. S. A. 116, 25172–25178. 10.1073/pnas.1906331116.

19. Kennedy, S.R., Salk, J.J., Schmitt, M.W., and Loeb, L.A. (2013). Ultra-sensitive sequencing reveals an age-related increase in somatic mitochondrial mutations that are inconsistent with oxidative damage. PLoS Genet. 9, e1003794. 10.1371/journal.pgen.1003794.

20. Brown, D.T., Samuels, D.C., Michael, E.M., Turnbull, D.M., and Chinnery, P.F. (2001). Random genetic drift determines the level of mutant mtDNA in human primary oocytes. Am. J. Hum. Genet. 68, 533–536. 10.1086/318190.

21. Gigarel, N., Hesters, L., Samuels, D.C., Monnot, S., Burlet, P., Kerbrat, V., Lamazou, F., Benachi, A., Frydman, R., Feingold, J., et al. (2011). Poor correlations in the levels of pathogenic mitochondrial DNA mutations in polar bodies versus oocytes and blastomeres in humans. Am. J. Hum. Genet. 88, 494–498. 10.1016/j.ajhg.2011.03.010.

22. Ancora, M., Orsini, M., Colosimo, A., Marcacci, M., Russo, V., De Santo, M., D’Aurora, M., Stuppia, L., Barboni, B., Cammà, C., et al. (2017). Complete sequence of human mitochondrial DNA obtained by combining multiple displacement amplification and next-generation sequencing on a single oocyte. Mitochondrial DNA A DNA Mapp Seq Anal 28, 180–181. 10.3109/19401736.2015.1115499.

23. Boucret, L., Bris, C., Seegers, V., Goudenège, D., Desquiret-Dumas, V., Domin-Bernhard, M., Ferré-L’Hotellier, V., Bouet, P.E., Descamps, P., Reynier, P., et al. (2017). Deep sequencing shows that oocytes are not prone to accumulate mtDNA heteroplasmic mutations during ovarian ageing. Preprint, 10.1093/humrep/dex268 https://doi.org/10.1093/humrep/dex268.

24. Mertens, J., Belva, F., van Montfoort, A.P.A., Regin, M., Zambelli, F., Seneca, S., Couvreu de Deckersberg, E., Bonduelle, M., Tournaye, H., Stouffs, K., et al. (2024). Children born after assisted reproduction more commonly carry a mitochondrial genotype associating with low birthweight. Nat. Commun. 15, 1232. 10.1038/s41467-024-45446-1.

25. Schmitt, M.W., Kennedy, S.R., Salk, J.J., Fox, E.J., Hiatt, J.B., and Loeb, L.A. (2012). Detection of ultra-rare mutations by next-generation sequencing. Proc. Natl. Acad. Sci. U. S. A. 109, 14508–14513. 10.1073/pnas.1208715109.

26. Arbeithuber, B., Hester, J., Cremona, M.A., Stoler, N., Zaidi, A., Higgins, B., Anthony, K., Chiaromonte, F., Diaz, F.J., and Makova, K.D. (2020). Age-related accumulation of de novo mitochondrial mutations in mammalian oocytes and somatic tissues. PLoS Biol. 18, e3000745. 10.1371/journal.pbio.3000745.

27. Arbeithuber, B., Cremona, M.A., Hester, J., Barrett, A., Higgins, B., Anthony, K., Chiaromonte, F., Diaz, F.J., and Makova, K.D. (2022). Advanced age increases frequencies of de novo mitochondrial mutations in macaque oocytes and somatic tissues. Proc. Natl. Acad. Sci. U. S. A. 119, e2118740119. 10.1073/pnas.2118740119.

28. Stoler, N., Arbeithuber, B., Povysil, G., Heinzl, M., Salazar, R., Makova, K.D., Tiemann-Boege, I., and Nekrutenko, A. (2020). Family reunion via error correction: an efficient analysis of duplex sequencing data. Preprint, 10.1186/s12859-020-3419-8 https://doi.org/10.1186/s12859-020-3419-8.

29. Stoler, N., Arbeithuber, B., Guiblet, W., Makova, K.D., and Nekrutenko, A. (2016). Streamlined analysis of duplex sequencing data with Du Novo. Genome Biol. 17, 180. 10.1186/s13059-016-1039-4.

30. Mikhailova, A.G., Mikhailova, A.A., Ushakova, K., Tretiakov, E.O., Iliushchenko, D., Shamansky, V., Lobanova, V., Kozenkov, I., Efimenko, B., Yurchenko, A.A., et al. (2022). A mitochondria-specific mutational signature of aging: increased rate of A > G substitutions on the heavy strand. Nucleic Acids Res. 50, 10264–10277. 10.1093/nar/gkac779.

31. Williams, S.L., Mash, D.C., Züchner, S., and Moraes, C.T. (2013). Somatic mtDNA mutation spectra in the aging human putamen. PLoS Genet. 9, e1003990. 10.1371/journal.pgen.1003990.

32. Li, M., Schröder, R., Ni, S., Madea, B., and Stoneking, M. (2015). Extensive tissue-related and allele-related mtDNA heteroplasmy suggests positive selection for somatic mutations. Proc. Natl. Acad. Sci. U. S. A. 112, 2491–2496. 10.1073/pnas.1419651112.

33. MITOMAP: A Human Mitochondrial Genome Database (2024). https://mitomap.org/.

34. Goto, Y., Nonaka, I., and Horai, S. (1990). A mutation in the tRNA(Leu)(UUR) gene associated with the MELAS subgroup of mitochondrial encephalomyopathies. Nature 348, 651–653. 10.1038/348651a0.

35. Yelverton, J.C., Dodson, K.M., Arnos, K., and Pandya, A. (2013). The clinical and audiologic features of hearing loss due to mitochondrial mutations: response to editor. Otolaryngol. Head Neck Surg. 149, 795–796. 10.1177/0194599813502927.

36. Yoneda, M., Chomyn, A., Martinuzzi, A., Hurko, O., and Attardi, G. (1992). Marked replicative advantage of human mtDNA carrying a point mutation that causes the MELAS encephalomyopathy. Proc. Natl. Acad. Sci. U. S. A. 89, 11164–11168. 10.1073/pnas.89.23.11164.

37. Millar, C.D., Dodd, A., Anderson, J., Gibb, G.C., Ritchie, P.A., Baroni, C., Woodhams, M.D., Hendy, M.D., and Lambert, D.M. (2008). Mutation and Evolutionary Rates in Adélie Penguins from the Antarctic. PLoS Genet. 4, e1000209. 10.1371/journal.pgen.1000209.

38. Barrett, A., Arbeithuber, B., Zaidi, A.A., Wilton, P., Paul, I.M., Nielsen, R., and Makova Kateryna, D. (2019). Pronounced somatic bottleneck in mitochondrial DNA of human hair. Philosophical Transactions B. 10.1098/rstb.2019-0175.

39. Hendy, M.D., Woodhams, M.D., and Dodd, A. (2009). Modelling mitochondrial site polymorphisms to infer the number of segregating units and mutation rate. Biol. Lett. 5, 397–400. 10.1098/rsbl.2009.0104.

40. Wei, W., Tuna, S., Keogh, M.J., Smith, K.R., Aitman, T.J., Beales, P.L., Bennett, D.L., Gale, D.P., Bitner-Glindzicz, M.A.K., Black, G.C., et al. (2019). Germline selection shapes human mitochondrial DNA diversity. Science 364. 10.1126/science.aau6520.

41. Floros, V.I., Pyle, A., Dietmann, S., Wei, W., Tang, W.C.W., Irie, N., Payne, B., Capalbo, A., Noli, L., Coxhead, J., et al. (2018). Segregation of mitochondrial DNA heteroplasmy through a developmental genetic bottleneck in human embryos. Nat. Cell Biol. 20, 144–151. 10.1038/s41556-017-0017-8.

42. Zhang, W., and Wu, F. (2023). Effects of adverse fertility-related factors on mitochondrial DNA in the oocyte: a comprehensive review. Reprod. Biol. Endocrinol. 21, 27. 10.1186/s12958-023-01078-6.

43. Tigges, J., Gordon, T.P., McClure, H.M., Hall, E.C., and Peters, A. (1988). Survival rate and life span of rhesus monkeys at the Yerkes regional primate research center. Am. J. Primatol. 15, 263–273. 10.1002/ajp.1350150308.

44. Alberts, S.C., Altmann, J., Brockman, D.K., Cords, M., Fedigan, L.M., Pusey, A., Stoinski, T.S., Strier, K.B., Morris, W.F., and Bronikowski, A.M. (2013). Reproductive aging patterns in primates reveal that humans are distinct. Proc. Natl. Acad. Sci. U. S. A. 110, 13440–13445. 10.1073/pnas.1311857110.

45. Marques, P., Madeira, T., and Gama, A. (2022). Menstrual cycle among adolescents: girls’ awareness and influence of age at menarche and overweight. Rev Paul Pediatr 40, e2020494. 10.1590/1984-0462/2022/40/2020494.

46. Goswami, D., and Conway, G.S. (2005). Premature ovarian failure. Hum. Reprod. Update 11, 391–410. 10.1093/humupd/dmi012.

47. Shen, Q., Liu, Y., Li, H., and Zhang, L. (2021). Effect of mitophagy in oocytes and granulosa cells on oocyte quality†. Biol. Reprod. 104, 294–304. 10.1093/biolre/ioaa194.

48. May-Panloup, P., Boucret, L., Chao de la Barca, J.-M., Desquiret-Dumas, V., Ferré-L’Hotellier, V., Morinière, C., Descamps, P., Procaccio, V., and Reynier, P. (2016). Ovarian ageing: the role of mitochondria in oocytes and follicles. Hum. Reprod. Update 22, 725–743. 10.1093/humupd/dmw028.

49. Yang, L., Lin, X., Tang, H., Fan, Y., Zeng, S., Jia, L., Li, Y., Shi, Y., He, S., Wang, H., et al. (2020). Mitochondrial DNA mutation exacerbates female reproductive aging via impairment of the NADH/NAD redox. Aging Cell 19, e13206. 10.1111/acel.13206.

50. Yao, Y.-G., Kajigaya, S., and Young, N.S. (2015). Mitochondrial DNA mutations in single human blood cells. Mutat. Res. 779, 68–77. 10.1016/j.mrfmmm.2015.06.009.

51. Guo, X., Xu, W., Zhang, W., Pan, C., Thalacker-Mercer, A.E., Zheng, H., and Gu, Z. (2023). High-frequency and functional mitochondrial DNA mutations at the single-cell level. Proc. Natl. Acad. Sci. U. S. A. 120, e2201518120. 10.1073/pnas.2201518120.

52. Sanchez-Contreras, M., Sweetwyne, M.T., Tsantilas, K.A., Whitson, J.A., Campbell, M.D., Kohrn, B.F., Kim, H.J., Hipp, M.J., Fredrickson, J., Nguyen, M.M., et al. (2023). The multi-tissue landscape of somatic mtDNA mutations indicates tissue-specific accumulation and removal in aging. Elife 12. 10.7554/eLife.83395.

53. Sanchez-Contreras, M., Sweetwyne, M.T., Kohrn, B.F., Tsantilas, K.A., Hipp, M.J., Schmidt, E.K., Fredrickson, J., Whitson, J.A., Campbell, M.D., Rabinovitch, P.S., et al. (2021). A replication-linked mutational gradient drives somatic mutation accumulation and influences germline polymorphisms and genome composition in mitochondrial DNA. Nucleic Acids Res. 49, 11103–11118. 10.1093/nar/gkab901.

54. An, J., Nam, C.H., Kim, R., Lee, Y., Won, H., Park, S., Lee, W.H., Park, H., Yoon, C.J., An, Y., et al. (2024). Mitochondrial DNA mosaicism in normal human somatic cells. Nat. Genet. 56, 1665–1677. 10.1038/s41588-024-01838-z.

55. Spelbrink, J.N., Toivonen, J.M., Hakkaart, G.A., Kurkela, J.M., Cooper, H.M., Lehtinen, S.K., Lecrenier, N., Back, J.W., Speijer, D., Foury, F., et al. (2000). In vivo functional analysis of the human mitochondrial DNA polymerase POLG expressed in cultured human cells. J. Biol. Chem. 275, 24818–24828. 10.1074/jbc.M000559200.

56. Zheng, W., Khrapko, K., Coller, H.A., Thilly, W.G., and Copeland, W.C. (2006). Origins of human mitochondrial point mutations as DNA polymerase gamma-mediated errors. Mutat. Res. 599, 11–20. 10.1016/j.mrfmmm.2005.12.012.

57. Ameur, A., Stewart, J.B., Freyer, C., Hagström, E., Ingman, M., Larsson, N.-G., and Gyllensten, U. (2011). Ultra-deep sequencing of mouse mitochondrial DNA: mutational patterns and their origins. PLoS Genet. 7, e1002028. 10.1371/journal.pgen.1002028.

58. Kreutzer, D.A., and Essigmann, J.M. (1998). Oxidized, deaminated cytosines are a source of C -> T transitions in vivo. Preprint, 10.1073/pnas.95.7.3578 https://doi.org/10.1073/pnas.95.7.3578.

59. Alseth, I., Dalhus, B., and Bjørås, M. (2014). Inosine in DNA and RNA. Preprint, 10.1016/j.gde.2014.07.008 https://doi.org/10.1016/j.gde.2014.07.008.

60. Falkenberg, M. (2018). Mitochondrial DNA replication in mammalian cells: overview of the pathway. Essays Biochem. 62, 287–296. 10.1042/EBC20170100.

61. Holt, I.J., Lorimer, H.E., and Jacobs, H.T. (2000). Coupled leading- and lagging-strand synthesis of mammalian mitochondrial DNA. Cell 100, 515–524. 10.1016/s0092-8674(00)80688-1.

62. Mechta, M., Ingerslev, L.R., Fabre, O., Picard, M., and Barrès, R. (2017). Evidence Suggesting Absence of Mitochondrial DNA Methylation. Front. Genet. 8, 166. 10.3389/fgene.2017.00166.

63. Bellizzi, D., D’Aquila, P., Scafone, T., Giordano, M., Riso, V., Riccio, A., and Passarino, G. (2013). The control region of mitochondrial DNA shows an unusual CpG and non-CpG methylation pattern. DNA Res. 20, 537–547. 10.1093/dnares/dst029.

64. Liu, B., Du, Q., Chen, L., Fu, G., Li, S., Fu, L., Zhang, X., Ma, C., and Bin, C. (2016). CpG methylation patterns of human mitochondrial DNA. Sci. Rep. 6, 23421. 10.1038/srep23421.

65. Fan, L.-H., Wang, Z.-B., Li, Q.-N., Meng, T.-G., Dong, M.-Z., Hou, Y., Ouyang, Y.-C., Schatten, H., and Sun, Q.-Y. (2019). Absence of mitochondrial DNA methylation in mouse oocyte maturation, aging and early embryo development. Biochem. Biophys. Res. Commun. 513, 912–918. 10.1016/j.bbrc.2019.04.100.

66. Patil, V., Cuenin, C., Chung, F., Rodriguez Aguilera, J.R., Fernandez-Jimenez, N., Romero-Garmendia, I., Bilbao, J.R., Cahais, V., Rothwell, J., and Herceg, Z. (2019). Human mitochondrial DNA is extensively methylated in a non-CpG context. Preprint, 10.1093/nar/gkz762 https://doi.org/10.1093/nar/gkz762.

67. López-Otín, C., Blasco, M.A., Partridge, L., Serrano, M., and Kroemer, G. (2013). The hallmarks of aging. Cell 153, 1194–1217. 10.1016/j.cell.2013.05.039.

68. Korotkevich, E., Conrad, D.N., Gartner, Z.J., and O’Farrell, P.H. (2024). Selection promotes age-dependent degeneration of the mitochondrial genome. bioRxiv. 10.1101/2024.09.27.615276.

69. Liu, F., Sun, X., Wei, C., Ji, L., Song, Y., Yang, C., Wang, Y., Liu, X., Wang, D., and Kang, J. (2024). Single-cell mitochondrial sequencing reveals low-frequency mitochondrial mutations in naturally aging mice. Aging Cell 23. 10.1111/acel.14242.

70. Li, M., Rothwell, R., Vermaat, M., Wachsmuth, M., Schröder, R., Laros, J.F.J., van Oven, M., de Bakker, P.I.W., Bovenberg, J.A., van Duijn, C.M., et al. (2016). Transmission of human mtDNA heteroplasmy in the Genome of the Netherlands families: support for a variable-size bottleneck. Genome Res. 26, 417–426. 10.1101/gr.203216.115.

71. Ebner, T., Moser, M., Shebl, O., Sommerguber, M., and Tews, G. (2008). Prognosis of oocytes showing aggregation of smooth endoplasmic reticulum. Reprod Biomed Online 16, 113–118. 10.1016/s1472-6483(10)60563-9.

72. Jayaprakash, A.D., Benson, E.K., Gone, S., Liang, R., Shim, J., Lambertini, L., Toloue, M.M., Wigler, M., Aaronson, S.A., and Sachidanandam, R. (2015). Stable heteroplasmy at the single-cell level is facilitated by intercellular exchange of mtDNA. Nucleic Acids Res. 43, 2177–2187. 10.1093/nar/gkv052.

73. Hazkani-Covo, E., Zeller, R.M., and Martin, W. (2010). Molecular poltergeists: mitochondrial DNA copies (numts) in sequenced nuclear genomes. PLoS Genet. 6, e1000834. 10.1371/journal.pgen.1000834.

74. Kennedy, S.R., Schmitt, M.W., Fox, E.J., Kohrn, B.F., Salk, J.J., Ahn, E.H., Prindle, M.J., Kuong, K.J., Shen, J.-C., Risques, R.-A., et al. (2014). Detecting ultralow-frequency mutations by Duplex Sequencing. Nat Protoc 9, 2586–2606. 10.1038/nprot.2014.170.

75. Schmitt, M.W., Fox, E.J., Prindle, M.J., Reid-Bayliss, K.S., True, L.D., Radich, J.P., and Loeb, L.A. (2015). Sequencing small genomic targets with high efficiency and extreme accuracy. Nat Methods 12, 423–425. 10.1038/nmeth.3351.

76. Galaxy Community (2024). The Galaxy platform for accessible, reproducible, and collaborative data analyses: 2024 update. Nucleic Acids Res 52, W83–W94. 10.1093/nar/gkae410.

77. Bolger, A.M., Lohse, M., and Usadel, B. (2014). Trimmomatic: a flexible trimmer for Illumina sequence data. Preprint, 10.1093/bioinformatics/btu170 https://doi.org/10.1093/bioinformatics/btu170.

78. Li, H., and Durbin, R. (2009). Fast and accurate short read alignment with Burrows-Wheeler transform. Bioinformatics 25, 1754–1760. 10.1093/bioinformatics/btp324.

79. Barnett, D.W., Garrison, E.K., Quinlan, A.R., Stromberg, M.P., and Marth, G.T. (2011). BamTools: a C API and toolkit for analyzing and managing BAM files. Preprint, 10.1093/bioinformatics/btr174 https://doi.org/10.1093/bioinformatics/btr174.

80. Jun, G., Wing, M.K., Abecasis, G.R., and Kang, H.M. (2015). An efficient and scalable analysis framework for variant extraction and refinement from population-scale DNA sequence data. Preprint, 10.1101/gr.176552.114 https://doi.org/10.1101/gr.176552.114.

81. Blankenberg, D., Von Kuster, G., Bouvier, E., Baker, D., Afgan, E., Stoler, N., Galaxy Team, Taylor, J., and Nekrutenko, A. (2014). Dissemination of scientific software with Galaxy ToolShed. Genome Biol. 15, 403. 10.1186/gb4161.

82. Nei, M., and Gojobori, T. (1986). Simple methods for estimating the numbers of synonymous and nonsynonymous nucleotide substitutions. Mol. Biol. Evol. 3, 418–426. 10.1093/oxfordjournals.molbev.a040410.

83. Benjamini, Y., and Hochberg, Y. (1995). Controlling the False Discovery Rate: A Practical and Powerful Approach to Multiple Testing. Preprint, 10.1111/j.2517-6161.1995.tb02031.x

