## Supplemental Figures for "Mitochondrial DNA mutations in human oocytes undergo frequency-dependent selection but do not increase with age"


**Supplementary figures** **2**

Figure S1. mtDNA enrichment efficiency, and sequencing depth. 2

Figure S2. Fixed differences in comparison to the human mtDNA reference sequence. 3

Figure S3. Distribution of high-confidence tissue-specific de novo mutations across tissues. 3

Figure S4. Mutations measured in multiple DCSs. 4

Figure S5. Strand bias. 5

Figure S6. Distribution of germline mutations across the mtDNA. 6

Figure S7. Distribution of somatic mutations across the mtDNA. 7

Figure S8. Disease-associated de novo mutations. 8

Figure S9. Correlation of MAFs between DCSs using Exonuclease V enrichment and from SSCS using targeted capture enrichment. 8

Figure S10. Heteroplasmic sites per donor depending on their age. 9

Figure S11. Correlation of heteroplasmy MAFs between somatic tissues and oocytes. 9

Figure S12. Correlation of high-confidence heteroplasmy MAFs between somatic tissues and oocytes. 10

Figure S13. Shifts of heteroplasmies with MAFs measured with high confidence. 11

### Supplementary figures

**Figure S1. mtDNA enrichment efficiency, and sequencing depth.**
(**A**) mtDNA enrichment efficiency in different sample types. (**B**) Mean duplex consensus sequence (DCS) sequencing depth for mtDNA. Asterisks indicate the median of the distribution. (**C**) The distribution of DCS sequencing depth across mtDNA for 28 randomly selected samples (12 oocytes, 9 blood, 7 saliva). The drop in sequencing depth around position 315 is caused by a polyC repeat (this region was excluded from mutation calling, see Methods).

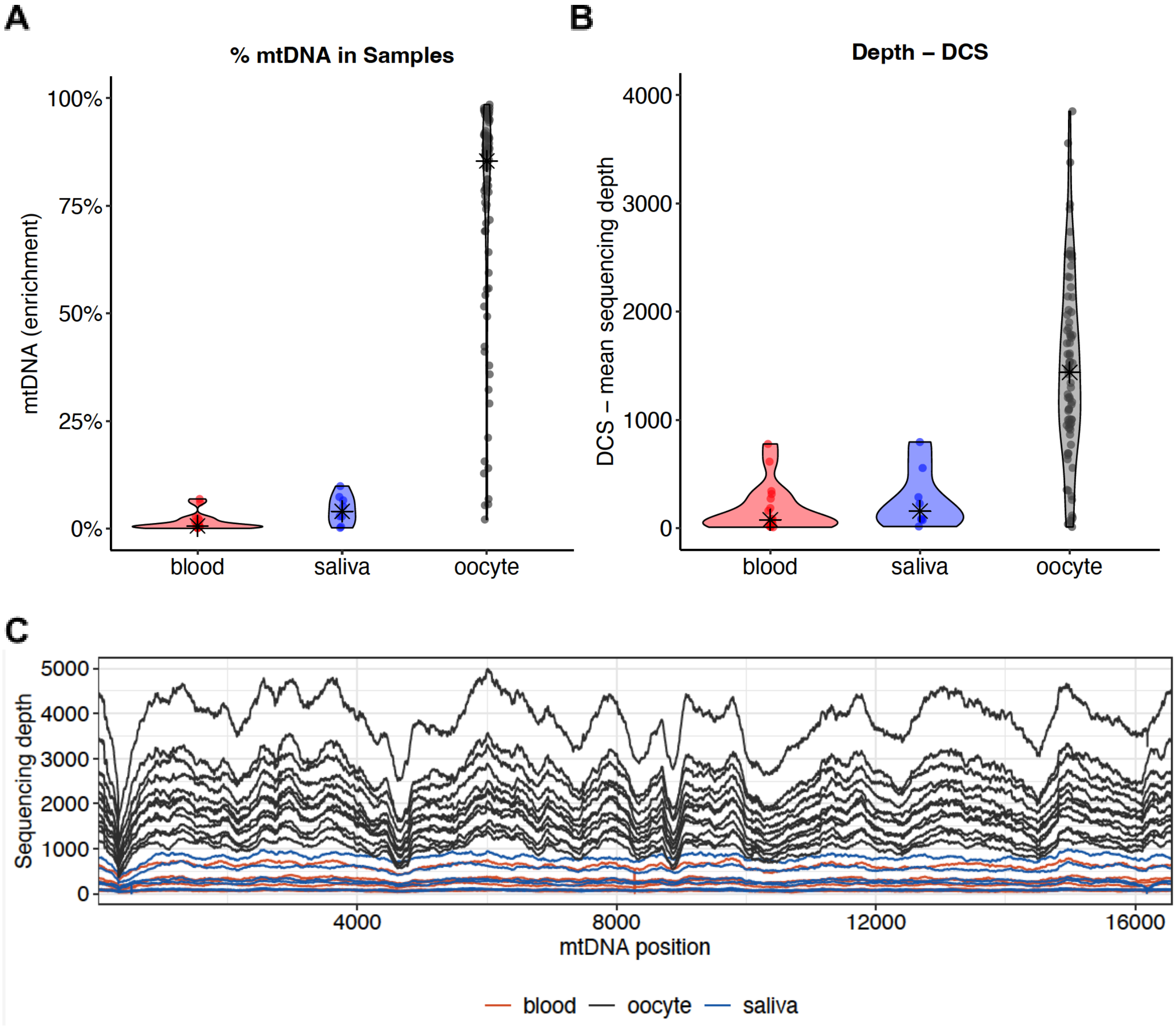

**Figure S2. Fixed differences in comparison to the human mtDNA reference sequence.**
Differences to the human reference sequence NC_012920.1 are shown. The analyzed women had 8-40 fixed differences per donor (520 in total; 219 different positions). Blue dashed lines indicate the boundaries of the D-loop region.

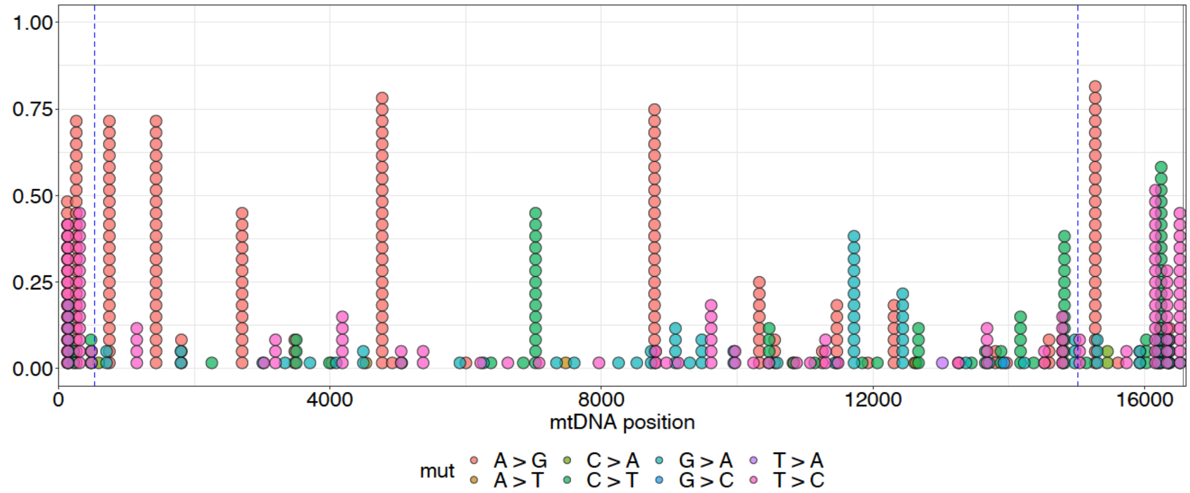


**Figure S3. Distribution of high-confidence tissue-specific *de novo* mutations across tissues.**
We identified a total of 3,525 high-confidence tissue-specific *de novo* mutations: 1,004 in blood, 644 in saliva, and 1,877 in oocytes. Mutations measured in multiple samples within a tissue type were only counted once.

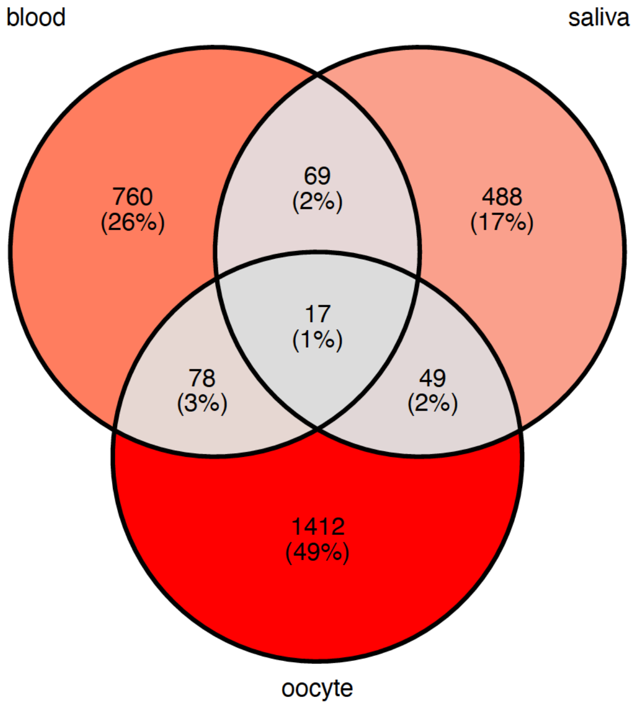

**Figure S4. Mutations measured in multiple DCSs.**

The distribution of high-frequency (MAF ≥1%) mutations across the mtDNA. Only mutations observed in at least 3 DCSs are shown (64 out of 247: 3 in blood, 2 in saliva, 59 in oocytes; 183 mutations occurred at sites with a low DCS depth and were found in one or two DCSs, therefore their frequency might have been overestimated and we excluded them from the subsequent analyses). Dot size corresponds to the number of molecules (i.e,. the number of DCSs) in which a mutation was observed. The orange dashed lines represent the boundaries of the D-loop region.

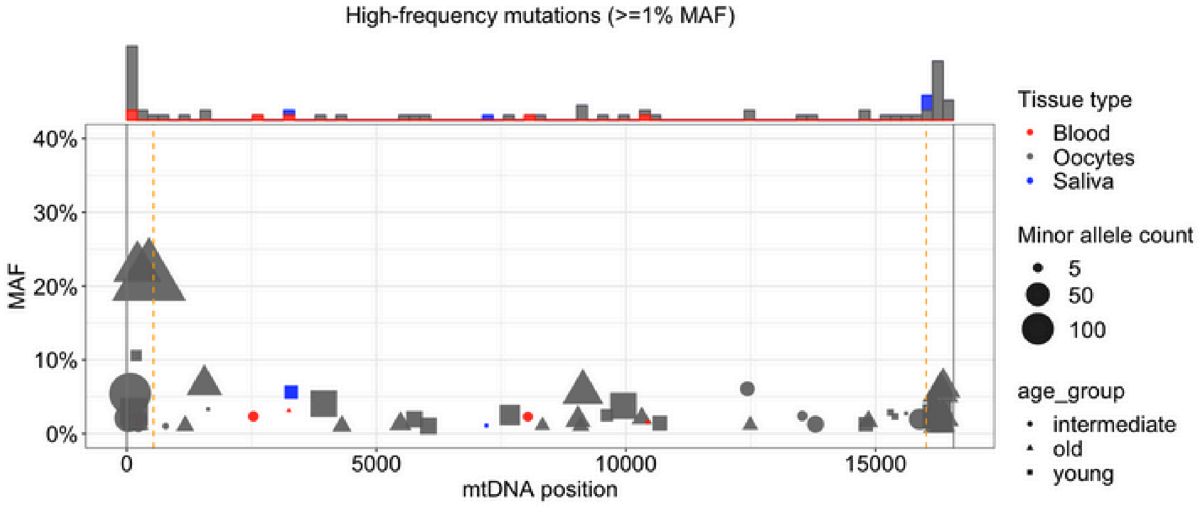

**Figure S5. Strand bias.**

(**A**) Different mutation types measured in blood and saliva, considering the whole mtDNA. (**B**) Different mutation types measured in oocytes, considering the whole mtDNA. (**C**) Different mutation types measured in blood and saliva, considering the D-loop only. (**D**) Different mutation types measured in oocytes, considering the D-loop only.

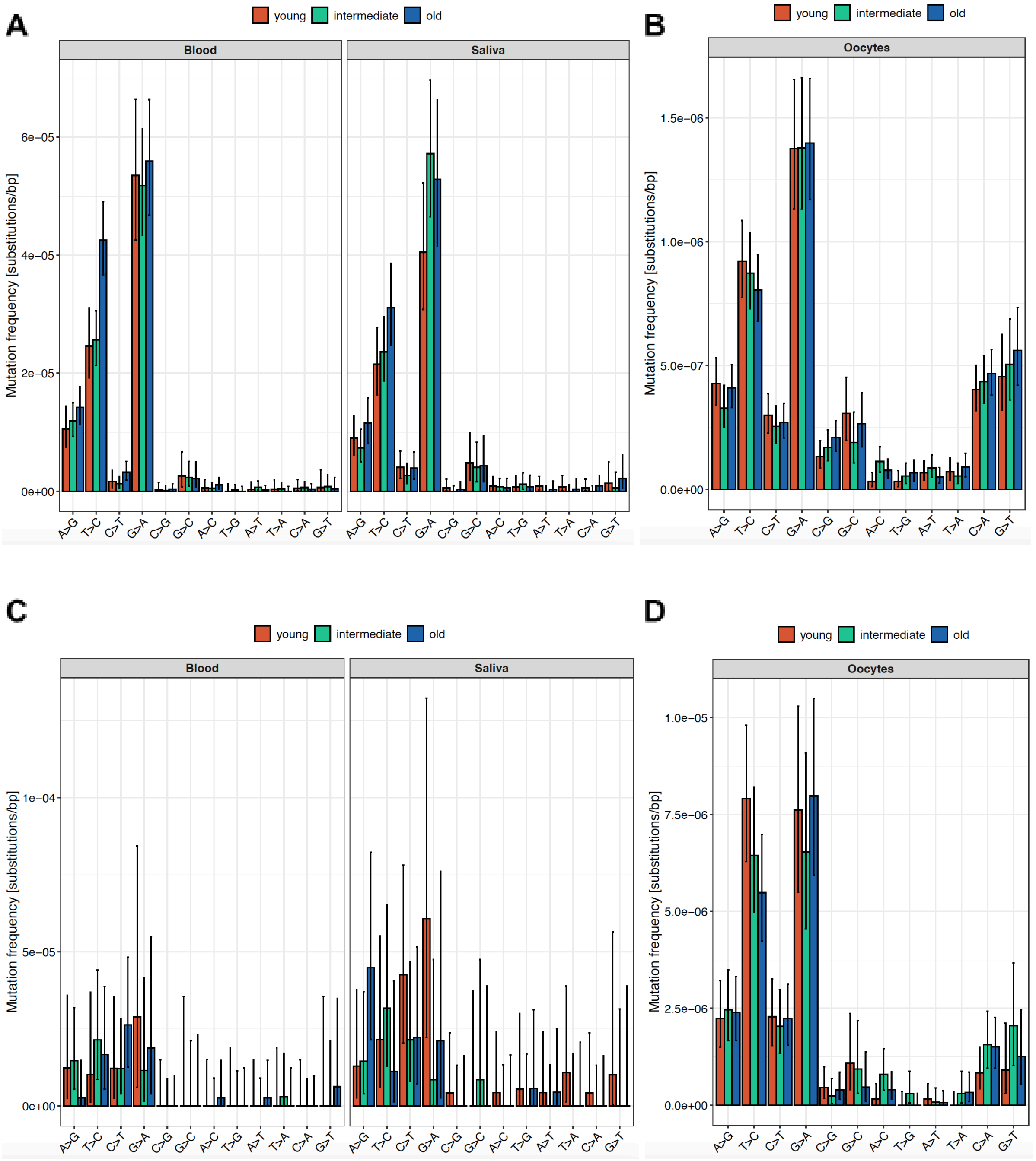

**Figure S6. Distribution of germline mutations across the mtDNA.**
Germline mutations, with their distribution displayed in 80 bins, each of 207 bp in size, in the upper part of each figure. Orange dashed lines indicate the boundaries of the D-loop region. Dot sizes correspond to the number of molecules (DCSs) in which the mutation was observed, separately shown for young, intermediate, and old women.

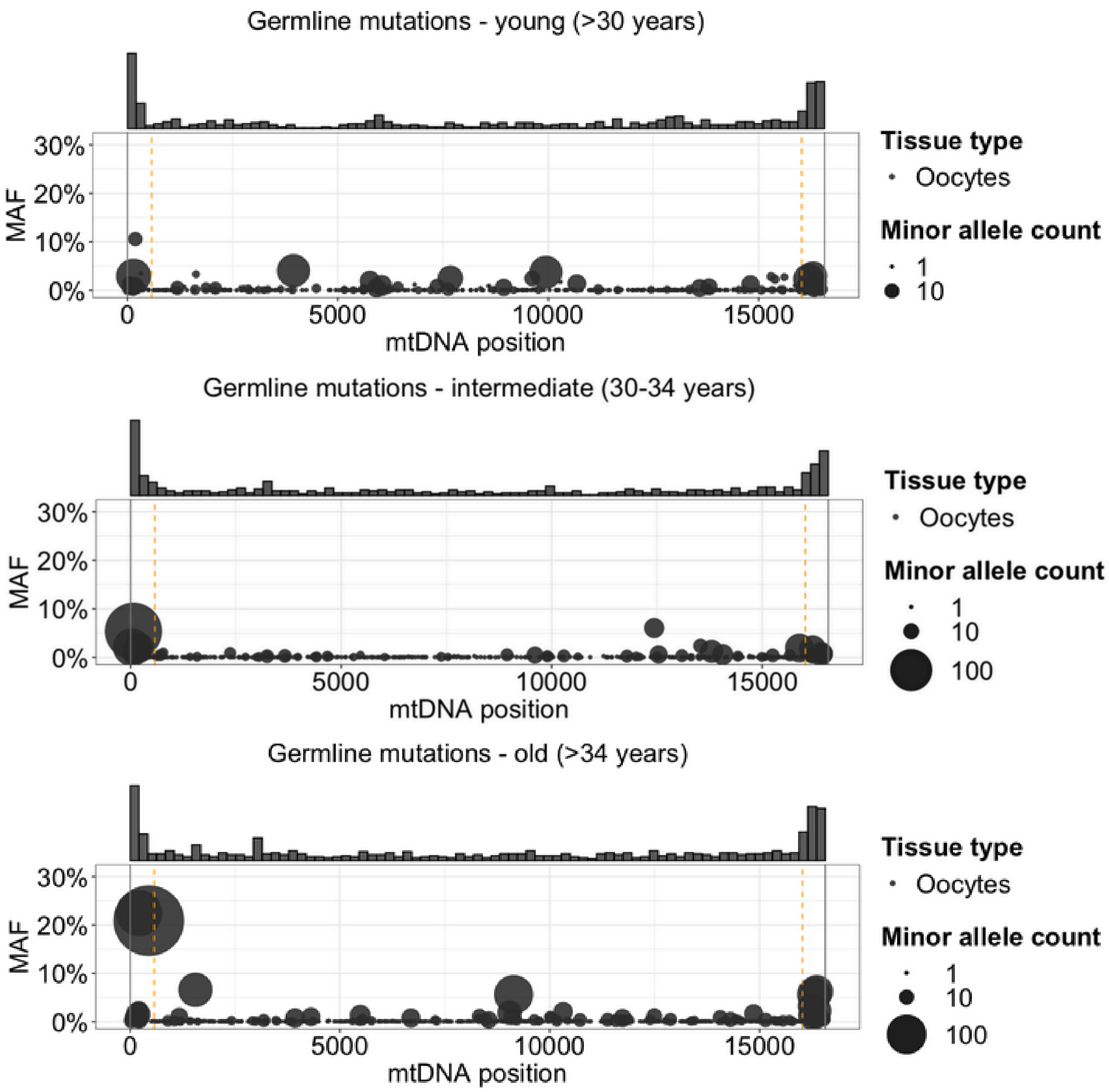

**Figure S7. Distribution of somatic mutations across the mtDNA.**
Somatic mutations, with their distribution displayed in 80 bins, each of 207 bp in size, in the upper part of each figure. Orange dashed lines indicate the boundaries of the D-loop region. Dot sizes correspond to the number of molecules (DCSs) in which the mutation was observed, separately shown for young, intermediate, and old women.

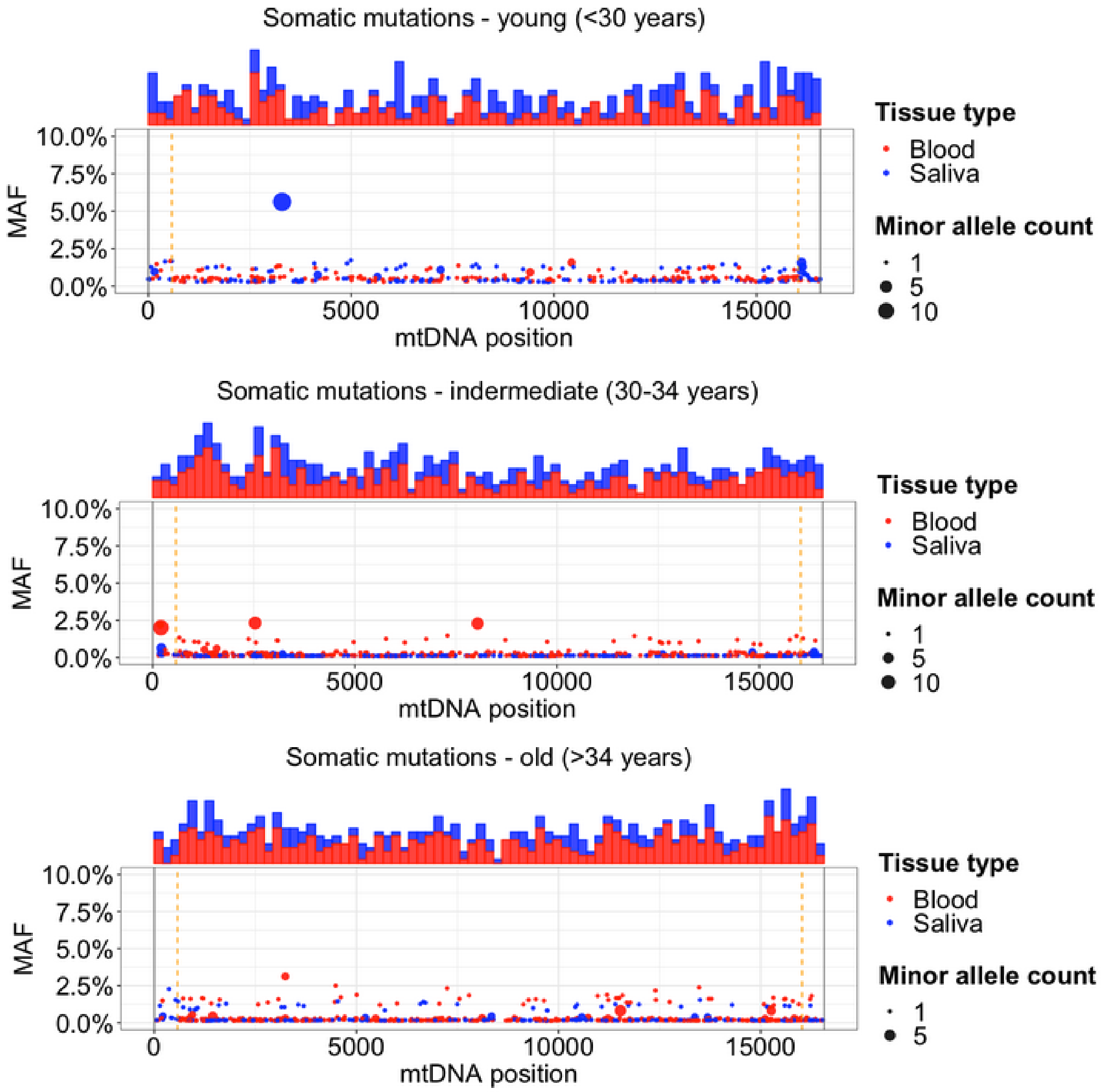

**Figure S8. Disease-associated *de novo* mutations.**

32 *de novo* mutations located at sites previously associated with diseases (14 in oocytes, 10 in blood, 8 in saliva); 109 disease-associated sites reported by MITOMAP ("MITOMAP: A Human Mitochondrial Genome Database" 2024), located in protein-coding regions, tRNA, and rRNA, were analyzed.

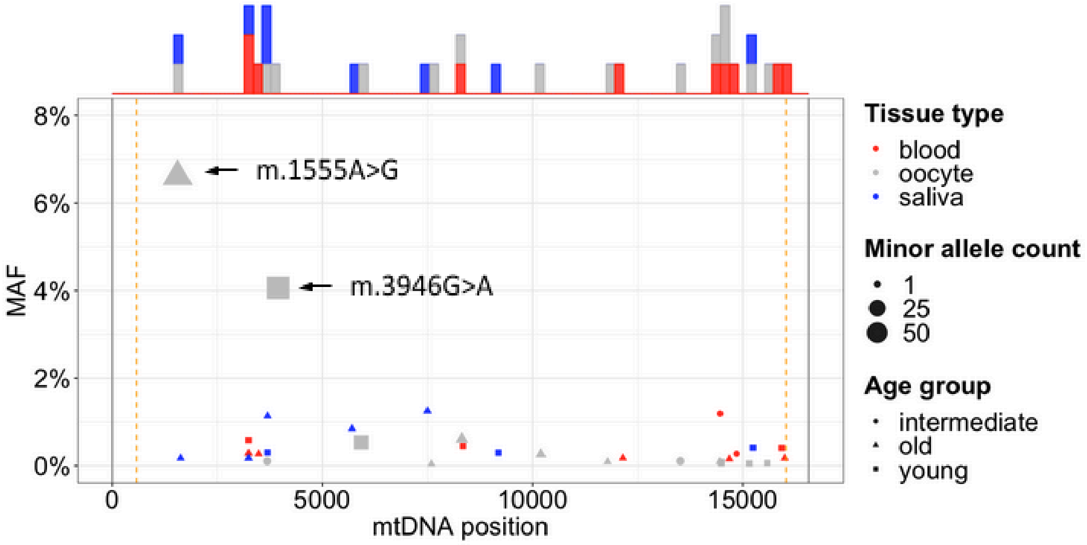


**Figure S9. Correlation of MAFs between DCSs using Exonuclease V enrichment and from SSCS using targeted capture enrichment.**

Heteroplasmies in blood (n=7) and saliva (n=6), which were sequenced using both mtDNA enrichment strategies (Exonuclease V and targeted capture), and for which the minor allele was measured in at least two DCSs, are shown.

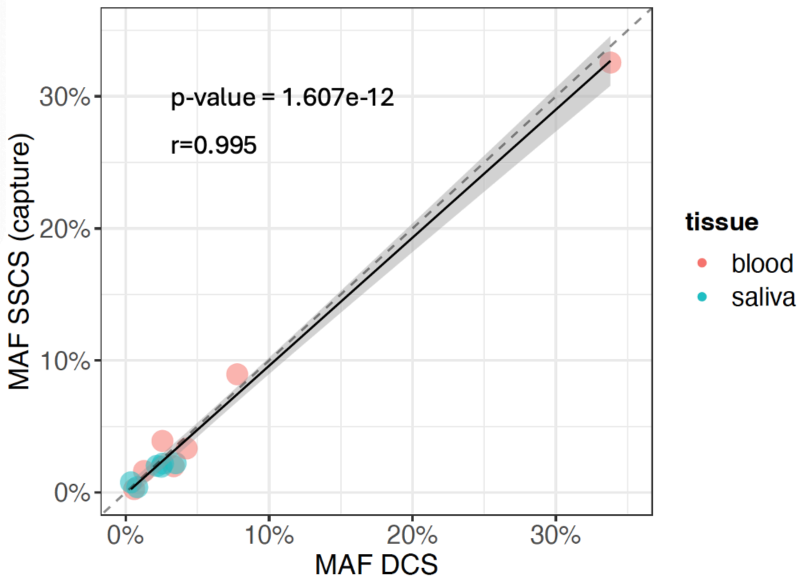

**Figure S10. Heteroplasmic sites per donor, depending on their age.**
Shown for the 22 assessed women. *P* value, r value, and confidence bounds for Pearson's correlation are shown.

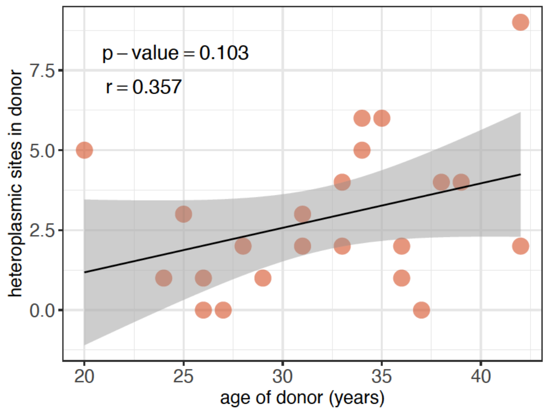


**Figure S11. Correlation of heteroplasmy MAFs between somatic tissues and oocytes.**
(**A**) Correlation of mean heteroplasmy frequencies in oocytes and somatic tissues, n=63 (if only one somatic tissue was available, the heteroplasmy frequency represents the available tissue). (**B**) Correlation of heteroplasmy frequencies in blood and saliva. Only heteroplasmies for which blood and saliva were assessed are included (n=18). (**C**) Normalized variance in somatic tissues in correlation with age. Only heteroplasmies for which blood and saliva were assessed are included (n=18). (**D**) Normalized variance in oocytes in correlation with age (n=63). *P* value, r value, and confidence bounds for Pearson's correlation are shown.

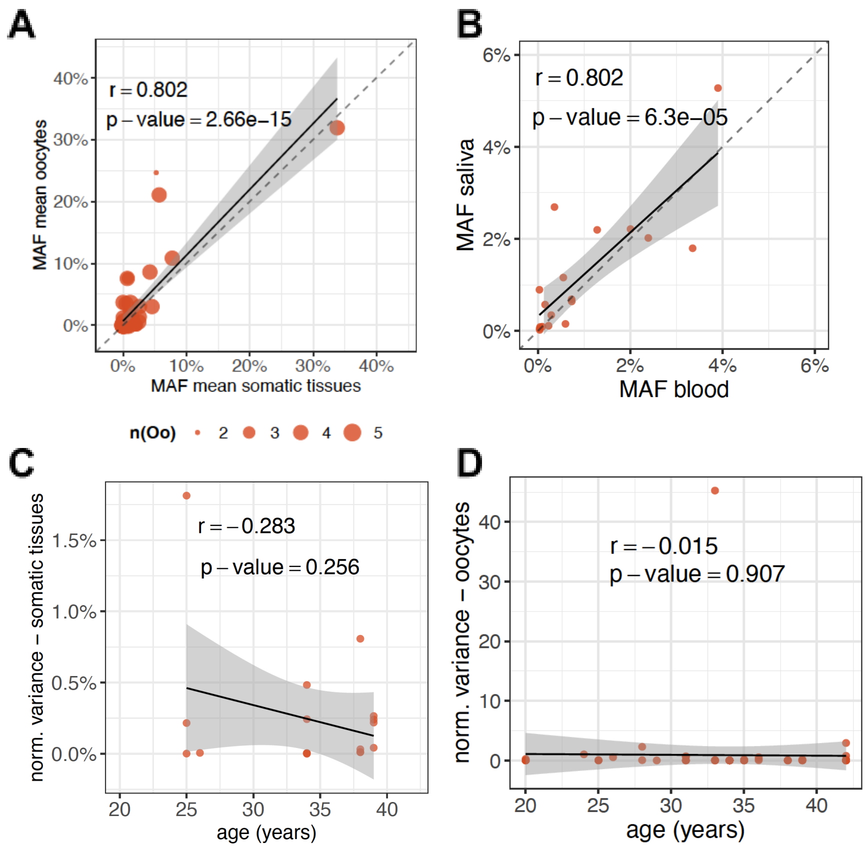

**Figure S12. Correlation of high-confidence heteroplasmy MAFs between somatic tissues and oocytes.** Only heteroplasmies measured in at least 3 consensus sequences were included to exclude potential bias from over- or underestimated MAFs. If this requirement was not fulfilled in somatic tissues, the whole heteroplasmic site was excluded; for oocytes, we excluded individual oocytes for which this requirement was not fulfilled. Oocytes with MAFs of 0% were only included if the sequencing depth at the corresponding position would allow the measurement of the MAF observed in somatic tissues in at least 3 consensus sequences. (**A**) Correlation of mean heteroplasmy frequencies in oocytes and somatic tissues, n=30 (if only one somatic tissue was available, the heteroplasmy frequency represents the available tissue). (**B**) Correlation of heteroplasmy frequencies in blood and saliva. Only heteroplasmies for which blood and saliva were assessed are included (n=10). (**C**) Normalized variance in somatic tissues in correlation with age. Only heteroplasmies for which blood and saliva were assessed are included (n=10). (**D**) Normalized variance in oocytes in correlation with age (n=30). *P* value, r value, and confidence bounds for Pearson's correlation are shown.

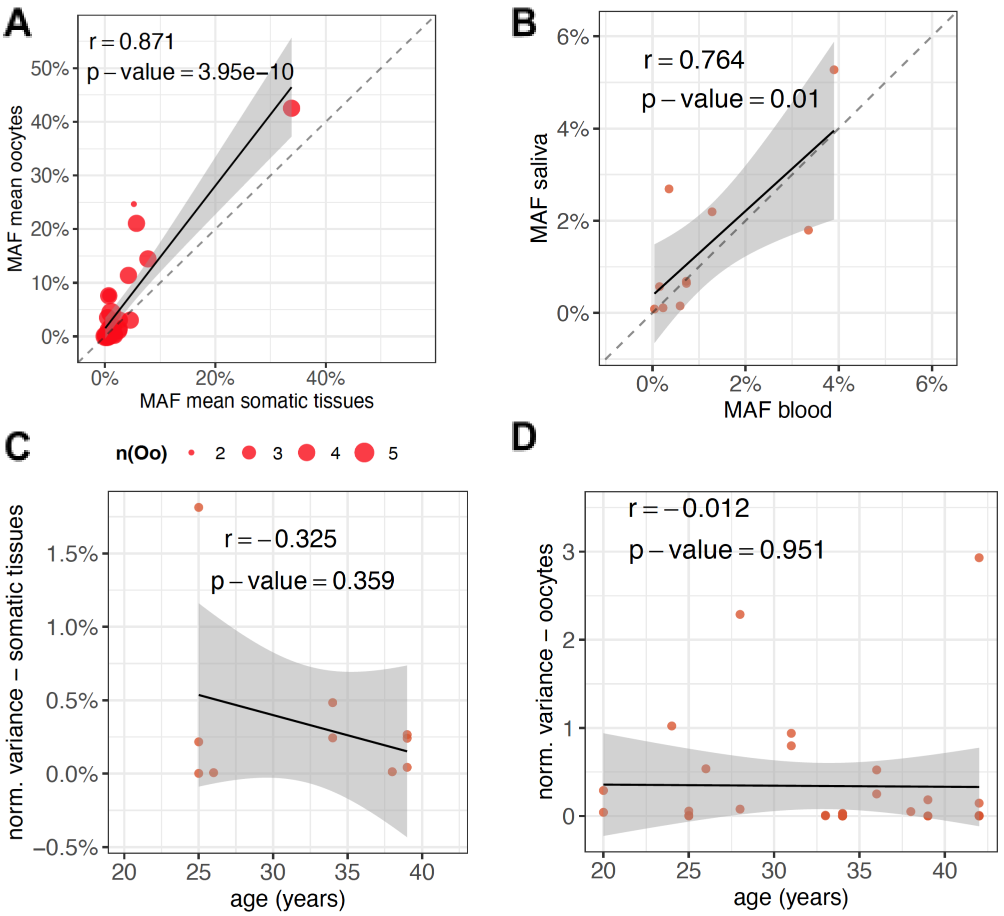

**Figure S13. Shifts of heteroplasmies with MAFs measured with high confidence.**
(**A**) MAF difference between oocytes and somatic tissues measured in percentage points, with a positive shift representing a higher MAF in oocytes compared to somatic tissues and a negative shift representing a lower MAF in oocytes compared to somatic tissue. Colors indicate their localization in different mtDNA regions, and shapes show the age group of each woman. (**B**) The number of heteroplasmies with an increased MAF in oocytes (positive shift) or decreased MAF (negative shift) is shown separately for different mtDNA regions. 29, 6, 20, and 3 of the heteroplasmy transmissions had a positive change; 82, 13, 59 and 10 of the heteroplasmy transmission had a negative change ($p$=9.06×10$^{-8}$, $p$=0.209, $p$=6.48×10$^{-5}$, and $p$=0.154, binomial test) considering all heteroplasmies, heteroplasmies in protein-coding regions, heteroplasmies in the D-loop, and heteroplasmies in rRNA, respectively. Left-pointing arrows indicate higher numbers of negative versus positive shifts, and right-pointing arrows indicate higher numbers of positive versus negative shifts. *$p$<0.05, **p<0.01, ***$p$<0.001, and ****$p$<0.0001; binomial test, corrected for multiple testing using the method of Benjamini-Hochberg to control the false discovery rate.

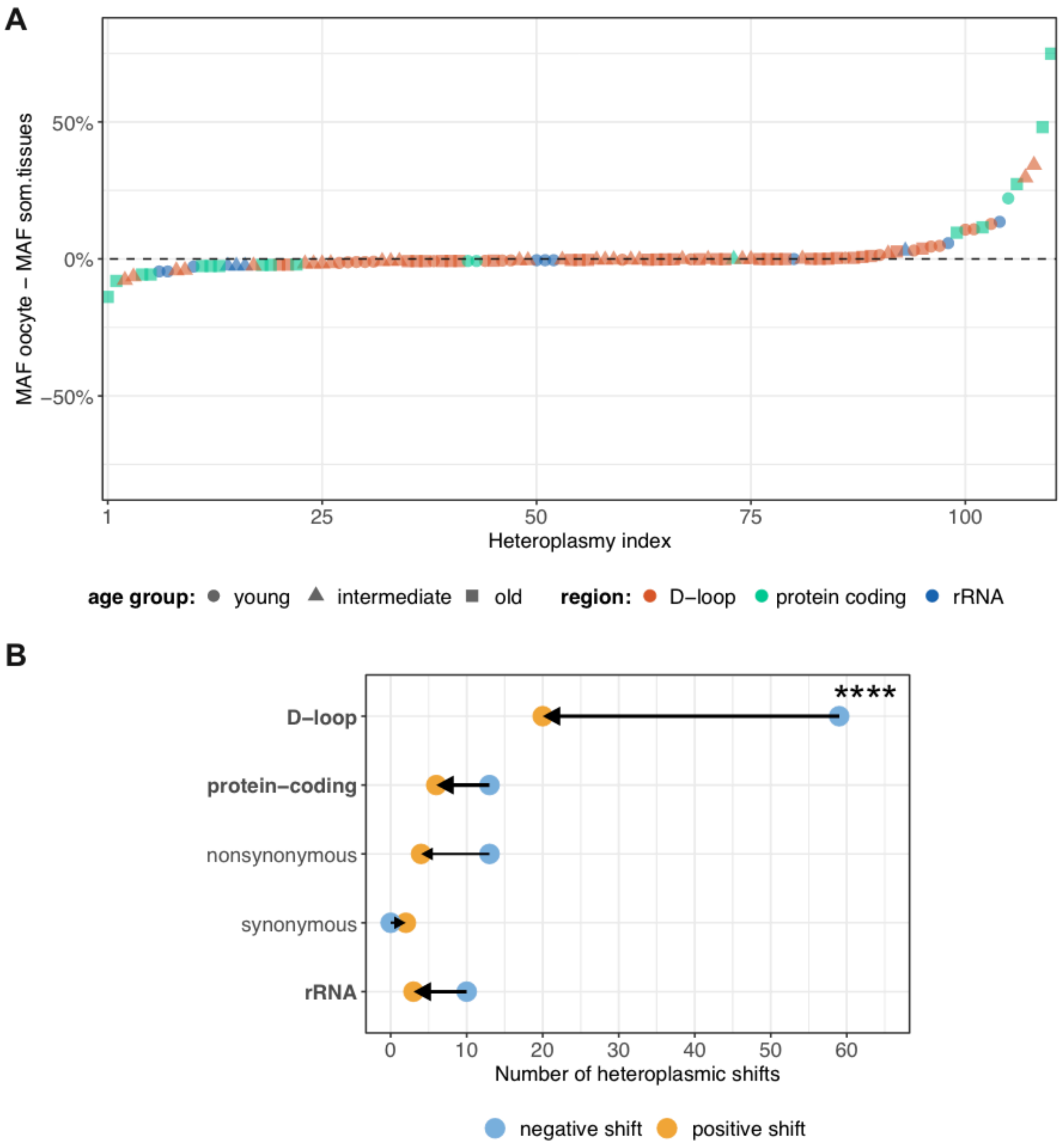
