## Supplemental Tables for "Mitochondrial DNA mutations in human oocytes undergo frequency-dependent selection but do not increase with age"

**Table S1. Samples analyzed with Duplex sequencing.** 22 women were included in the study, ranging in age from 20-42 years. Haplogroups were determined using HaploGrep 2.0.

| donor ID | age (years) | age group | origin | Haplogroup | Somatic tissue (s) | N sequenced oocytes | N heteroplasmic sites |
|---|---|---|---|---|---|---|---|
| hs001 | 27 | young | Macedonia | H47a | blood | 1 | 0 |
| hs002 | 34 | intermediate | Austria | H16 | blood | 4 | 6 |
| hs003 | 29 | young | Philippines | E1a1a1a | saliva | 5 | 1 |
| hs004 | 39 | old | Austria | H13a1a6 | blood | 2 | 4 |
| hs005 | 24 | young | Egypt | X1c | saliva | 4 | 1 |
| hs006 | 25 | young | Austria | U5a2a1+152 | blood, saliva* | 4 | 3 |
| hs007 | 36 | old | Austria | K1a4a1 | blood | 4 | 1 |
| hs008 | 26 | young | Hungary | H+195 | blood, saliva | 2 | 0 |
| hs009 | 37 | old | Romania | H44a | blood | 4 | 0 |
| hs010 | 26 | young | Austria | HV0 | blood, saliva | 4 | 1 |
| hs011 | 38 | old | Austria | T1a1 | blood*, saliva | 3 | 4 |
| hs012 | 42 | old | Austria | HV10 | saliva | 4 | 9 |
| hs014 | 34 | intermediate | Austria | U5a1c2a1 | blood* | 5 | 5 |
| hs015 | 42 | old | China | B4c1b2c2 | blood* | 4 | 2 |
| hs016 | 33 | intermediate | Austria | K1a+195 | blood* | 2 + 2* | 2 |
| hs017 | 20 | young | Austria | U5a1c1 | blood* | 5 | 5 |
| hs018 | 36 | old | Austria | H16 | blood* | 2 | 2 |
| hs019 | 35 | old | Austria | J1c2t | blood | 4 | 6 |
| hs020 | 28 | young | Austria | H4a1 | blood* | 3 | 2 |
| hs021 | 31 | intermediate | Hungary | H7g | blood | 4 | 2 |
| hs022 | 31 | intermediate | Austria | I4a | blood | 4 | 3 |
| hs023 | 33 | intermediate | Austria | K1b1a1 | saliva | 4 | 4 |

* mean DCS (duplex consensus sequence) depth is <50x

**Table S2. Median duplex consensus sequence (DCS) depths and mtDNA enrichment in the different tissues and age groups.**

| tissue | N samples | median DCS depth | min. DCS depth | max. DCS depth | median mtDNA enrichment | min. mtDNA enrichment | max. mtDNA enrichment |
|---|---|---|---|---|---|---|---|
| blood (all) | 18 | 78 | 9 | 778 | 0.65% | 0.08% | 6.91% |
| saliva (all) | 8 | 158 | 16 | 796 | 4.02% | 0.20% | 9.89% |
| oocytes (all) | 80 | 1440 | 12 | 3850 | 85.39% | 2.13% | 98.50% |
| blood (DCS-depth >50x) | 11 | 186 | 57 | 778 | 1.60% | 0.46% | 6.91% |
| saliva (DCS-depth >50x) | 7 | 228 | 75 | 796 | 5.16% | 0.46% | 9.89% |
| oocytes (DCS-depth >50x) | 78 | 1464 | 68 | 3850 | 86.13% | 2.13% | 98.50% |

**Table S3. Numbers of mutations identified in different tissues and age groups.** One-sided permutation test *p*-values were corrected for multiple testing using the method of Benjamini-Hochberg to control the false discovery rate. The permutations test could not be sucessfully performed for comparisons with saliva samples of the intermediate age group, as the measurement is based on a single donor, hence referred to as NA.

| tissue | number of mutations (all age groups) | number of mutations (young) | number of mutations (intermediate) | number of mutations (old) | median mut_freq (young) | median mut_freq (intermediate) | median mut_freq (old) | fold- difference (old/young) | fold- difference (intermediate/ young) | fold- difference (old/intermediate) | p-value (young - old) | p-value (young - intermediate) | p-value (intermediate - old) |
|---|---|---|---|---|---|---|---|---|---|---|---|---|---|
| blood | 1,004 | 209 | 360 | 435 | 1.82E-05 | 1.86E-05 | 2.44E-05 | 1.34 | 1.02 | 1.31 | 2.96E-02 | 0.5345 | 0.0845 |
| saliva | 644 | 186 | 232 | 226 | 1.76E-05 | 1.76E-05 | 2.32E-05 | 1.32 | 1.00 | 1.32 | 6.66E-02 | NA | NA |
| oocytes | 1,877 | 594 | 570 | 713 | 1.03E-06 | 9.52E-07 | 1.02E-06 | 0.99 | 0.92 | 1.08 | 0.5499 | 0.7569 | 0.3935 |

**Table S4. Mixed-effects logistic regression model.** Fixed effects: donor age (scaled), tissue (blood; saliva; oocyte). Random effects: donor ID (multiple observations correspond to the same woman, 22 women in total). Response: mutation frequency with weights given by the number of sequenced nucleotides corresponding to each sample. Oocytes with a mean DCS depth <50× where not considered. Marginal pseudo-R2 (represents the variance explained by the fixed effects): 29.21%. Conditional pseudo-R2 (represents the variance explained by both fixed and random effects): 29.30%.

**Random effects:**

| Groups | Name | Variance | Std.Dev. | Chi-square test p-value |
|---|---|---|---|---|
| donor ID | Intercept | 0.0043 | 0.0657 | 0.0635 |

Number of obs: 96, groups: donor_ID, 22

Conditional modes of random effects:

| Min. | 1st Qu. | Median | Mean | 3rd Qu. | Max. |
|---|---|---|---|---|---|
| -0.0815 | -0.0319 | 0.0062 | 0.0005 | 0.0216 | 0.0755 |

**Fixed effects:**

| | Estimate | Std. error | z value | Pr(>\|z\|) | Significance code |
|---|---|---|---|---|---|
| Intercept | -10.843 | 0.040 | -269.717 | < 2.00E-16 | *** |
| saliva | -0.057 | 0.064 | -0.898 | 0.369 | |
| oocyte | -3.000 | 0.045 | -66.900 | < 2.00E-16 | *** |
| scaled age | 0.152 | 0.051 | 3.005 | 0.003 | ** |
| scaled age * saliva | -0.067 | 0.066 | -1.020 | 0.308 | |
| scaled age * oocyt | -0.141 | 0.055 | -2.574 | 0.010 | * |

Signif. codes: 0 ‘***’ 0.001 ‘**’ 0.01 ‘*’ 0.05 ‘.’ 0.1 ‘ ’ 1

**Fixed effects with age in years (not scaled) and all tissues made explicit**

| | Estimate | Z test p-value |
|---|---|---|
| slope for blood | 0.026 | 2.65E-03 |
| slope for saliva | 0.014 | 5.67E-02 |
| slope for oocyte | 0.002 | 0.6876 |

**Table S5. Mutation frequencies in different regions of mtDNA.** nt_region refers to the number of nucleotides sequenced for a specific tissue, age group, and region. nt_mut refers to the number of mutations measured in a specific tissue, age group, and region. The D-loop (DL; 1,122 bp) and intergenic DNA (i.e. non-coding regions outside the DL; 88 bp) are listed separately to allow for better comparison to reported results on the DL region solely. Significance in differences in the mutation frequencies were tested for the different regions and tissues for adjacent age groups as well as between young and old women. Additionally, significance in differences were tested for mutation frequencies in the DL compared to coding region (i.e. protein-coding (11,341 bp), rRNA (2,513 bp), and tRNA (1,505 bp)). The Fisher's Exact test (FET) p-values were corrected for multiple testing using the method of Benjamini-Hochberg to control the false discovery rate.

| region | tissue | n_mut (young) | nt_region (young) | n_mut (int.) | nt_region (int.) | n_mut (old) | nt_region (old) | mut_freq (young) | mut_freq (int.) | mut_freq (old) | FET (young-old) | fold-change (old/young) | FET (young-int.) | fold-change (int./young) | FET (int.-old) | fold-change (old/int.) | FET (DL-nDL) (young) | FET (DL-nDL) (int) | FET (DL-nDL) (old) |
|---|---|---|---|---|---|---|---|---|---|---|---|---|---|---|---|---|---|---|---|
| total mtDNA | blood | 209 | 11,681,145 | 360 | 19,567,989 | 435 | 17,977,365 | 1.79E-05 | 1.84E-05 | 2.42E-05 | **2.96E-03** | 1.35 | 1.000 | 1.03 | **8.65E-04** | 1.32 | | | |
| total mtDNA | oocyte | 594 | 620,194,239 | 570 | 602,713,944 | 713 | 719,028,324 | 9.58E-07 | 9.46E-07 | 9.92E-07 | 0.779 | 1.04 | 1.000 | 0.99 | 0.647 | 1.05 | | | |
| total mtDNA | saliva | 186 | 11,101,230 | 232 | 13,188,924 | 226 | 10,670,436 | 1.68E-05 | 1.76E-05 | 2.12E-05 | 0.054 | 1.26 | 1.000 | 1.05 | 0.203 | 1.20 | | | |
| D-loop | blood | 11 | 791,010 | 21 | 1,325,082 | 22 | 1,217,370 | 1.39E-05 | 1.58E-05 | 1.81E-05 | 0.779 | 1.30 | 1.000 | 1.14 | 0.841 | 1.15 | 0.552 | 0.598 | 0.271 |
| D-loop | oocyte | 215 | 41,997,582 | 213 | 40,813,872 | 238 | 48,690,312 | 5.12E-06 | 5.22E-06 | 4.89E-06 | 0.779 | 0.96 | 1.000 | 1.02 | 0.695 | 0.94 | **8.24E-96** | **2.01E-97** | **6.91E-97** |
| D-loop | saliva | 31 | 751,740 | 19 | 893,112 | 21 | 722,568 | 4.12E-05 | 2.13E-05 | 2.91E-05 | 0.467 | 0.71 | 0.483 | 0.52 | 0.600 | 1.37 | **1.26E-05** | 0.462 | 0.261 |
| coding regions | blood | 198 | 10,828,095 | 336 | 18,138,979 | 412 | 16,664,515 | 1.83E-05 | 1.85E-05 | 2.47E-05 | **2.96E-03** | 1.35 | 1.000 | 1.01 | **8.65E-04** | 1.34 | | | |
| coding regions | oocyte | 376 | 574,902,729 | 355 | 558,698,984 | 473 | 666,519,164 | 6.54E-07 | 6.35E-07 | 7.10E-07 | 0.467 | 1.09 | 1.000 | 0.97 | 0.350 | 1.12 | | | |
| coding regions | saliva | 155 | 10,290,530 | 211 | 12,225,764 | 205 | 9,891,196 | 1.51E-05 | 1.73E-05 | 2.07E-05 | **1.39E-02** | 1.37 | 1.000 | 1.15 | 0.238 | 1.20 | | | |
| intergenic | blood | 0 | 62,040 | 3 | 103,928 | 1 | 95,480 | 0.00E+00 | 2.89E-05 | 1.05E-05 | 1.000 | NA | 1.000 | NA | 0.731 | 0.36 | | | |
| intergenic | oocyte | 3 | 3,293,928 | 2 | 3,201,088 | 2 | 3,818,848 | 9.11E-07 | 6.25E-07 | 5.24E-07 | 0.779 | 0.58 | 1.000 | 0.69 | 1.000 | 0.84 | | | |
| intergenic | saliva | 0 | 58,960 | 2 | 70,048 | 0 | 56,672 | 0.00E+00 | 2.86E-05 | 5.29E-05 | 1.000 | NA | 1.000 | NA | 0.695 | 1.85 | | | |
| protein-coding | blood | 131 | 7,995,405 | 218 | 13,393,721 | 294 | 12,304,985 | 1.64E-05 | 1.63E-05 | 2.39E-05 | **2.96E-03** | 1.46 | 1.000 | 0.99 | **3.56E-04** | 1.47 | | | |
| protein-coding | oocyte | 273 | 424,504,971 | 254 | 412,540,216 | 320 | 492,154,036 | 6.43E-07 | 6.16E-07 | 6.50E-07 | 0.997 | 1.01 | 1.000 | 0.96 | 0.695 | 1.06 | | | |
| protein-coding | saliva | 123 | 7,598,470 | 150 | 9,027,436 | 138 | 7,303,604 | 1.62E-05 | 1.66E-05 | 1.89E-05 | 0.467 | 1.17 | 1.000 | 1.02 | 0.600 | 1.14 | | | |
| rRNA | blood | 47 | 1,771,665 | 87 | 2,967,853 | 81 | 2,726,605 | 2.65E-05 | 2.93E-05 | 2.97E-05 | 0.779 | 1.12 | 1.000 | 1.11 | 0.986 | 1.01 | | | |
| rRNA | oocyte | 67 | 94,064,103 | 65 | 91,412,888 | 99 | 109,054,148 | 7.12E-07 | 7.11E-07 | 9.08E-07 | 0.365 | 1.28 | 1.000 | 1.00 | 0.358 | 1.28 | | | |
| rRNA | saliva | 23 | 1,683,710 | 45 | 2,000,348 | 41 | 1,618,372 | 1.37E-05 | 2.25E-05 | 2.53E-05 | 0.054 | 1.85 | 0.544 | 1.64 | 0.728 | 1.12 | | | |
| tRNA | blood | 20 | 1,061,025 | 31 | 1,777,405 | 37 | 1,632,925 | 1.88E-05 | 1.74E-05 | 2.27E-05 | 0.779 | 1.21 | 1.000 | 0.93 | 0.600 | 1.30 | | | |
| tRNA | oocyte | 36 | 56,333,655 | 36 | 54,745,880 | 54 | 65,310,980 | 6.39E-07 | 6.58E-07 | 8.27E-07 | 0.467 | 1.29 | 1.000 | 1.03 | 0.600 | 1.26 | | | |
| tRNA | saliva | 9 | 1,008,350 | 16 | 1,197,980 | 26 | 969,220 | 8.93E-06 | 1.34E-05 | 2.68E-05 | **1.46E-02** | 3.00 | 1.000 | 1.50 | 0.154 | 2.00 | | | |

DL ... D-loop

nDL ... outside D-loop

**Table S6. Transition and transversion mutations in the different tissues, age groups, and regions.** Ti: transitions, Tv: transversions.

| ***Young:*** | all regions | | | D-loop | | | outside the D-loop | | |
|---|---|---|---|---|---|---|---|---|---|
| **tissue** | **n_Ti** | **n_Tv** | **Ti/Tv** | **n_Ti** | **n_Tv** | **Ti/Tv** | **n_Ti** | **n_Tv** | **Ti/Tv** |
| blood | 197 | 12 | 16.42 | 11 | 0 | NA | 186 | 12 | 15.50 |
| saliva | 163 | 23 | 7.09 | 23 | 8 | 2.88 | 140 | 15 | 9.33 |
| oocyte | 393 | 201 | 1.96 | 183 | 32 | 5.72 | 210 | 169 | 1.24 |

| ***Intermediate:*** | all regions | | | D-loop | | | outside the D-loop | | |
|---|---|---|---|---|---|---|---|---|---|
| **tissue** | **n_Ti** | **n_Tv** | **Ti/Tv** | **n_Ti** | **n_Tv** | **Ti/Tv** | **n_Ti** | **n_Tv** | **Ti/Tv** |
| blood | 337 | 23 | 14.65 | 20 | 1 | 20.00 | 317 | 22 | 14.41 |
| saliva | 217 | 15 | 14.47 | 18 | 1 | 18.00 | 199 | 14 | 14.21 |
| oocyte | 348 | 222 | 1.57 | 157 | 56 | 2.80 | 191 | 166 | 1.15 |

| ***Old:*** | all regions | | | D-loop | | | outside the D-loop | | |
|---|---|---|---|---|---|---|---|---|---|
| **tissue** | **n_Ti** | **n_Tv** | **Ti/Tv** | **n_Ti** | **n_Tv** | **Ti/Tv** | **n_Ti** | **n_Tv** | **Ti/Tv** |
| blood | 418 | 17 | 24.59 | 19 | 3 | 6.33 | 399 | 14 | 28.50 |
| saliva | 207 | 19 | 10.89 | 19 | 2 | 9.50 | 188 | 17 | 11.06 |
| oocyte | 427 | 286 | 1.49 | 187 | 51 | 3.67 | 240 | 235 | 1.02 |

**Table S7. Frequencies of different mutation types in somatic tissues and oocytes.** 'n_mut' refers to the number of mutations measured in a specific tissue, age group, and region.'nt_type' is the number of sequenced nucleotides that can undergo the particular mutation type (e.g. all Cs for C>T mutations). 'int.' - intermediate age group. The Fisher's Exact test (FET) *p*-values were corrected for multiple testing using the method of Benjamini-Hochberg to control the false discovery rate.

| type | tissue | n_mut (young) | nt_type (young) | n_mut (int.) | nt_type (int.) | n_mut (old) | nt_type (old) | mut_freq (young) | mut_freq (int.) | mut_freq (old) | FET (young-old) | FET (young-int.) | FET (int.-old) |
|---|---|---|---|---|---|---|---|---|---|---|---|---|---|
| A>C/T>G | blood | 2 | 6,493,755 | 4 | 10,878,191 | 6 | 9,993,935 | 3.08E-07 | 3.68E-07 | 6.00E-07 | 0.849 | 1.000 | 0.875 |
| A>G/T>C | blood | 109 | 6,493,755 | 196 | 10,878,191 | 268 | 9,993,935 | 1.68E-05 | 1.80E-05 | 2.68E-05 | **4.26E-04** | 1.000 | **4.02E-04** |
| A>T/T>A | blood | 2 | 6,493,755 | 6 | 10,878,191 | 1 | 9,993,935 | 3.08E-07 | 5.52E-07 | 1.00E-07 | 0.849 | 1.000 | 0.561 |
| C>A/G>T | blood | 3 | 5,187,390 | 6 | 8,689,798 | 3 | 7,983,430 | 5.78E-07 | 6.90E-07 | 3.76E-07 | 0.950 | 1.000 | 0.875 |
| C>G/G>C | blood | 5 | 5,187,390 | 7 | 8,689,798 | 7 | 7,983,430 | 9.64E-07 | 8.06E-07 | 8.77E-07 | 1.000 | 1.000 | 1.000 |
| C>T/G>A | blood | 88 | 5,187,390 | 141 | 8,689,798 | 150 | 7,983,430 | 1.70E-05 | 1.62E-05 | 1.88E-05 | 0.849 | 1.000 | 0.561 |
| A>C/T>G | saliva | 5 | 6,171,370 | 7 | 7,331,956 | 4 | 5,931,884 | 8.10E-07 | 9.55E-07 | 6.74E-07 | 1.000 | 1.000 | 0.941 |
| A>G/T>C | saliva | 90 | 6,171,370 | 107 | 7,331,956 | 120 | 5,931,884 | 1.46E-05 | 1.46E-05 | 2.02E-05 | 0.114 | 1.000 | 0.145 |
| A>T/T>A | saliva | 5 | 6,171,370 | 0 | 7,331,956 | 2 | 5,931,884 | 8.10E-07 | 0.00E+00 | 3.37E-07 | 0.849 | 0.179 | 0.561 |
| C>A/G>T | saliva | 4 | 4,929,860 | 1 | 5,856,968 | 6 | 4,738,552 | 8.11E-07 | 1.71E-07 | 1.27E-06 | 0.849 | 0.726 | 0.302 |
| C>G/G>C | saliva | 9 | 4,929,860 | 7 | 5,856,968 | 7 | 4,738,552 | 1.83E-06 | 1.20E-06 | 1.48E-06 | 1.000 | 1.000 | 0.941 |
| C>T/G>A | saliva | 73 | 4,929,860 | 110 | 5,856,968 | 87 | 4,738,552 | 1.48E-05 | 1.88E-05 | 1.84E-05 | 0.654 | 0.717 | 0.941 |
| A>C/T>G | oocyte | 11 | 344,776,941 | 29 | 335,059,336 | 29 | 399,720,556 | 3.19E-08 | 8.66E-08 | 7.26E-08 | 0.114 | 0.071 | 0.875 |
| A>G/T>C | oocyte | 223 | 344,776,941 | 191 | 335,059,336 | 234 | 399,720,556 | 6.47E-07 | 5.70E-07 | 5.85E-07 | 0.823 | 0.726 | 0.941 |
| A>T/T>A | oocyte | 24 | 344,776,941 | 24 | 335,059,336 | 27 | 399,720,556 | 6.96E-08 | 7.16E-08 | 6.75E-08 | 1.000 | 1.000 | 0.941 |
| C>A/G>T | oocyte | 115 | 275,417,298 | 122 | 267,654,608 | 158 | 319,307,768 | 4.18E-07 | 4.56E-07 | 4.95E-07 | 0.654 | 1.000 | 0.875 |
| C>G/G>C | oocyte | 51 | 275,417,298 | 47 | 267,654,608 | 72 | 319,307,768 | 1.85E-07 | 1.76E-07 | 2.25E-07 | 0.823 | 1.000 | 0.561 |
| C>T/G>A | oocyte | 170 | 275,417,298 | 157 | 267,654,608 | 193 | 319,307,768 | 6.17E-07 | 5.87E-07 | 6.04E-07 | 1.000 | 1.000 | 0.941 |

**Table S8. Mutations in the CpG context.** Fisher's exact test (FET) *p*-values were corrected for multiple testing using the method of Benjamini-Hochberg to control the false discovery rate. 'nt_type' is the number of sequenced nucleotides that can effectively lead to the observed mutation type.

| age_group | tissue | mutation type | n_mut (CpG) | nt_type (CpG) | n_mut (non-CpG) | nt_type (non-CpG) | mut_freq (CpG) | mut_freq (non-CpG) | FET p |
|---|---|---|---|---|---|---|---|---|---|
| intermediate | blood | C>T | 0 | 516,097 | 8 | 5,605,026 | 0.00E+00 | 1.43E-06 | 1.000 |
| intermediate | blood | G>A | 29 | 516,097 | 104 | 2,052,578 | 5.62E-05 | 5.07E-05 | 0.797 |
| old | blood | C>T | 0 | 474,145 | 18 | 5,149,410 | 0.00E+00 | 3.50E-06 | 0.797 |
| old | blood | G>A | 29 | 474,145 | 103 | 1,885,730 | 6.12E-05 | 5.46E-05 | 0.797 |
| young | blood | C>T | 0 | 308,085 | 6 | 3,345,930 | 0.00E+00 | 1.79E-06 | 1.000 |
| young | blood | G>A | 24 | 308,085 | 58 | 1,225,290 | 7.79E-05 | 4.73E-05 | 0.235 |
| intermediate | saliva | C>T | 0 | 347,852 | 11 | 3,777,816 | 0.00E+00 | 2.91E-06 | 0.797 |
| intermediate | saliva | G>A | 29 | 347,852 | 70 | 1,383,448 | 8.34E-05 | 5.06E-05 | 0.235 |
| old | saliva | C>T | 0 | 281,428 | 13 | 3,056,424 | 0.00E+00 | 4.25E-06 | 0.797 |
| old | saliva | G>A | 15 | 281,428 | 59 | 1,119,272 | 5.33E-05 | 5.27E-05 | 1.000 |
| young | saliva | C>T | 0 | 292,790 | 14 | 3,179,820 | 0.00E+00 | 4.40E-06 | 0.797 |
| young | saliva | G>A | 18 | 292,790 | 41 | 1,164,460 | 6.15E-05 | 3.52E-05 | 0.235 |
| intermediate | oocyte | C>T | 1 | 15,896,312 | 47 | 172,640,496 | 6.29E-08 | 2.72E-07 | 0.667 |
| intermediate | oocyte | G>A | 17 | 15,896,312 | 92 | 63,221,488 | 1.07E-06 | 1.46E-06 | 0.726 |
| old | oocyte | C>T | 2 | 18,964,052 | 59 | 205,957,416 | 1.05E-07 | 2.86E-07 | 0.726 |
| old | oocyte | G>A | 24 | 18,964,052 | 108 | 75,422,248 | 1.27E-06 | 1.43E-06 | 0.797 |
| young | oocyte | C>T | 0 | 16,357,347 | 58 | 177,647,526 | 0.00E+00 | 3.26E-07 | 0.235 |
| young | oocyte | G>A | 19 | 16,357,347 | 93 | 65,055,078 | 1.16E-06 | 1.43E-06 | 0.797 |
| intermediate | blood | C>T/G>A | 29 | 1,032,194 | 112 | 7,657,604 | 2.81E-05 | 1.46E-05 | **8.38E-03** |
| old | blood | C>T/G>A | 29 | 948,290 | 121 | 7,035,140 | 3.06E-05 | 1.72E-05 | **1.38E-02** |
| young | blood | C>T/G>A | 24 | 616,170 | 64 | 4,571,220 | 3.90E-05 | 1.40E-05 | **3.70E-04** |
| intermediate | saliva | C>T/G>A | 29 | 695,704 | 81 | 5,161,264 | 4.17E-05 | 1.57E-05 | **3.57E-04** |
| old | saliva | C>T/G>A | 15 | 562,856 | 72 | 4,175,696 | 2.66E-05 | 1.72E-05 | 0.201 |

| | | | | | | | | | |
|---|---|---|---|---|---|---|---|---|---|
| young | saliva | C>T/G>A | 18 | 585,580 | 55 | 4,344,280 | 3.07E-05 | 1.27E-05 | **8.38E-03** |
| intermediate | oocyte | C>T/G>A | 18 | 31,792,624 | 139 | 235,861,984 | 5.66E-07 | 5.89E-07 | 1.000 |
| old | oocyte | C>T/G>A | 26 | 37,928,104 | 167 | 281,379,664 | 6.86E-07 | 5.94E-07 | 0.647 |
| young | oocyte | C>T/G>A | 19 | 32,714,694 | 151 | 242,702,604 | 5.81E-07 | 6.22E-07 | 1.000 |

**Table S9. Analysis of mutation strand bias.** *p*-values for differences in mutation frequencies of the two complementary mutation types (strand bias) were calculated with Fisher's exact test (FET) and corrected for multiple testing using the method of Benjamini-Hochberg to control the false discovery rate. 'nt (type)' is the number of sequenced nucleotides that can effectively lead to the observed mutation type.

| type 1 | type 2 | tissue | age group | n_mut (type 1) | nt (type 1) | n_mut (type 2) | nt (type 2) | mut_freq (type 1) | mut_freq (type 2) | type 2 / type 1 | FET p |
|---|---|---|---|---|---|---|---|---|---|---|---|
| A>C | T>G | blood | intermediate | 3 | 6,044,358 | 1 | 4,833,833 | 4.96E-07 | 2.07E-07 | 0.42 | 1.000 |
| A>C | T>G | blood | old | 6 | 5,553,030 | 0 | 4,440,905 | 1.08E-06 | 0.00E+00 | 0.00 | 0.334 |
| A>C | T>G | blood | young | 2 | 3,608,190 | 0 | 2,885,565 | 5.54E-07 | 0.00E+00 | 0.00 | 1.000 |
| A>C | T>G | saliva | intermediate | 3 | 4,073,928 | 4 | 3,258,028 | 7.36E-07 | 1.23E-06 | 1.67 | 1.000 |
| A>C | T>G | saliva | old | 2 | 3,295,992 | 2 | 2,635,892 | 6.07E-07 | 7.59E-07 | 1.25 | 1.000 |
| A>C | T>G | saliva | young | 3 | 3,429,060 | 2 | 2,742,310 | 8.75E-07 | 7.29E-07 | 0.83 | 1.000 |
| A>C | T>G | oocyte | intermediate | 21 | 186,172,368 | 8 | 148,886,968 | 1.13E-07 | 5.37E-08 | 0.48 | 0.410 |
| A>C | T>G | oocyte | old | 17 | 222,100,728 | 12 | 177,619,828 | 7.65E-08 | 6.76E-08 | 0.88 | 1.000 |
| A>C | T>G | oocyte | young | 6 | 191,571,858 | 5 | 153,205,083 | 3.13E-08 | 3.26E-08 | 1.04 | 1.000 |
| A>G | T>C | blood | intermediate | 72 | 6,044,358 | 124 | 4,833,833 | 1.19E-05 | 2.57E-05 | 2.15 | **2.47E-07** |
| A>G | T>C | blood | old | 79 | 5,553,030 | 189 | 4,440,905 | 1.42E-05 | 4.26E-05 | 2.99 | **6.48E-17** |
| A>G | T>C | blood | young | 38 | 3,608,190 | 71 | 2,885,565 | 1.05E-05 | 2.46E-05 | 2.34 | **1.96E-05** |
| A>G | T>C | saliva | intermediate | 30 | 4,073,928 | 77 | 3,258,028 | 7.36E-06 | 2.36E-05 | 3.21 | **3.83E-08** |
| A>G | T>C | saliva | old | 38 | 3,295,992 | 82 | 2,635,892 | 1.15E-05 | 3.11E-05 | 2.70 | **2.47E-07** |
| A>G | T>C | saliva | young | 31 | 3,429,060 | 59 | 2,742,310 | 9.04E-06 | 2.15E-05 | 2.38 | **7.01E-05** |
| A>G | T>C | oocyte | intermediate | 61 | 186,172,368 | 130 | 148,886,968 | 3.28E-07 | 8.73E-07 | 2.66 | **2.98E-10** |
| A>G | T>C | oocyte | old | 91 | 222,100,728 | 143 | 177,619,828 | 4.10E-07 | 8.05E-07 | 1.96 | **4.27E-07** |
| A>G | T>C | oocyte | young | 82 | 191,571,858 | 141 | 153,205,083 | 4.28E-07 | 9.20E-07 | 2.15 | **5.12E-08** |
| A>T | T>A | blood | intermediate | 4 | 6,044,358 | 2 | 4,833,833 | 6.62E-07 | 4.14E-07 | 0.63 | 1.000 |
| A>T | T>A | blood | old | 1 | 5,553,030 | 0 | 4,440,905 | 1.80E-07 | 0.00E+00 | 0.00 | 1.000 |
| A>T | T>A | blood | young | 1 | 3,608,190 | 1 | 2,885,565 | 2.77E-07 | 3.47E-07 | 1.25 | 1.000 |
| A>T | T>A | saliva | intermediate | 0 | 4,073,928 | 0 | 3,258,028 | 0.00E+00 | 0.00E+00 | NA | 1.000 |
| A>T | T>A | saliva | old | 1 | 3,295,992 | 1 | 2,635,892 | 3.03E-07 | 3.79E-07 | 1.25 | 1.000 |
| A>T | T>A | saliva | young | 3 | 3,429,060 | 2 | 2,742,310 | 8.75E-07 | 7.29E-07 | 0.83 | 1.000 |
| A>T | T>A | oocyte | intermediate | 16 | 186,172,368 | 8 | 148,886,968 | 8.59E-08 | 5.37E-08 | 0.63 | 1.000 |
| A>T | T>A | oocyte | old | 11 | 222,100,728 | 16 | 177,619,828 | 4.95E-08 | 9.01E-08 | 1.82 | 1.000 |

| | | | | | | | | | | | |
|---|---|---|---|---|---|---|---|---|---|---|---|
| A>T | T>A | oocyte | young | 13 | 191,571,858 | 11 | 153,205,083 | 6.79E-08 | 7.18E-08 | 1.06 | 1.000 |
| C>A | G>T | blood | intermediate | 4 | 6,121,123 | 2 | 2,568,675 | 6.53E-07 | 7.79E-07 | 1.19 | 1.000 |
| C>A | G>T | blood | old | 2 | 5,623,555 | 1 | 2,359,875 | 3.56E-07 | 4.24E-07 | 1.19 | 1.000 |
| C>A | G>T | blood | young | 2 | 3,654,015 | 1 | 1,533,375 | 5.47E-07 | 6.52E-07 | 1.19 | 1.000 |
| C>A | G>T | saliva | intermediate | 0 | 4,125,668 | 1 | 1,731,300 | 0.00E+00 | 5.78E-07 | NA | 0.880 |
| C>A | G>T | saliva | old | 3 | 3,337,852 | 3 | 1,400,700 | 8.99E-07 | 2.14E-06 | 2.38 | 0.880 |
| C>A | G>T | saliva | young | 2 | 3,472,610 | 2 | 1,457,250 | 5.76E-07 | 1.37E-06 | 2.38 | 0.880 |
| C>A | G>T | oocyte | intermediate | 82 | 188,536,808 | 40 | 79,117,800 | 4.35E-07 | 5.06E-07 | 1.16 | 0.880 |
| C>A | G>T | oocyte | old | 105 | 224,921,468 | 53 | 94,386,300 | 4.67E-07 | 5.62E-07 | 1.20 | 0.880 |
| C>A | G>T | oocyte | young | 78 | 194,004,873 | 37 | 81,412,425 | 4.02E-07 | 4.54E-07 | 1.13 | 0.880 |
| C>G | G>C | blood | intermediate | 1 | 6,121,123 | 6 | 2,568,675 | 1.63E-07 | 2.34E-06 | 14.30 | **7.04E-03** |
| C>G | G>C | blood | old | 2 | 5,623,555 | 5 | 2,359,875 | 3.56E-07 | 2.12E-06 | 5.96 | **3.75E-02** |
| C>G | G>C | blood | young | 1 | 3,654,015 | 4 | 1,533,375 | 2.74E-07 | 2.61E-06 | 9.53 | **3.75E-02** |
| C>G | G>C | saliva | intermediate | 0 | 4,125,668 | 7 | 1,731,300 | 0.00E+00 | 4.04E-06 | NA | **1.77E-03** |
| C>G | G>C | saliva | old | 1 | 3,337,852 | 6 | 1,400,700 | 3.00E-07 | 4.28E-06 | 14.30 | **7.04E-03** |
| C>G | G>C | saliva | young | 2 | 3,472,610 | 7 | 1,457,250 | 5.76E-07 | 4.80E-06 | 8.34 | **7.04E-03** |
| C>G | G>C | oocyte | intermediate | 32 | 188,536,808 | 15 | 79,117,800 | 1.70E-07 | 1.90E-07 | 1.12 | 0.750 |
| C>G | G>C | oocyte | old | 47 | 224,921,468 | 25 | 94,386,300 | 2.09E-07 | 2.65E-07 | 1.27 | 0.412 |
| C>G | G>C | oocyte | young | 26 | 194,004,873 | 25 | 81,412,425 | 1.34E-07 | 3.07E-07 | 2.29 | **7.04E-03** |
| C>T | G>A | blood | intermediate | 8 | 6,121,123 | 133 | 2,568,675 | 1.31E-06 | 5.18E-05 | 39.62 | **7.12E-59** |
| C>T | G>A | blood | old | 18 | 5,623,555 | 132 | 2,359,875 | 3.20E-06 | 5.59E-05 | 17.48 | **9.43E-50** |
| C>T | G>A | blood | young | 6 | 3,654,015 | 82 | 1,533,375 | 1.64E-06 | 5.35E-05 | 32.57 | **6.09E-36** |
| C>T | G>A | saliva | intermediate | 11 | 4,125,668 | 99 | 1,731,300 | 2.67E-06 | 5.72E-05 | 21.45 | **1.13E-39** |
| C>T | G>A | saliva | old | 13 | 3,337,852 | 74 | 1,400,700 | 3.89E-06 | 5.28E-05 | 13.56 | **1.18E-26** |
| C>T | G>A | saliva | young | 14 | 3,472,610 | 59 | 1,457,250 | 4.03E-06 | 4.05E-05 | 10.04 | **1.79E-19** |
| C>T | G>A | oocyte | intermediate | 48 | 188,536,808 | 109 | 79,117,800 | 2.55E-07 | 1.38E-06 | 5.41 | **1.03E-24** |
| C>T | G>A | oocyte | old | 61 | 224,921,468 | 132 | 94,386,300 | 2.71E-07 | 1.40E-06 | 5.16 | **1.98E-28** |
| C>T | G>A | oocyte | young | 58 | 194,004,873 | 112 | 81,412,425 | 2.99E-07 | 1.38E-06 | 4.60 | **2.14E-22** |

**Table S10. Observed versus expected numbers of de novo mutations. (A)** Somatic and germline mutations in young women. **(B)** Somatic and germline mutations in women belonging to the intermediate age group. **(C)** Somatic and germline mutations in old women. **(D)** Variant hotspots. *p*-values were calculated with a two-sided binomial test. The observed numbers of variants and the numbers expected under random and neutral expectations (based on the frequency at specific sites) are shown. Blue indicates higher observed numbers than expected, red indicates lower observed numbers than expected.

| **A) young** | | ***de novo mutations (blood)*** | | | ***de novo mutations (saliva)*** | | | ***de novo mutations (oocytes)*** | | |
|---|---|---|---|---|---|---|---|---|---|---|
| **region** | **nt** | **observed** | **expected** | ***p-value*** | **observed** | **expected** | ***p-value*** | **observed** | **expected** | ***p-value*** |
| **all** | 16569 | 209 | | | 186 | | | 594 | | |
| **D-loop** | 1122 | 11 | 14 | 0.490 | 31 | 13 | **3.20E-06** | 215 | 40 | **3.16E-96** |
| **intergenic** | 88 | 0 | 1 | 0.633 | 0 | 1 | 1.000 | 3 | 3 | 1.000 |
| **non-coding** | 1210 | 11 | 15 | 0.350 | 31 | 14 | **1.51E-05** | 218 | 43 | **1.06E-92** |
| **coding** | 15359 | 198 | 194 | 0.350 | 155 | 172 | **1.51E-05** | 376 | 551 | **1.06E-92** |
| **tRNA** | 1505 | 20 | 19 | 0.809 | 9 | 17 | **4.10E-02** | 36 | 54 | **8.21E-03** |
| **rRNA** | 2513 | 47 | 32 | **4.95E-03** | 23 | 28 | 0.357 | 67 | 90 | **7.13E-03** |
| **protein coding** | 11341 | 131 | 143 | 0.074 | 123 | 127 | 0.528 | 273 | 407 | **1.20E-29** |

| **B) intermediate** | | ***de novo mutations (blood)*** | | | ***de novo mutations (saliva)*** | | | ***de novo mutations (oocytes)*** | | |
|---|---|---|---|---|---|---|---|---|---|---|
| **region** | **nt** | **observed** | **expected** | ***p-value*** | **observed** | **expected** | ***p-value*** | **observed** | **expected** | ***p-value*** |
| **all** | 16569 | 360 | | | 232 | | | 570 | | |
| **D-loop** | 1122 | 21 | 24 | 0.530 | 19 | 16 | 0.361 | 213 | 39 | **1.83E-98** |
| **intergenic** | 88 | 3 | 2 | 0.446 | 2 | 1 | 0.349 | 2 | 3 | 0.775 |
| **non-coding** | 1210 | 24 | 26 | 0.761 | 21 | 17 | 0.311 | 215 | 42 | **3.98E-94** |
| **coding** | 15359 | 336 | 334 | 0.761 | 211 | 215 | 0.311 | 355 | 528 | **3.98E-94** |
| **tRNA** | 1505 | 31 | 33 | 0.854 | 16 | 21 | 0.303 | 36 | 52 | **1.95E-02** |
| **rRNA** | 2513 | 87 | 55 | **8.80E-06** | 45 | 35 | 0.081 | 65 | 86 | **1.19E-02** |
| **protein coding** | 11341 | 218 | 246 | **1.77E-03** | 150 | 159 | 0.230 | 254 | 390 | **6.63E-32** |

| **C) old** | | ***de novo mutations (blood)*** | | | ***de novo mutations (saliva)*** | | | ***de novo mutations (oocytes)*** | | |
|---|---|---|---|---|---|---|---|---|---|---|
| **region** | **nt** | **observed** | **expected** | ***p-value*** | **observed** | **expected** | ***p-value*** | **observed** | **expected** | ***p-value*** |
| **all** | 16569 | 435 | | | 226 | | | 713 | | |
| **D-loop** | 1122 | 22 | 29 | 0.181 | 21 | 15 | 0.143 | 238 | 48 | **1.02E-97** |

| | | | | | | | | | | |
|---|---|---|---|---|---|---|---|---|---|---|
| **intergenic** | 88 | 1 | 2 | 0.735 | 0 | 1 | 0.638 | 2 | 4 | 0.600 |
| **non-coding** | 1210 | 23 | 32 | 0.117 | 21 | 17 | 0.249 | 240 | 52 | **1.06E-92** |
| **coding** | 15359 | 412 | 403 | 0.117 | 205 | 209 | 0.249 | 473 | 661 | **1.06E-92** |
| **tRNA** | 1505 | 37 | 40 | 0.739 | 26 | 21 | 0.203 | 54 | 65 | 0.171 |
| **rRNA** | 2513 | 81 | 66 | 0.052 | 41 | 34 | 0.227 | 99 | 108 | 0.375 |
| **protein coding** | 11341 | 294 | 298 | 0.718 | 138 | 155 | **1.82E-02** | 320 | 488 | **1.66E-38** |

| **D) variant hotspots (sites)** | | | | |
|---|---|---|---|---|
| **region** | **nt** | **observed** | **expected** | ***p-value*** |
| **all** | 16569 | 58 | | |
| **D-loop** | 1122 | 40 | 4 | **2.22E-33** |
| **intergenic** | 88 | 0 | 0 | 1.000 |
| **non-coding** | 1210 | 40 | 4 | **4.12E-32** |
| **coding** | 15359 | 18 | 54 | **4.12E-32** |
| **tRNA** | 1505 | 4 | 5 | 0.818 |
| **rRNA** | 2513 | 4 | 9 | 0.097 |
| **protein coding** | 11341 | 10 | 40 | **1.17E-15** |

**Table S11. Nonsynonymous-to-synonymous rate ratios (hN/hS) for variants found in a tissue.** NE represents the neutral expectations (bootstrapped for 100 replicates). Values outside of neutral expectations are shown in bold.

| age_group | tissue | variants | hN/hS | median (NE) | *lower (NE)* | *upper (NE)* | *n sites* |
|---|---|---|---|---|---|---|---|
| young | blood | all | **1.64** | 1.07 | *0.78* | *1.57* | *131* |
| intermediate | blood | all | 1.50 | 1.09 | *0.85* | *1.50* | *218* |
| old | blood | all | 1.25 | 1.05 | *0.78* | *1.31* | *294* |
| young | saliva | all | **2.01** | 1.09 | *0.76* | *1.69* | *123* |
| intermediate | saliva | all | **2.54** | 1.17 | *0.83* | *1.57* | *150* |
| old | saliva | all | 1.45 | 1.17 | *0.87* | *1.59* | *138* |
| young | oocyte | all | 0.9 | 0.97 | *0.79* | *1.24* | *273* |
| intermediate | oocyte | all | 1.19 | 0.97 | *0.78* | *1.25* | *254* |
| old | oocyte | all | 0.99 | 0.99 | *0.78* | *1.19* | *320* |
| young | somatic | non-hotspot | **1.72** | 1.06 | *0.85* | *1.33* | *244* |
| intermediate | somatic | non-hotspot | **1.77** | 1.04 | *0.81* | *1.3* | *360* |
| old | somatic | non-hotspot | **1.26** | 1.01 | *0.83* | *1.25* | *420* |
| young | oocyte* | non-hotspot | 0.9 | 1.01 | *0.77* | *1.25* | *273* |
| intermediate | oocyte* | non-hotspot | 1.19 | 0.95 | *0.76* | *1.26* | *254* |
| old | oocyte* | non-hotspot | 0.99 | 0.99 | *0.82* | *1.26* | *320* |

*None of the oocyte hotspots were located in protein-coding regions, hence results are equal with the analysis of all mutations.

**Table S12. Analysis of potential variant hotspots.** Variants found in a tissue type in two or more women for somatic tissues or three or more women for oocytes are listed. The hotspot probability represents the probability of a variant to be present by random chance exactly in those individual samples in which we observe it (considering the mtDNA sequencing depth of the individual samples). n_hs refers to the number of women with a specific mutation.

| hotspot type | mutation | region | n_hs | prob |
|---|---|---|---|---|
| oocyte hotspot | 72 T > C | D-loop | 9 | 6.38E-16 |
| oocyte hotspot | 10 T > C | D-loop | 9 | 8.65E-16 |
| oocyte hotspot | 3242 G > A | tRNA | 6 | 4.28E-15 |
| oocyte hotspot | 103 G > A | D-loop | 8 | 1.37E-13 |
| oocyte hotspot | 16468 T > C | D-loop | 8 | 2.06E-13 |
| oocyte hotspot | 16310 G > A | D-loop | 8 | 3.70E-13 |
| oocyte hotspot | 318 T > C | D-loop | 7 | 1.22E-12 |
| oocyte hotspot | 16103 A > G | D-loop | 7 | 4.85E-12 |
| oocyte hotspot | 206 T > C | D-loop | 7 | 1.79E-11 |
| oocyte hotspot | 115 T > C | D-loop | 6 | 5.20E-10 |
| oocyte hotspot | 16129 G > A | D-loop | 6 | 1.15E-09 |
| oocyte hotspot | 16157 T > C | D-loop | 5 | 4.11E-09 |
| oocyte hotspot | 146 T > C | D-loop | 5 | 5.69E-09 |
| oocyte hotspot | 16092 T > C | D-loop | 5 | 6.53E-09 |
| oocyte hotspot | 16362 T > C | D-loop | 5 | 9.40E-09 |
| oocyte hotspot | 131 T > C | D-loop | 5 | 1.26E-08 |
| oocyte hotspot | 16437 T > C | D-loop | 5 | 1.33E-08 |
| oocyte hotspot | 139 T > C | D-loop | 5 | 1.59E-08 |
| oocyte hotspot | 16477 G > A | D-loop | 5 | 2.34E-08 |
| oocyte hotspot | 366 G > A | D-loop | 4 | 3.09E-08 |
| oocyte hotspot | 60 T > C | D-loop | 4 | 9.89E-08 |
| oocyte hotspot | 227 A > G | D-loop | 4 | 1.37E-07 |
| oocyte hotspot | 16224 T > C | D-loop | 4 | 1.39E-07 |
| oocyte hotspot | 191 A > G | D-loop | 4 | 1.45E-07 |

| | | | | |
|---|---|---|---|---|
| oocyte hotspot | 193 A > G | D-loop | 4 | 1.63E-07 |
| oocyte hotspot | 153 A > G | D-loop | 4 | 1.75E-07 |
| oocyte hotspot | 16516 G > A | D-loop | 4 | 1.77E-07 |
| oocyte hotspot | 16381 T > C | D-loop | 4 | 1.80E-07 |
| oocyte hotspot | 326 A > C | D-loop | 4 | 2.62E-07 |
| oocyte hotspot | 16156 G > A | D-loop | 4 | 2.73E-07 |
| oocyte hotspot | 16474 G > A | D-loop | 4 | 3.51E-07 |
| oocyte hotspot | 226 T > C | D-loop | 4 | 3.96E-07 |
| oocyte hotspot | 188 A > G | D-loop | 4 | 4.19E-07 |
| oocyte hotspot | 16319 G > A | D-loop | 4 | 6.55E-07 |
| oocyte hotspot | 203 G > A | D-loop | 4 | 9.31E-07 |
| oocyte hotspot | 16324 T > C | D-loop | 4 | 9.56E-07 |
| somatic hotspot | 3244 G > A | tRNA | 5 | 1.63E-12 |
| somatic hotspot | 15924 A > G | tRNA | 3 | 8.73E-11 |
| somatic hotspot | 15173 G > A | protein coding | 3 | 9.04E-11 |
| somatic hotspot | 13063 G > A | protein coding | 3 | 1.51E-10 |
| somatic hotspot | 3022 G > A | rRNA | 3 | 2.81E-10 |
| somatic hotspot | 3243 A > G | tRNA | 3 | 3.46E-10 |
| somatic hotspot | 309 C > T | D-loop | 3 | 7.20E-10 |
| somatic hotspot | 16086 T > C | D-loop | 4 | 7.34E-10 |
| somatic hotspot | 215 A > G | D-loop | 3 | 1.30E-09 |
| somatic hotspot | 2623 A > G | rRNA | 4 | 2.37E-09 |
| somatic hotspot | 2371 T > C | rRNA | 3 | 1.23E-07 |
| somatic hotspot | 533 A > G | D-loop | 3 | 1.44E-07 |
| somatic hotspot | 13676 A > G | protein coding | 3 | 1.79E-07 |
| somatic hotspot | 204 T > C | D-loop | 3 | 2.17E-07 |
| somatic hotspot | 14858 G > A | protein coding | 3 | 3.32E-07 |
| somatic hotspot | 6933 T > C | protein coding | 3 | 4.06E-07 |
| somatic hotspot | 7994 G > A | protein coding | 3 | 7.26E-07 |

| | | | | |
|---|---|---|---|---|
| somatic hotspot | 10974 T > C | protein coding | 3 | 8.79E-07 |
| somatic hotspot | 9128 T > C | protein coding | 3 | 9.24E-07 |
| somatic hotspot | 5262 G > A | protein coding | 3 | 1.04E-06 |
| somatic hotspot | 1071 T > C | rRNA | 3 | 1.32E-06 |
| somatic hotspot | 9445 G > A | protein coding | 3 | 1.84E-06 |

**Table S13. Inheritable heteroplasmies.**

| donor ID | age (years) | age group | position | variant type | localization | hotspot colocalization | MAF blood | MAF saliva | MAF mean (somatic tissues) | MAF_Oo1 | MAF_Oo2 | MAF_Oo3 | MAF_Oo4 | MAF_Oo5 | MAF_Oo6 | MAF mean (oocytes) |
|---|---|---|---|---|---|---|---|---|---|---|---|---|---|---|---|---|
| hs002 | 34 | intermediate | 234 | A > G | D-loop | - | 1.28% | 2.19% | 1.74% | 0.00% | NA | 0.00% | 0.00% | 0.82% | NA | 0.21% |
| hs002 | 34 | intermediate | 941 | G > A | rRNA | - | 3.35% | 1.79% | 2.57% | 0.00% | NA | 5.53% | 0.00% | 0.00% | NA | 1.38% |
| hs004 | 39 | old | 204 | T > C | D-loop | somatic | 0.60% | 0.15% | 0.37% | 4.12% | 0.42% | NA | NA | NA | NA | 2.27% |
| hs004 | 39 | old | 16519 | T > C | D-loop | - | 0.55% | 1.16% | 0.85% | 0.03% | 0.03% | NA | NA | NA | NA | 0.03% |
| hs005 | 24 | young | 2551 | G > A | rRNA | - | NA | 0.48% | 0.48% | 0.00% | 14.02% | 0.00% | 0.00% | NA | NA | 3.51% |
| hs006 | 25 | young | 195 | T > C | D-loop | - | 0.36% | 2.69% | 1.52% | 1.88% | 0.05% | 0.28% | NA | 0.02% | NA | 0.56% |
| hs006 | 25 | young | 3010 | G > A | rRNA | - | 3.90% | 5.27% | 4.59% | 1.77% | 0.00% | 0.00% | NA | 10.34% | NA | 3.03% |
| hs007 | 36 | old | 15773 | G > A | protein coding | - | 33.80% | NA | 33.80% | 81.89% | 19.94% | 25.75% | 0.05% | NA | NA | 31.91% |
| hs010 | 26 | young | 16519 | T > C | D-loop | - | 0.73% | 0.64% | 0.69% | 13.43% | 0.07% | NA | NA | 5.44% | 11.36% | 7.57% |
| hs011 | 38 | old | 143 | G > A | D-loop | - | 0.05% | 0.09% | 0.07% | 1.13% | 0.00% | 0.38% | NA | NA | NA | 0.50% |
| hs011 | 38 | old | 204 | T > C | D-loop | somatic | 2.00% | 2.21% | 2.11% | 0.00% | 0.00% | 0.03% | NA | NA | NA | 0.01% |
| hs011 | 38 | old | 15152 | G > A | protein coding | - | 2.39% | 2.02% | 2.20% | 0.00% | 0.00% | 0.02% | NA | NA | NA | 0.01% |
| hs012 | 42 | old | 143 | G > A | D-loop | - | NA | 0.03% | 0.03% | 1.28% | 0.26% | 0.06% | NA | 3.21% | NA | 1.21% |
| hs012 | 42 | old | 3644 | T > C | protein coding | - | NA | 2.60% | 2.60% | 12.17% | 0.00% | 0.00% | NA | 0.00% | NA | 3.04% |
| hs014 | 34 | intermediate | 16286 | T > C | D-loop | - | 0.82% | NA | 0.82% | 0.02% | 2.71% | 0.03% | 0.71% | 0.00% | NA | 0.69% |
| hs015 | 42 | old | 9110 | T > C | protein coding | - | 5.71% | NA | 5.71% | 3.67% | 80.61% | 0.00% | 0.00% | NA | NA | 21.07% |
| hs016 | 33 | intermediate | 16093 | C > T | D-loop | - | 1.66% | NA | 1.66% | 0.00% | 0.05% | 0.82% | 0.00% | NA | NA | 0.22% |
| hs017 | 20 | young | 16344 | C > T | D-loop | - | 1.13% | NA | 1.13% | 0.00% | 11.93% | 0.03% | 5.63% | 0.00% | NA | 3.52% |
| hs017 | 20 | young | 16519 | T > C | D-loop | - | 1.26% | NA | 1.26% | 0.82% | 0.28% | 0.45% | 0.25% | 4.31% | NA | 1.22% |
| hs018 | 36 | old | 13443 | T > C | protein coding | - | 5.24% | NA | 5.24% | 16.77% | NA | 32.55% | NA | NA | NA | 24.66% |
| hs019 | 35 | old | 146 | T > C | D-loop | oocyte | 0.13% | NA | 0.13% | 2.84% | 0.02% | 0.02% | 0.00% | NA | NA | 0.72% |
| hs020 | 28 | young | 103 | G > A | D-loop | oocyte | 0.16% | NA | 0.16% | 1.56% | 0.00% | 0.07% | NA | NA | NA | 0.55% |
| hs020 | 28 | young | 9975 | T > C | protein coding | - | 0.77% | NA | 0.77% | 0.00% | 22.87% | 0.00% | NA | NA | NA | 7.62% |
| hs021 | 31 | intermediate | 146 | T > C | D-loop | oocyte | 4.26% | NA | 4.26% | 0.00% | 0.32% | 33.95% | 0.16% | NA | NA | 8.61% |
| hs021 | 31 | intermediate | 16168 | T > C | D-loop | - | 7.78% | NA | 7.78% | 0.05% | 1.29% | 0.00% | 42.05% | NA | NA | 10.85% |
| hs023 | 33 | intermediate | 16092 | T > C | D-loop | oocyte | NA | 0.01% | 0.01% | 0.00% | 0.00% | 14.75% | 0.04% | NA | NA | 3.70% |
| hs023 | 33 | intermediate | 16093 | C > T | D-loop | - | NA | 2.49% | 2.49% | 0.03% | 0.00% | 0.04% | 2.10% | NA | NA | 0.54% |
| hs002 | 34 | intermediate | 10197 | G > A | protein coding | - | 0.09% | 0.09% | 0.09% | 0.02% | NA | 0.00% | 0.00% | 0.00% | NA | 0.01% |
| hs002 | 34 | intermediate | 12539 | G > A | protein coding | - | 0.04% | 0.02% | 0.03% | 0.02% | NA | 0.00% | 0.08% | 0.00% | NA | 0.03% |
| hs002 | 34 | intermediate | 16129 | G > A | D-loop | oocyte | 0.29% | 0.34% | 0.32% | 0.02% | NA | 0.00% | 0.00% | 0.01% | NA | 0.01% |
| hs002 | 34 | intermediate | 16519 | C > T | D-loop | - | 0.04% | 0.05% | 0.04% | 0.00% | NA | 0.00% | 0.05% | 0.01% | NA | 0.02% |
| hs003 | 29 | young | 3027 | C > T | rRNA | - | NA | 0.04% | 0.04% | 0.00% | 0.02% | 0.00% | 0.00% | 0.00% | NA | 0.00% |
| hs004 | 39 | old | 16131 | T > C | D-loop | - | 0.23% | 0.11% | 0.17% | 0.18% | 0.00% | NA | NA | NA | NA | 0.09% |
| hs004 | 39 | old | 16207 | G > A | D-loop | - | 0.15% | 0.57% | 0.36% | 0.00% | 0.17% | NA | NA | NA | NA | 0.08% |
| hs006 | 25 | young | 16519 | T > C | D-loop | - | 0.73% | 0.69% | 0.71% | 0.39% | 0.01% | 0.16% | NA | 0.00% | NA | 0.14% |
| hs011 | 38 | old | 313 | C > T | D-loop | - | 0.03% | 0.89% | 0.46% | 0.15% | 0.07% | 0.00% | NA | NA | NA | 0.07% |

| | | | | | | | | | | | | | | | | |
|---|---|---|---|---|---|---|---|---|---|---|---|---|---|---|---|---|
| hs012 | 42 | old | 72 | T > C | D-loop | oocyte | NA | 0.77% | 0.77% | 0.00% | 0.00% | 0.00% | NA | 0.02% | NA | 0.01% |
| hs012 | 42 | old | 73 | A > G | D-loop | - | NA | 0.04% | 0.04% | 0.02% | 0.03% | 0.00% | NA | 0.00% | NA | 0.01% |
| hs012 | 42 | old | 152 | T > C | D-loop | - | NA | 0.81% | 0.81% | 0.00% | 0.00% | 0.00% | NA | 0.17% | NA | 0.04% |
| hs012 | 42 | old | 189 | A > G | D-loop | - | NA | 0.08% | 0.08% | 0.00% | 0.00% | 0.00% | NA | 0.46% | NA | 0.12% |
| hs012 | 42 | old | 204 | T > C | D-loop | somatic | NA | 0.11% | 0.11% | 0.01% | 0.00% | 0.03% | NA | 0.00% | NA | 0.01% |
| hs012 | 42 | old | 13711 | G > A | protein coding | - | NA | 0.09% | 0.09% | 0.02% | 0.00% | 0.00% | NA | 0.00% | NA | 0.01% |
| hs012 | 42 | old | 16519 | T > C | D-loop | - | NA | 0.31% | 0.31% | 0.01% | 0.32% | 0.00% | NA | 0.07% | NA | 0.10% |
| hs014 | 34 | intermediate | 10 | T > C | D-loop | oocyte | 0.03% | NA | 0.03% | 0.00% | 0.03% | 0.00% | 0.04% | 0.00% | NA | 0.02% |
| hs014 | 34 | intermediate | 146 | T > C | D-loop | oocyte | 0.09% | NA | 0.09% | 0.02% | 0.00% | 0.00% | 0.08% | 0.00% | NA | 0.02% |
| hs014 | 34 | intermediate | 16129 | G > A | D-loop | oocyte | 0.34% | NA | 0.34% | 0.02% | 0.00% | 0.03% | 0.18% | 0.00% | NA | 0.05% |
| hs014 | 34 | intermediate | 16519 | T > C | D-loop | - | 0.52% | NA | 0.52% | 0.02% | 0.03% | 0.00% | 0.19% | 0.00% | NA | 0.05% |
| hs015 | 42 | old | 143 | G > A | D-loop | - | 0.03% | NA | 0.03% | 0.00% | 0.28% | 0.06% | 0.00% | NA | NA | 0.08% |
| hs016 | 33 | intermediate | 103 | G > A | D-loop | oocyte | 0.03% | NA | 0.03% | 0.00% | 0.05% | 0.00% | 0.00% | NA | NA | 0.01% |
| hs017 | 20 | young | 103 | G > A | D-loop | oocyte | 0.04% | NA | 0.04% | 0.00% | 0.00% | 0.04% | 0.12% | 0.00% | NA | 0.03% |
| hs017 | 20 | young | 15014 | C > T | protein coding | - | 0.87% | NA | 0.87% | 0.00% | 0.12% | 0.00% | 0.02% | 0.02% | NA | 0.03% |
| hs017 | 20 | young | 15466 | G > A | protein coding | - | 0.02% | NA | 0.02% | 0.00% | 0.00% | 0.00% | 0.05% | 0.03% | NA | 0.02% |
| hs018 | 36 | old | 143 | G > A | D-loop | - | 0.09% | NA | 0.09% | 0.80% | NA | 0.04% | NA | NA | NA | 0.42% |
| hs019 | 35 | old | 73 | G > A | D-loop | - | 0.04% | NA | 0.04% | 0.04% | 0.11% | 0.01% | 0.00% | NA | NA | 0.04% |
| hs019 | 35 | old | 103 | G > A | D-loop | oocyte | 0.02% | NA | 0.02% | 0.93% | 0.54% | 0.03% | 0.33% | NA | NA | 0.46% |
| hs019 | 35 | old | 188 | G > A | D-loop | - | 0.06% | NA | 0.06% | 0.00% | 0.04% | 0.00% | 0.06% | NA | NA | 0.03% |
| hs019 | 35 | old | 3877 | G > A | protein coding | - | 0.70% | NA | 0.70% | 0.00% | 0.02% | 0.00% | 0.00% | NA | NA | 0.01% |
| hs019 | 35 | old | 16126 | C > T | D-loop | - | 0.39% | NA | 0.39% | 0.00% | 0.02% | 0.00% | 0.02% | NA | NA | 0.01% |
| hs022 | 31 | intermediate | 226 | T > C | D-loop | oocyte | 0.03% | NA | 0.03% | 0.11% | 0.00% | 0.14% | 0.03% | NA | NA | 0.07% |
| hs022 | 31 | intermediate | 709 | G > A | rRNA | - | 0.14% | NA | 0.14% | 0.00% | 0.02% | 0.04% | 0.00% | NA | NA | 0.01% |
| hs022 | 31 | intermediate | 15218 | A > G | protein coding | - | 0.12% | NA | 0.12% | 0.02% | 0.00% | 0.03% | 0.03% | NA | NA | 0.02% |
| hs023 | 33 | intermediate | 143 | G > A | D-loop | - | NA | 0.01% | 0.01% | 0.00% | 0.00% | 0.02% | 0.00% | NA | NA | 0.01% |
| hs023 | 33 | intermediate | 16468 | T > C | D-loop | oocyte | NA | 0.01% | 0.01% | 0.00% | 0.56% | 0.19% | 0.00% | NA | NA | 0.19% |

**Table S14. Evaluation of a germline bottleneck.** High-frequency heteroplasmies were measured in at least 3 consensus sequences. High-frequency heteroplasmies were defined as heteroplasmies with a frequency of at least 1% in at least one sample.

| age group | n (heteroplasmies) | bottleneck size (N) | *95% CI (lower bound)* | *95% CI(upper bound)* |
|---|---|---|---|---|
| **all** | **63** | **909.39** | ***0.47*** | ***5,329.73*** |
| young | 13 | 752.84 | *0.47* | *4,649.81* |
| intermediate | 22 | 1,439.77 | *0.02* | *9,692.42* |
| old | 28 | 565.36 | *0.92* | *4,157.39* |
| **High-confidence (all)** | **30** | **1,040.03** | ***0.34*** | ***11,089.05*** |
| High-confidence (young) | 9 | 227.33 | *0.44* | *1,832.95* |
| High-confidence (intermediate) | 10 | 1,318.28 | *1.07* | *4,270.24* |
| High-confidence (old) | 11 | 1,452.01 | *0.34* | *11,089.05* |
| **High-frequency (all)** | **27** | **30.41** | ***0.02*** | ***178.53*** |
| High-frequency (young) | 8 | 25.03 | *0.47* | *98.42* |
| High-frequency (intermediate) | 8 | 38.60 | *0.02* | *74.42* |
| High-frequency (old) | 11 | 28.38 | *0.38* | *178.53* |

**Table S15. The numbers and proportions of mtDNA variants in oocytes depending on their frequency, heritability, and age group.** These are shown separately for *de novo* mutations with MAF <1%, *de novo* mutations with high-confidence MAF ≥1% (measured in ≥3 DCS to ensure proper representation on MAFs; 10 mutations measured in <3 DCS were excluded), and inheritable heteroplasmies. The neutral expectations confidence intervals (CIs) represents the neutral expectations based on bootstrapping for 100 replicates.

| | | Region length | Number (and proportion) of mutations in young women | Number (and proportion) of mutations in intermediate women | Number (and proportion) of mutations in old women |
|---|---|---|---|---|---|
| ***De novo mutations with MAF<1%*** | | | **561** | **560** | **687** |
| | D-loop | 1,122 bp (6.8%) | 198 (35.3%) | 208 (37.1%) | 223 (32.5%) |
| | intergenic | 88 bp (0.5%) | 3 (0.5%) | 2 (0.4%) | 2 (0.3%) |
| | protein-coding | 11,341 bp (68.4%) | 260 (46.3%) | 250 (44.6%) | 313 (45.6%) |
| | *synonymous* | | *77* | *59* | *85* |
| | *non-synonymous* | | *183* | *191* | *228* |
| | *n(nsyn)/n(syn)* | | *2.38* | *3.24* | *2.68* |
| | *hN/hS (neutral expectation CI)* | | *0.79 (0.77-1.20)* | *1.08 (0.77-1.24)* | *0.89 (0.80-1.27)* |
| | rRNA | 2,513 bp (15.2%) | 67 (11.9%) | 64 (11.4%) | 97 (14.1%) |
| | tRNA | 1,505 bp (9.1%) | 33 (5.9%) | 36 (6.4%) | 52 (7.6%) |
| ***De novo mutations with MAF≥1%*** | | | **23** | **10** | **26** |
| | D-loop | 1,122 bp (6.8%) | 13 (56.5%) | 5 (50%) | 15 (57.7%) |
| | intergenic | 88 bp (0.5%) | 0 (0.0%) | 0 (0.0%) | 0 (0.0%) |
| | protein-coding | 11,341 bp (68.4%) | 9 (39.1%) | 4 (40.0%) | 7 (26.9%) |
| | *synonymous* | | *2* | *1* | *2* |
| | *non-synonymous* | | *7* | *3* | *5* |
| | *n(nsyn)/n(syn)* | | *3.50* | *3.00* | *2.50* |
| | *hN/hS (neutral expectation CI)* | | *1.16 (0.46-2.94)* | *1.00 (0.12-1.10)* | *0.83 (0.15-2.21)* |
| | rRNA | 2,513 bp (15.2%) | 0 (0.0%) | 1 (10.0%) | 2 (7.7%) |
| | tRNA | 1,505 bp (9.1%) | 1 (4.3%) | 0 (0.0%) | 2 (7.7%) |
| **Inheritable heteroplasmies** | | | **13** | **22** | **28** |
| | D-loop | 1,122 bp (6.8%) | 7 (53.8%) | 17 (77.3%) | 21 (75.0%) |
| | intergenic | 88 bp (0.5%) | 0 (0.0%) | 0 (0.0%) | 0 (0.0%) |
| | protein-coding | 11,341 bp (68.4%) | 3 (23.1%) | 3 (13.6%) | 7 (25.0%) |

| | | | | | |
|---|---|---|---|---|---|
| | *synonymous* | | *2* | *0* | *2* |
| | *non-synonymous* | | *1* | *3* | *5* |
| | *n(nsyn)/n(syn)* | | *0.50* | *NA* | *2.50* |
| | *hN/hS (neutral expectation CI)* | | *0.17 (0.07-1.13)* | *NA* | *0.83 (0.28-2.21)* |
| | rRNA | 2,513 bp (15.2%) | 3 (23.1%) | 2 (9.1%) | 0 (0.0%) |
| | tRNA | 1,505 bp (9.1%) | 0 (0.0%) | 0 (0.0%) | 0 (0.0%) |
